# A Bias-Accounting Meta-Analytic Approach Refines and Expands the Cerebellar Behavioral Topography

**DOI:** 10.1101/2024.10.31.621398

**Authors:** Neville Magielse, Aikaterina Manoli, Simon B. Eickhoff, Peter T. Fox, Amin Saberi, Sofie L. Valk

## Abstract

The cerebellum plays important roles in motor, cognitive, and emotional behaviors. Previous cerebellar coordinate-based meta-analyses (CBMAs) have complemented precision-mapping and parcellation approaches by finding generalizable cerebellar activations across the largest possible set of behaviors. However, cerebellar CBMAs face challenges due to inherent methodological limitations exacerbated by historical cerebellar neglect in neuroimaging. Here, we show overrepresentation of superior activations, rendering the null hypothesis of standard activation likelihood estimation (ALE) unsuitable. Our new method, cerebellum-specific ALE (C-SALE), finds behavioral convergence beyond baseline activation rates. It does this by testing experimental activations versus null models sampled from a data-driven probability distribution of finding activations at any cerebellar location. Task-specific mappings in the BrainMap meta-analytic database illustrated improved specificity of the new method. Multiple (sub)domains reached convergence in specific cerebellar subregions, supporting dual motor representations and placing cognition in posterior-lateral regions. We show our method and findings were replicable within NeuroSynth. Across both databases, 54/138 task domains or behavioral terms, including sustained attention, somesthesis, inference, anticipation and rhythm, reached convergence in specific cerebellar subgregions. Maps largely corresponded with cerebellar atlases but also showed many complementary mappings. Repeated subsampling showed that motor behaviors, and to a lesser extent language and working memory, mapped to especially consistent cerebellar subregions. Lastly, we found that cerebellar clusters were parts of brain-wide coactivation networks with cortical and subcortical regions implied in these behaviors. Together, our method further complements and expands understanding of cerebellar involvement in human behavior, highlighting regions for future investigation in both basic and clinical applications.

**Highlights:**

● Biases in reported cerebellar activations strongly favors superior regions.
● A new method of meta-analysis increases cerebellar mapping specificity and accuracy.
● Large-scale meta-analyses support cerebellar roles in cognitive, affective, and motor behaviors.
● 54 task domains/ terms converged, including sustained attention, somesthesis, inference, anticipation and rhythm.

Graphical abstract

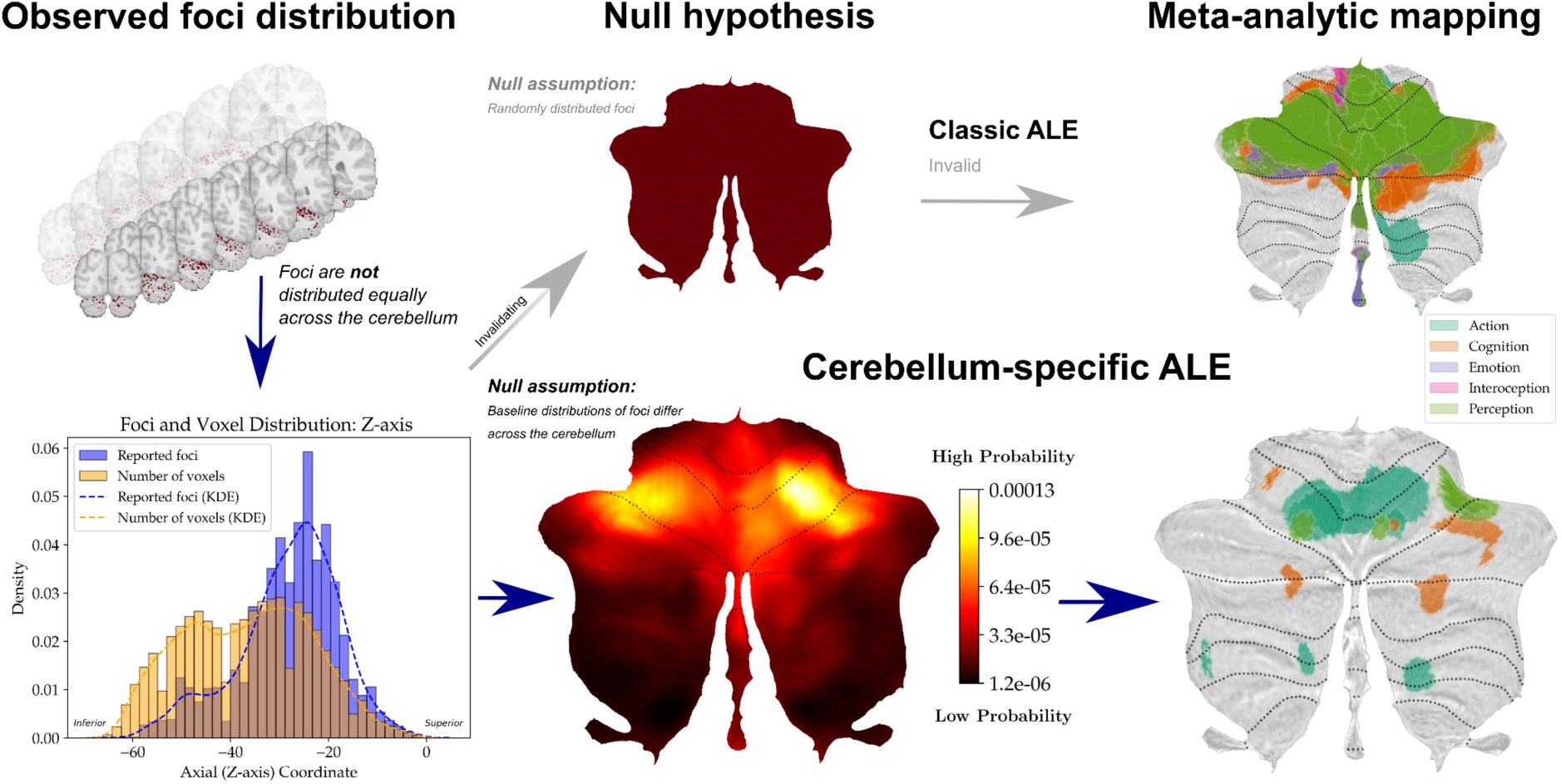

## 1. Introduction

The cerebellum has received considerable attention due to its involvement in modulating activity of widespread regions of the brain (Eccles et al., 1967; Habas, 2021; Ito, 1993; Palesi et al., 2020; Schmahmann, 1996). This perspective is consistent with current neuroscientific paradigms that conceptualize the brain as a complex system of interconnected networks rather than discrete, isolated areas (Srivastava et al., 2022). Although the cerebellum has traditionally been associated with motor control, consensus is now that it plays integrative roles in brain-wide networks spanning behavioral domains (BDs) including emotion, language, and social cognition across the lifespan (Adamaszek et al., 2017; Arleo et al., 2023; Caligiore et al., 2017; Koziol et al., 2014; Leto et al., 2015; Mariën et al., 2014; Van Overwalle et al., 2020b).

The cerebellar circuit is evolutionarily organized into relatively separate functional modules that are connected to diverse extra-cerebellar areas in reciprocal loops (Bolk, 1906; Bostan and Strick, 2018; Chauvel et al., 2023; Glickstein and Voogd, 2009; Kelly and Strick, 2003; Larsell and Jansen, 1970; Magielse et al., 2023, 2022; Nieuwenhuys, 1967; Palesi et al., 2017, 2015; Ramnani, 2006). This organization likely facilitates cerebellar involvement in various human brain networks and with that, behaviours. Decisively mapping cerebellar functions to its subregions is, however, challenging. First, the small cerebellum (10% of brain volume) and its many folds (Azevedo et al., 2009; Heuer et al., 2022; Magielse et al., 2023) exaggerate partial volume effects at relatively crude neuroimaging resolutions. These spatial inaccuracies are magnified in the case of cerebellar spatial misalignment (Diedrichsen et al., 2009), compounding substantial interindividual variability in cerebellar behavioral mapping (Marek et al., 2018; Xue et al., 2020). Secondly, the signal-to-noise ratio (SNR) in the cerebellum is generally low, with common physiological noise and artifacts due to the location in the head and proximity to blood vessels (Schlerf et al., 2014). Together, such challenges can make it difficult to reliably obtain functional signals in individual brain imaging studies.

Despite these challenges, several strategies have successfully mapped cerebellar functions using functional magnetic resonance imaging (fMRI). First, high-resolution mapping in a few individuals has revealed accurate, individualized resting-state (Marek et al., 2018; Xue et al., 2020) and task-based (Boillat et al., 2020; Saadon-Grosman et al., 2024) topographies. Large, high-quality datasets have been used to establish cerebellar subregion-specific activity in behaviors, replicating the established double motor representation and revealing three separate working memory, social cognition, and language mappings (Guell et al., 2018a). Second, a rich task-based framework across multiple individuals divided the cerebellum into non-overlapping subregions (parcels) corresponding to different functional communities. Through this approach, it was revealed that cerebellar functional borders readily cross lobular boundaries (King et al., 2019). Recently, a functional fusion framework expanded on this parcellation by integrating large-scale task and resting-state fMRI data to improve the discriminability of cerebellar functional borders (Nettekoven et al., 2024). Here, the fusion (Zhi et al., 2023) improved inferences made solely from either task-based or resting-state data (Buckner et al., 2011; Xue et al., 2020). Together, these and other parcellations and mapping studies have revealed much of the known cerebellar functional topography. They paint a relatively coherent picture: motor representations are consistently located in both anterior and posterior-inferior regions, whereas cognitive functions tend to occupy posterior-lateral Crura I-II and inferior lobules IX-X. However, these efforts differ in many respects, such as the number and specific set of included behaviours and their mapping to the cerebellum. For instance, a comparison of the social cognition domain in state-of-the-art mappings (Saadon-Grosman et al., 2024) and parcellations (Nettekoven et al., 2024) shows different localisations. This is not surprising, as these two frameworks have different goals: whereas mappings maximize for correspondence with behaviors, parcellations maximize for border discriminability across functional communities (King et al., 2019; Nettekoven et al., 2024; Zhi et al., 2022).

Importantly, individual neuroimaging studies aiming to map behaviors have shortcomings, such as often limited sample sizes leading to lower power, methodological flexibility (Carp, 2012), and site– or population-specific idiosyncrasies (Button et al., 2013). The smaller the study, the more susceptible it is to observe random effects. From their outset, coordinate-based meta-analyses (CBMAs) were developed to address these shortcomings, by identifying the most consistent findings across as many individual studies as possible.

Specifically, activation likelihood estimation (ALE) (Eickhoff et al., 2012, 2009; Laird et al., 2005a) identifies brain regions that are reported to be involved in a task or domain above chance-level. In other words, it finds regions where activity converges in space despite imaging inaccuracies, noise, and low statistical power of individual studies (Button et al., 2013). This convergence implies highly consistent activity in that region. CBMAs, both at the whole-brain-level and using the cerebellum as region-of-interest (ROI), have provided important complementary understanding of brain and cerebellar functions. They were among the first to help establish cerebellar involvement in specific cognitive and behavioral processes (Stoodley and Schmahmann, 2009). Studies at the whole-brain-level have provided insight on whether the cerebellum and its subregions are activated above chance in these processes. In this way, cerebellar involvement in e.g., audition (Petacchi et al., 2005), verbal working memory (Emch et al., 2019), and cognitive control (Niendam et al., 2012; Radua et al., 2014) has been established. CBMAs within the cerebellar ROI have enabled localization of behaviors to cerebellar subregions, including motor (learning) behavior, emotion, and several aspects of social cognition and executive function (Baumann and Mattingley, 2012; Bernard and Seidler, 2013; E et al., 2012; Kruithof et al., 2023; Pierce et al., 2023; Stoodley and Schmahmann, 2009; Van Overwalle et al., 2020a, 2014). CBMAs have also been used to map cerebellar topography more comprehensively, providing indications that diverse motor and cognitive functions occupy distinct cerebellar subregions (Stoodley and Schmahmann, 2009), establishing the common division of cerebellum into motor and non-motor parts. Comparable large-scale meta-analyses have been performed within the socio-cognitive domain (Van Overwalle et al., 2020a, 2014). Recognizing that cerebellar behavioral topography largely reflects brain-wide connectivity, a CBMA adaptation – meta-analytic connectivity modeling (MACM) – has also helped reveal generalizable cerebellar coactivation networks across the brain (Balsters et al., 2014).

While early cerebellar meta-analyses focused on specific behaviours or task domains, a recent meta-analytic atlas (Van Overwalle et al., 2023) evaluated a wide range of domains using the activation coordinates reported in the NeuroSynth database (Yarkoni et al., 2011). However, both this and the earlier cerebellar CBMA work do not address the marked cerebellar reporting biases in the neuroimaging literature (Balsters et al., 2014; Desmond and Fiez, 1998; Van Overwalle et al., 2023) that need to be overcome for accurate, statistically valid behavioral mapping of the cerebellar cortex (Wang et al., 2025). The cerebellum has been systematically underrepresented in neuroimaging research (Wang et al., 2025). Historically, the cerebellum was often not retained in the field-of-view (FOV) of neuroimaging scans, as shifting the FOV to frontal regions improves data quality locally and increases cerebral ‘buffer’ (Wang et al., 2025), at the cost of cerebellar coverage (Desmond and Fiez, 1998; Van Overwalle et al., 2023). Through this design choice, a substantial bias has made its way into the neuroimaging literature (Van Overwalle et al., 2023). This neglect may be caused by several other factors, including: poor to no coverage of cerebellar functions during medical/neuroscience training, limiting understanding of its structure and function, multiple pipelines focusing on cerebral rather than cerebellar analysis, a lack of integration of cerebellar analysis tools into common pipelines, and ultra-high-resolution 7T MRI coils being unsuitable for posterior cerebellar imaging (Kraff and Quick, 2019), at least historically (Wang et al., 2025) (but see cerebellar imaging advances in (Priovoulos et al., 2023; Priovoulos and Bazin, 2023)). These factors contributed to the perception of the cerebellum as an unimportant structure, leading to its vast underrepresentation in the number of research papers and funded projects (Wang et al., 2025). This underrepresentation has led foci to be strongly skewed to superior cerebellar locations by cropping out posterior regions. Counterintuitively, a large proportion of cerebellar activation coordinates may thus come from studies that did not actively consider the cerebellum, or even those that consciously moved the FOV frontally to exclude inferior parts of the cerebellum. This biased neglect appears to have exacerbated the inherent unequal distribution of foci across the brain (Langner et al., 2014).

This means that the traditional implementation of ALE may report convergence in well-reported, superior cerebellar regions. Convergence in underreported, inferior regions is much less likely. This can lead to behavioral clusters that reflect, to an unknown extent, baseline reporting rates and obscure real behavioral signals. Therefore, in these cases, where the assumption of randomly distributed activations is violated (Eickhoff et al., 2009), ALE is statistically invalid.

As a result, despite extensive research, there is still no accurate and comprehensive meta-analytic synthesis of cerebellar activations across the neuroimaging literature that accounts for the systematic reporting bias. Such a synthesis is important to complement and further refine state-of-the-art mappings and parcellation studies by finding cerebellar activations that are generalizable despite conflicting results in individual studies. Hence, we propose an adaptation of ALE, cerebellum-specific ALE (C-SALE), that updates its null hypothesis to account for reporting biases. We apply this method to data from BrainMap (Laird et al., 2005b), a manually curated database that categorizes experiments into diverse behavioral domains. We assess replicability of our findings in NeuroSynth (Yarkoni et al., 2011). In doing so, we aim to provide answers to the following research questions: (1) For any given behavior, what cerebellar subregion(s) show activity convergence above chance relative to the rest of the cerebellum? (2) What spatial association is there between the subregions [found in (1)] and previous maps and atlases? and (3) For any given behavior, what area(s) of the brain are coactivated with the cerebellar subregion(s) [found in (1)] above chance? Together, answering these questions across two large-scale databases synthesizes the vast neuroimaging literature to pinpoint cerebellar involvement across a widespread range of behaviors.

## 2. Results

### 2.1 Study inclusion

We identified behavioral datasets by querying the BrainMap (Laird et al., 2005b) database in February 2024 for task-based fMRI and positron emission tomography (PET) data in healthy subjects, creating eligible datasets for five BDs and thirty-two subdomains. These behavioral (sub)domains refer to the division of tasks into human-understandable categories by the BrainMap team. Each subdomain includes tasks describing a distinct behavioral construct, and belongs to a more global BD (‘Action’, ‘Cognition’, ‘Emotion’, ‘Interoception’, ‘Perception’) (Fox et al., 2005; Laird et al., 2009, 2005b). Note that a single task or experiment can map to several (sub)domains. Overall, 1,109 unique studies (2,322 experiments, with 16,159 participants and 4,644 cerebellar foci) were included. The number of experiments per (sub)domain can be found in **Supplementary Figure 1a**.

We validated our method and replicated C-SALE maps in NeuroSynth (Yarkoni et al., 2011). From the NeuroSynth database, we selected 101 terms that describe behavioral and cognitive functions. In total, 5,898 experiments were included. The number of experiments per term can be found in **Supplementary Figure 1b**. For the overall number of experiments, coordinates, and subjects included in each (sub)domain and term, see *Data availability*.

### 2.2 Assessing improvements of C-SALE over classic ALE

Previous work has used ALE to map activations in behavioral (sub)domains to cerebellar subareas. The regular ALE method assumes equal spatial distributions of reported effects (foci). However, visualizing locations of cerebellar foci in BrainMap and voxels along the z-axis illustrates substantial spatial biases (**Figure 1a**). In the NeuroSynth database, we replicated this strong bias (**Supplementary Figure 2b**). Overall, the BrainMap and NeuroSynth cerebellar baseline distributions were strongly correlated (voxel-wise correlation: r = .96, *p* < .001). In both datasets, foci were strongly skewed towards superior regions. Along the x– and y-axes, distributions of foci largely mirrored those of cerebellar voxels although it appeared that right cerebellar foci were somewhat overrepresented (**Supplementary Figure 3a, b**). C-SALE considers these observed, unequal probabilities of finding foci at any voxel (**Figure 1b**). This baseline cerebellar probabilistic foci distribution, generated from domain-general foci reported in BrainMap, revealed two substantial probability hotspots in bilateral anterior lobules. Inferior-posterior regions had lowest probabilities, and the distribution was generally left-right symmetrical albeit slightly skewed to the right cerebellum. Note that these unequal distributions of reported effects were not unique to the cerebellum, but common across the brain (BrainMap: **Supplementary Figure 4** and NeuroSynth: **Supplementary Figure 5**).

**Figure 1:**
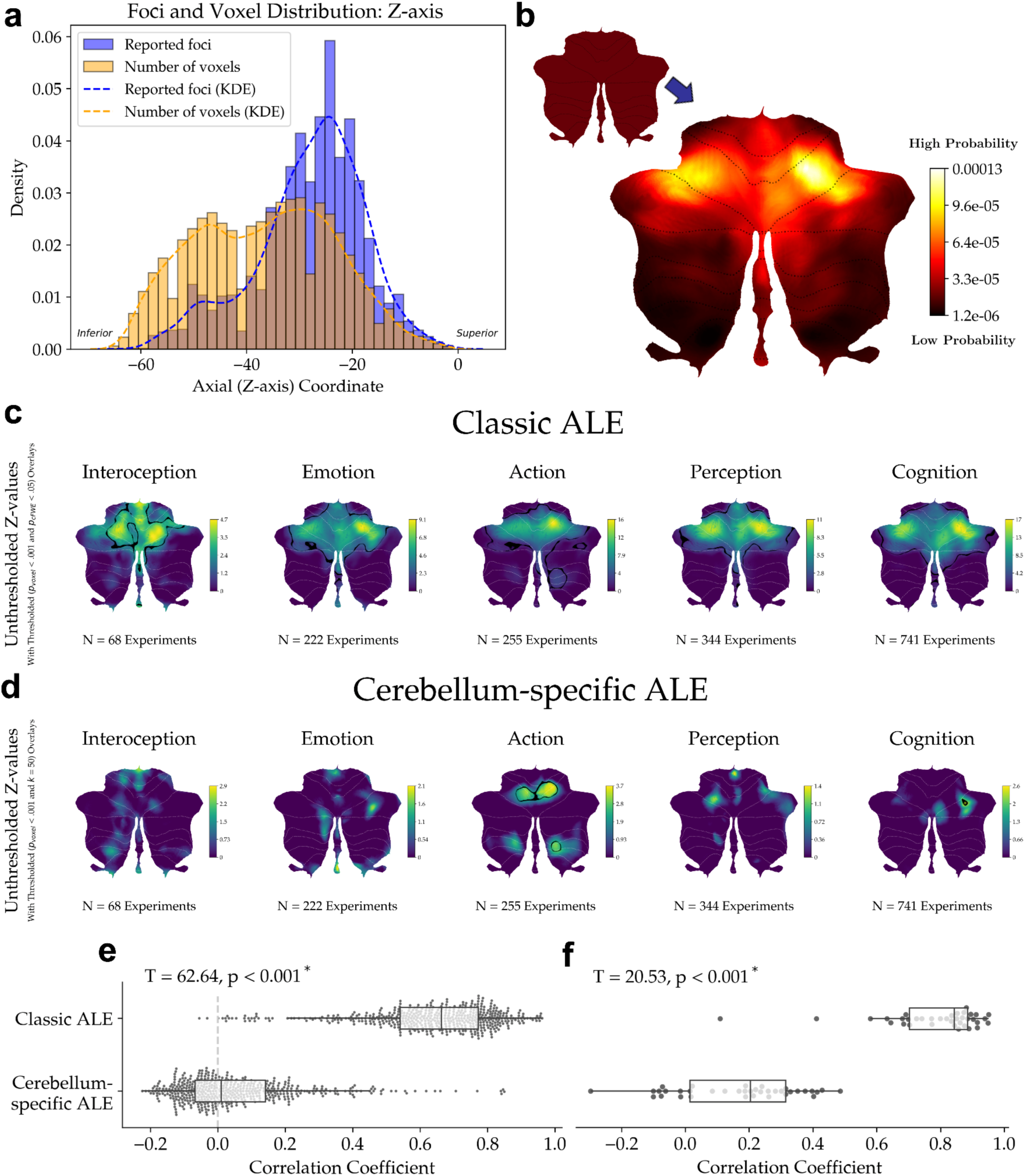
Improvements of the new cerebellum-specific ALE method over classic ALE for cerebellar meta-analysis. Methodological assessment of cerebellum-specific activation-likelihood estimation (C-SALE). (**a**) Maps the distribution of reported effects versus the number of cerebellar voxels across the Z-axis. (**b**) Illustrates the null hypotheses of classic ALE (smaller flatmap) and C-SALE (larger flatmap). (**c, d**) Show differences between classic ALE (**c**) and C-SALE (**d**) behavioral domain (BD) maps. Unthresholded z-maps are shown with an outline of clusters that reached convergence (*p_voxel_* < .001 and k = 50 for C-SALE; and *p*_cFWE._ < .05 using a height threshold of *p_voxel_* < .001 for classic ALE). BDs are ordered by sample size from small to large. (**e**) Shows distributions of correlation coefficients between each pair of unthresholded (sub)domain z-maps. Correlations were significantly lower in C-SALE (T = 62.64; *p* < .001*). The vertical dotted line illustrates the null assumption that within-cerebellum spatial correlation centers on 0.0 in accurate within-cerebellar CBMAs. (**f**) Shows correlation of unthresholded z-maps with the biased baselines distribution (pictured in **b**). C-SALE resembled the baseline significantly less (T = 20.53; *p* < .001*). For C-SALE validation in NeuroSynth (Yarkoni et al., 2011), see **Supplementary Figure 2**. KDE = kernel density estimation.

### 2.3 Comparing ALE and C-SALE methods

Whereas classic ALE constructs null models by sampling random cerebellar gray matter (GM) coordinates, C-SALE uses the biased baseline to sample GM coordinates weighted by the probability distribution. Essentially, comparing peak activation coordinates within a behavioral (sub)domain to this null model tests where in the cerebellum activity converges beyond baseline probabilities of cerebellar reported effects. Comparing BD maps for classic ALE (**Figure 1c**) and C-SALE (**Figure 1d**) revealed improved specificity of our new method. First, in ALE most of the superior half of the cerebellum reached convergence invariant of BD and resembled the baseline reporting rates to a large (but BD-specific) extent. In C-SALE, significant convergence was observed for ‘Action’ in bilateral lobules V-VI and right VIIIa-b, as well as for ‘Cognition’ in right Crus I. The other BDs did not reach the threshold for significance. Examining unthresholded z-maps for these BDs illustrated local peaks of (subthreshold) convergence in inferior regions (**Figure 1c, d**). Within NeuroSynth, a selected subset of five terms (‘Emotion’, ‘Movement’, ‘Perception’, ‘Social Cognition’, and ‘Working Memory’) illustrated similarly unspecific convergence in superior regions in ALE. These regions again resembled baseline reporting rates within the dataset (**Supplementary Figure 2b**) to a large (but term-specific) extent (**Supplementary Figure 2c**). In contrast, C-SALE found convergence for all terms but did so in smaller, distributed cerebellar regions that bore no direct resemblance to the baseline reporting rates (**Supplementary Figure 2d**).

We compared spatial correlations of unthresholded maps for pairs of BDs and subdomains. Whereas some spatial correlation across behaviors is expected, excessive correlations indicate a lack of specificity given the premise of within-cerebellar CBMA. ALE subdomains illustrated such excessive (median = .66, IQR = [.54, .77]) correlations, whereas the correlations were much lower (T = 62.64, *p* < .001) in C-SALE (median = .05, IQR = [-.07, .14]) (**Figure 1e**). ALE maps were more strongly correlated with the biased baseline distribution (median = .78, IQR = [.70, .88]) than C-SALE maps (median = .17, IQR = [.02, .31]) (T = 20.53, *p* < .001) (**Figure 1f**). Within NeuroSynth, similar patterns were observed: ALE maps showed excessive between-term correlations (median = .74, IQR = [.63, .83]) (**Supplementary Figure 2e**), whereas these were much lower (T = 273.04, *p* < .001) in C-SALE (median = .01, IQR = [-.07, .12]) (**Supplementary Figure 2e**). ALE maps (median = .85, IQR = [.77, .91]) also correlated much more strongly (T = 37.94, *p* < .001) to the biased baseline than C-SALE maps (median = .11, IQR = [-.05, .24]) (**Supplementary Figure 2f**).

Full hierarchically clustered correlation heatmaps for BrainMap are reported for BDs (**Supplementary Figure 6)** and subdomains (**Supplementary Figure 7).** Both illustrate that whereas classic ALE finds high correlations across most combinations, C-SALE primarily finds high correlations between related behaviors. As we established the improved performance of C-SALE, we continue to only report results for this method. In turn, the BD maps reported in **Figure 1d** represent BD-level results.

### 2.4 Cerebellum-Specific Activation Likelihood Estimation (C-SALE)

Next, we use C-SALE to find if and where reported activations in each behavioral (sub)domain converged. C-SALE results for BDs (**Figure 1d**; **Figure 2a**) and subdomains (**Figure 2b**, **Figure 3**) are reported on the cerebellar flatmap. For both, unthresholded maps are plotted alongside an outline of thresholded (*p_voxel_* < .001 and k = 50) clusters (**Figure 1d**; **Figure 3**). Binary locations of convergence for BDs are illustrated in **Figure 2a**, and **Figure 2b** illustrates how subdomain convergence maps to the BD they are organized under in BrainMap. For BDs, full unthresholded and thresholded (*p_voxel_* < .001 and k = 50) z-maps are reported in **Supplementary Figure 8a-b**. Full z-maps for subdomains are provided in **Supplementary Figures 9-11.**

**Figure 2:**
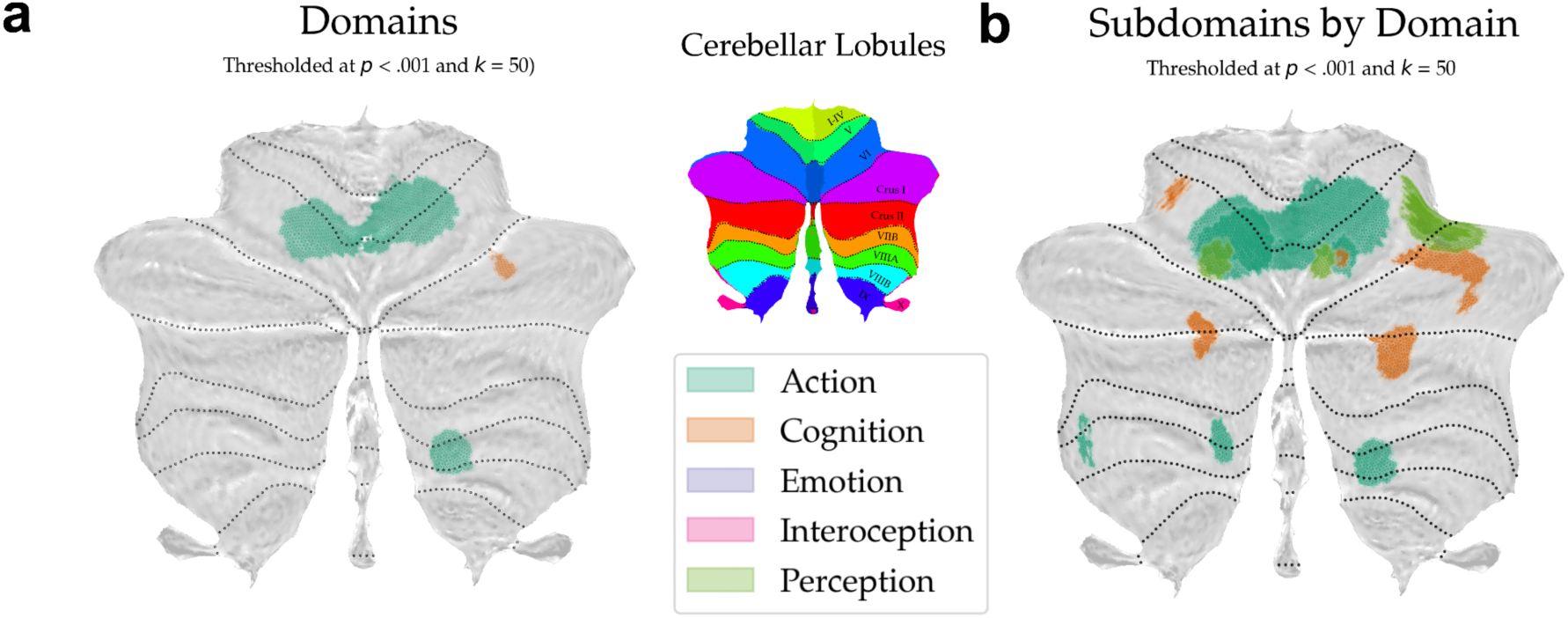
Overlap of convergence across behavioral (sub)domains. (**a**) Summary of converging cerebellum-specific ALE (C-SALE) maps for behavioral domains (BDs) (**Figure 1**). Binary locations of convergence (*p_voxel_* < .001 and k = 50) are plotted to a common flatmap. The next panel shows the cerebellar lobular definition (Diedrichsen et al., 2009; Larsell and Jansen, 1973; Schmahmann et al., 2000) (top) and a color mapping for BDs (bottom). In (**b**), converging subdomain results (see **Figure 3**) are summarized by BD. Binary locations of convergence are colored by the BD the subdomains belong to and plotted onto a common flatmap. Legends apply to both **a** and **b**. Only BDs and subdomains that reached convergence are shown.

**Figure 3:**
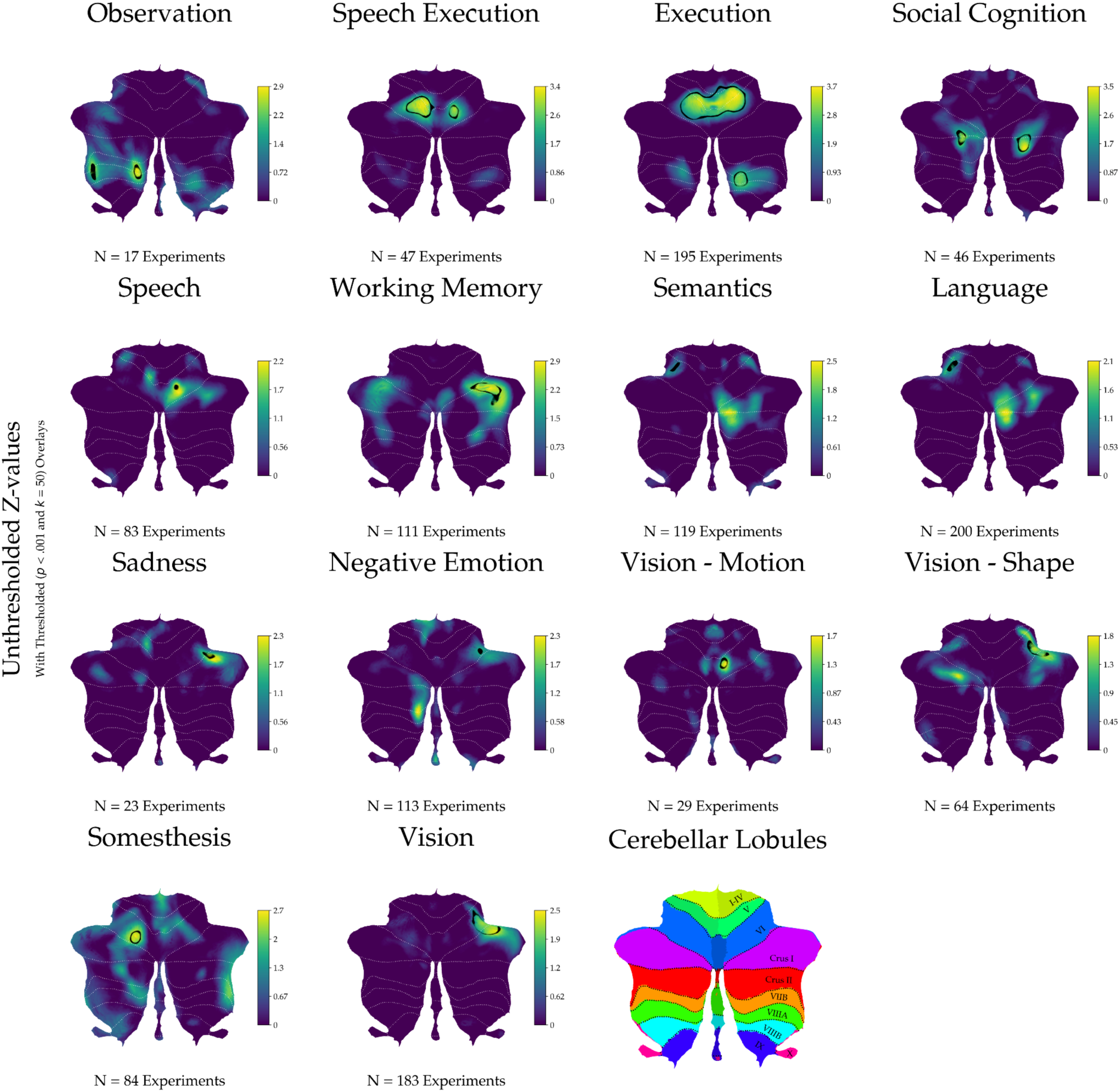
Cerebellum-specific ALE maps for behavioral subdomains. Unthresholded z-maps are shown with an outline of the locations that reached convergence (*p_voxel_* < .001 and k = 50). Subdomains are first ordered by behavioral domain (alphabetically) and then by sample size (from small to large). The last panel shows the cerebellar lobular definition (Diedrichsen et al., 2009; Larsell and Jansen, 1973; Schmahmann et al., 2000). For equivalent maps for NeuroSynth (Yarkoni et al., 2011) terms, see **Supplementary Figure 12**.

### 2.5 Behavioral subdomain topography of the cerebellum

Behavioral subdomains aim to capture more specific behavioral constructs. Hence, after investigating BDs, we considered cerebellar regional convergence in subdomains and found significant convergence in fourteen out of thirty-two subdomains (**Figure 3**). As **Figure 2b** and **Supplementary Figure 10** illustrate, these subdomains belonged to ‘Action’, ‘Cognition’, ‘Emotion’, and ‘Perception’. Clusters were spread relatively equally across left-right and inferior-superior regions, somewhat favoring superior and right cerebellar regions. Subdomains within BDs tended to map to similar cerebellar subregions (**Figure 3**). Within ‘Action’, ‘Observation’, ‘Speech Execution’, and ‘Execution’ reached convergence. ‘Observation’ converged in two clusters in left VIIb and VIIIa, ‘Speech Execution’ in left V-VI and right VI, and ‘Execution’ in bilateral (including vermal) V-VI and right VIIb-VIIIb. Next, within ‘Cognition’, ‘Social Cognition’, ‘Working Memory’, ‘Language’, ‘Speech’, and ‘Semantics’ converged. ‘Social Cognition’ converged in bilateral Crura I-II, ‘Working Memory’ in right VI-Crus I, ‘Speech’ in right VI, and ‘Language’ and ‘Semantics’ in left VI. Within ‘Emotion’, ‘Sadness’ converged in right VI-Crus I, and ‘Negative Emotion’ converged in right VI. Within ‘Perception’, ‘Vision – Motion’ converged in right VI, whereas ‘Vision – Shape’ and ‘Vision’ converged in right V-Crus I. Lastly, ‘Somesthesis’ converged in left VI.

We also evaluated C-SALE in all 101 selected NeuroSynth terms, finding convergence in thirty-eight. All maps with significant convergence can be found in **Supplementary Figure 12.** These maps corresponded closely to BrainMap (sub)domain maps, as evidenced by many high and significant correlations after accounting for spatial autocorrelation (SA) (**Supplementary Figure 13**). To highlight the similarity between BrainMap and NeuroSynth mappings, we plotted z-maps for the ten pairs showing highest spatial correlations between unique maps (**Supplementary Figure 14**).

### 2.6 Spatial stability of C-SALE maps

Then, to assess how consistently behaviors map to cerebellar subareas, we performed a complementary set of repeated subsampling analyses. Specifically, for several subsampling parameters, we repeated C-SALE analyses in fifty random subsets of experiments per (sub)domain. We then compared spatial correlations between each of the fifty subsampled C-SALE z-maps for all parameters and (sub)domains. To account for the variability in the number of experiments across (sub)domains, two complementary subsampling strategies were used, controlling: **(1)** the absolute number of experiments and **(2)** the proportion of experiments. For BDs, we illustrated how stability develops along a range of absolute **(1)** and proportional **(2)** parameters. For subdomains, running all parameters was deemed too costly, so we used the following arbitrary parameters **(1)** n_subsample_ = 50; and **(2)** n_subsample_ = 0.2 * n_(sub)domain_. The repeated subsampling datasets correspond to those reported for the main analysis but were only run for (sub)domains with n_experiments_ ≥ 60, leaving five of five BDs and seventeen of thirty-two subdomains.

Spatial correlations of unthresholded subsample maps revealed that most BDs (**Figure 4a, b**) and subdomains (**Figure 4d, e**) were moderately stable. First, BD mapping stability naturally increased as sample sizes increased (**Figure 4a, b**). This was partially driven by increasing proportions of overlapping experiments across subsamples, illustrated by the large increase in mapping stability between n = 25 and n = 50 subsampling for ‘Interoception’ (the smallest BD) relative to other BDs (**Figure 4a**). Together, the results also show that larger CBMA analyses tend to be more stable (Eickhoff et al., 2016). Thus, varying the subsampling proportion allowed for comparison of mapping stability between BDs. Across BDs stability generally increased as subsampling proportions increased (**Figure 4b**). ‘Action’ stood out for high stability. Stability was high even at low proportions and appeared close to plateauing at the .8 proportion (median correlation = .94; SD ± .02. To a lesser extent, stability of ‘Cognition’ increased faster than other BDs with increasing subsampling proportions. This was particularly evident at high (i.e., .8) proportions with a spatial correlation of .88 (SD ± .06). Median spatial correlations were somewhat lower for ‘Interoception’ (.77; SD ± .06), ‘Emotion’ (.79; SD ± .08), and ‘Perception’ (.68; SD ± .10). These results reveal domain-specific differences in the consistency of cerebellar localizations.

**Figure 4:**
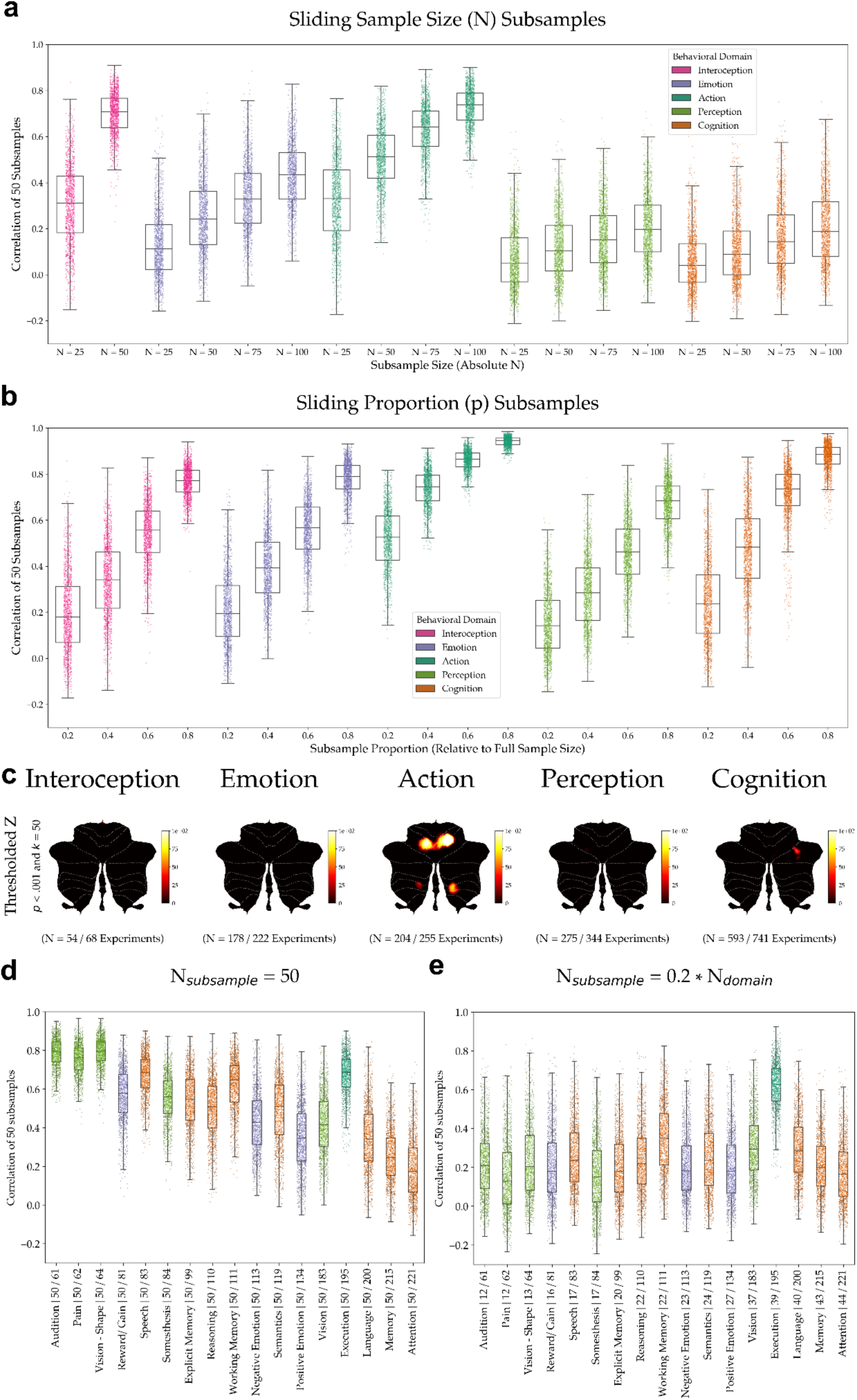
Stability of cerebellum-specific ALE maps. To assess stability of cerebellum-specific ALE (C-SALE) maps, repeated subsampling analyses were performed. Two different subsample strategies were used for both behavioral domains (BDs) and subdomains. One sampled an absolute number of experiments for each subsample (**a, d**) and the other a proportion of overall (sub)domain sample size (**b, e**). (**a**) Shows spatial correlation between fifty random subsamples at different (sliding) absolute sizes (n = 25, 50, 75, and 100) within BDs. In turn, (**b**) shows spatial correlations for different (sliding) proportions of overall BD sample size (proportion (p) = 0.2, 0.4, 0.6, and 0.8). (**c**) To highlight stability of thresholded (*p_voxel_* < .001 and k = 50) z-maps, percentages of the highest (p = 0.8) proportion subsamples that reached convergence at any given voxel were mapped to a common flatmap per BD. In (**d, e**) stability of unthresholded maps for subdomains are shown. Here, fifty random subsamples were used as input to C-SALE, consisting either of fifty experiments per subsample (**d**), or of a number proportional to subdomain sample size (p = .2) (**e**).

### 2.7 Consistency of voxel-wise cerebellar convergence

We also assessed how consistently convergence exceeded the threshold for significance (*p_voxel_* < .001 and k = 50). Specifically, we mapped subsamples per (sub)domain to a common flatmap, visualizing proportions of subsamples that reached convergence at each voxel for BDs (**Figure 4c, Supplementary Figure 15**) and subdomains (**Supplementary Figure 16**). Focusing on BDs, whereas unthresholded maps were rather comparable (based on relatively high spatial correlations (**Figure 4b**)), thresholds were only reached consistently in ‘Action’, and to a lesser extent in ‘Cognition’. In the n_subsample_ = 50 analyses, larger subdomain sizes (and thus smaller proportions) led to decreased stability. ‘Execution’ stood out as being remarkably stable. Generally, .2-proportion subdomain and BD (**Figure 4b**) subsamples were similarly stable. ‘Execution’ was again elevated above other subdomains. Notably, stable unthresholded maps (e.g., ‘Action’, ‘Execution’, ‘Vision’, ‘Working Memory’, and ‘Language’) reached convergence (**Figures 1-3**). Together, these results illustrate that for many behaviors, large samples are necessary to find consistent regions of convergence. For sampling sizes and proportions, see *Data availability*.

### 2.8 Correspondence of C-SALE maps to the MDTB and the hierarchical cerebellar atlas

To contextualize meta-analytical cerebellar mappings, we next compared them with several established mappings and parcellations, including the cerebellar multi-domain task-battery (MDTB) (King et al., 2019) (**Figure 5a**), mid-granularity cerebellar hierarchical atlas (Nettekoven et al., 2024) (**Figure 5b**). First, MDTB comparisons revealed two main clusters, largely separating ‘Cognition’ and ‘Emotion’ from ‘Action’ and ‘Perception’ (including subdomains) (**Figure 5c**). Of 592 combinations, ninety-eight were correlated significantly. Most notably, positive significant correlations (*p_variogram, FDR_* < .05) were observed between meta-analytical and MDTB maps including ‘Social Cognition’ with Animated Movie (r = .486) and Theory of Mind (r = .478), ‘Action’ and ‘Execution’ with Finger Sequence (r = .473 and .461, respectively), ‘Phonology’ with Verb Generation (r = .469) and ‘Working Memory’ with Math (r = .394), Verb Generation (r = .331) and 2-back (r = .294). For the mid-granularity hierarchical atlas (Nettekoven et al., 2024) (**Figure 5b**), two main clusters separated ‘Action’ and ‘Execution’ from all other (sub)domains (**Figure 5d**). Of 592 comparisons, nineteen were significant after accounting for SA and multiple comparisons. ‘Action’ and ‘Execution’ corresponded significantly to M2 and M3 parcels. Among others, significant correspondence was also found between ‘Action Observation’ and A1, ‘Explicit Memory’ and S5, and ‘Social Cognition’ and S2 and S3. For putative behavioral labels of these parcels, see **Supplementary Figure 17c**. For comparisons with cerebellar resting-state atlases (Buckner et al., 2011; Ji et al., 2019), lobular definitions (Diedrichsen et al., 2009; Larsell and Jansen, 1973; Schmahmann et al., 2000), and functional gradients (Guell et al., 2018b), see ***sections 2.81 – 2.8.3*** and **Supplementary Figures 17-19.** Note that for continuous comparisons, spatial correlations were calculated between every *C-SALE map* and *target map*. For parcellations, heatmaps report mean z-values for each C-SALE map within each parcel. For both types of comparison, reported heatmaps were hierarchically clustered. Statistical significance of spatial correspondence was assessed after accounting for SA and multiple comparisons. For full correlations or mean z-values and *p*-values for all comparisons, see *Data availability*.

**Figure 5:**
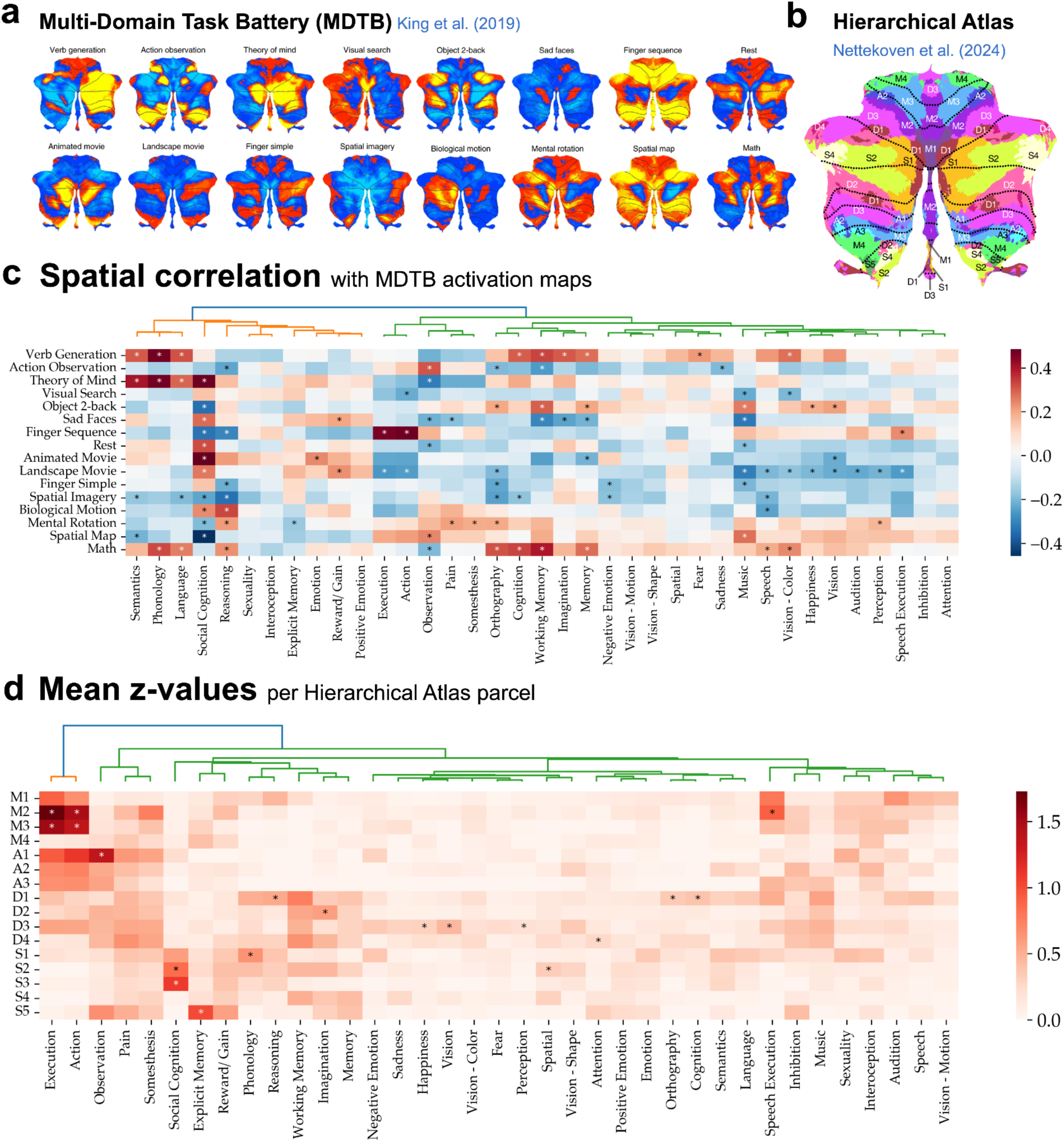
Correspondence of cerebellum-specific ALE maps with the multi-domain task battery maps and the cerebellar hierarchical atlas. To contextualize behavioral maps and relate them to existing cerebellar parcellations, we compared unthresholded cerebellum-specific ALE (C-SALE) z-maps with the multi-domain task battery (MDTB) (King et al., 2019) **(a)** and the symmetrical mid-granularity hierarchical cerebellar atlas (Nettekoven et al., 2024) **(b)** that was merged across left and right parcels. **(c)** Illustrates spatial correlations between each (sub)domain’s z-map and MDTB group-level task activation map^60^. Variograms were used to account for spatial autocorrelation (SA). Asterisks indicate significant correlations (*p_variogram, FDR_* < .05). **(d)** Illustrates mean z-values of each (sub)domain’s C-SALE map within each parcel. Asterisks indicate significant combinations (*p_variogram, FDR_* < .05). Both heatmaps **(c, d)** were hierarchically clustered. For putative functional labels of the hierarchical atlas parcels in (**d**), see **Supplementary Figure 17c**. (**a, b**) were adapted from (King et al., 2019), and (Nettekoven et al., 2024) after obtaining permission from the authors.

#### 2.8.1 Resting-state atlases

Comparisons with the resting-state atlases (Buckner et al., 2011; Ji et al., 2019) (**Supplementary Figure 17a, b**) both produced two clusters separating ‘Action’ and ‘Execution’ from other C-SALE maps. Notably, apart from primarily overlapping somatomotor networks (SMN), ‘Action’ maps also substantially overlapped other networks: the Dorsal and Ventral Attention Networks (DAN and VAN) (Buckner et al., 2011), as well as both visual networks, the Cingulo-Opercular network (CON), and DAN (Ji et al., 2019). The frontoparietal network (FPN) of both parcellations corresponded to several C-SALE maps: ‘Cognition’, ‘Reasoning’, and ‘(Working) Memory’. ‘Language Orthography’ and ‘Phonology’ mapped uniquely to the Buckner and Ji parcellation’s FPN, respectively. ‘Social Cognition’ mapped to the Default-Mode Network (DMN) in both parcellations.

#### 2.8.2 Correspondence with lobular boundaries

To assess how well cerebellar lobular boundaries define behavioral mapping, we examined spatial correspondence between C-SALE maps and lobules (Diedrichsen et al., 2009). First, calculating mean z-values across the deterministic lobular segmentation revealed three clusters (**Supplementary Figure 18a**). ‘Action’ and ‘Somesthesis’ (including ‘Pain’) were separated from two rather mixed groups of behaviors. Of these, five mapped to vermal (Reasoning) and right (‘Cognition’, ‘Language Phonology’, ‘Memory’, and ‘Spatial Cognition’) cerebellum. ‘Speech Execution’ mapped to left VI, whereas ‘Action Observation’ mapped to left VIIIa. In total, only seven of 1184 comparisons were significant. Spatial correlations with the probabilistic lobular segmentation (**Supplementary Figure 18b**) also revealed three clusters. These separated aspects of ‘Action’ and ‘Somesthesis’ (as well as ‘Perception’, ‘Audition’, and ‘Music’) from two behaviorally diverse clusters. Several correlations stood out. Most notable were those with left Crus II, especially in ‘Cognition’, including ‘Language – Phonology’, ‘Spatial Cognition’, and ‘Memory’. ‘Speech Execution’ corresponded to vermal VI, whereas ‘Execution’ corresponded to left VI. In total, only eleven of 1184 comparisons were significant.

#### 2.8.3 Loading onto cerebellar functional gradients

Behavioral C-SALE clusters occupied distinct locations along cerebellar functional gradients (Guell et al., 2019, 2018b) (**Supplementary Figure 19**). Differences in loadings across behavioral domains (BDs) were evident across both gradients, but primarily G1. ‘Action’ and its subdomains ‘Execution’, ‘Speech Execution’, and ‘Observation’ loaded towards the motor anchor of G1. Whereas the former three also mapped to the task-unfocused anchor of G2, ‘Observation’ instead loaded more centrally. This loading primarily corresponded to VAN and DAN of (Buckner et al., 2011), contrasting stronger overlap with SMN and VAN for ‘Action’, ‘Execution’ and ‘Speech Execution’. ‘Vision’, ‘Vision – Motion’, ‘Vision – Shape’, and Somesthesis (all perception subdomains) mapped more centrally on G1 and G2. All overlapped VAN and FPN, except ‘Vision – Motion’ that only had a tiny cluster within VAN. ‘Cognition’ and its subdomains loaded to central-to-positive locations on G1 (towards the DMN). Loadings across both gradients illustrated the diversity of ‘Cognition’: whereas ‘Language’, including ‘Language – Speech’ and ‘Language – Semantics’, loaded closer to the motor anchor of G1 (overlapping VAN), ‘Working Memory’ loaded to positive G1 values (primarily FPN, partially DMN). Both occupied central G2 loadings. ‘Social Cognition’ on the other hand mapped cleanly to DMN and task-unfocused anchors of both gradients, completely overlapping Buckner’s DMN. ‘Emotion’ subdomains ‘Negative Emotion’ and ‘Sadness’ mapped centrally on both gradients, overlapping Buckner’s FPN.

### 2.9 Meta-analytic connectivity modeling

Since much of cerebellar functional topography reflects connectivity, we last used C-SALE clusters as seeds for whole-brain MACM analyses, revealing brain-wide coactivation networks. As in C-SALE, instead of assuming spatial homogeneity, MACM adjusts its null model to reflect the unequal probability distribution of finding foci across the brain (**Supplementary Figure 4**). Here, the overall set of analyses was restricted to (sub)domains showing cerebellar convergence (*p_voxel_* < .001 and k = 50) in the main C-SALE analyses (‘Action’, ‘Cognition’, and fourteen subdomains). Experiments within each (sub)domain were then restricted to those that had at least one coordinate within the regions converging in C-SALE analysis (**Figures 1-3**). To provide stable MACM analyses, an additional prerequisite was that at least seventeen such experiments existed (Eickhoff et al., 2016). This restricted the set of valid analyses to five (sub)domains including ‘Action’, ‘Execution’, ‘Execution Speech’, ‘Working Memory’, and ‘Vision’. Thresholded MACM (*p_voxel_* < .001 and k = 50) coactivation maps for all (sub)domains can be found in **Figure 6**. To aid interpretation, common subcortical (Tian et al., 2020), cerebral cortical (García-Cabezas et al., 2019; Ji et al., 2019), and cerebellar (Diedrichsen et al., 2009) parcellations are plotted in **Figure 6a**.

**Figure 6:**
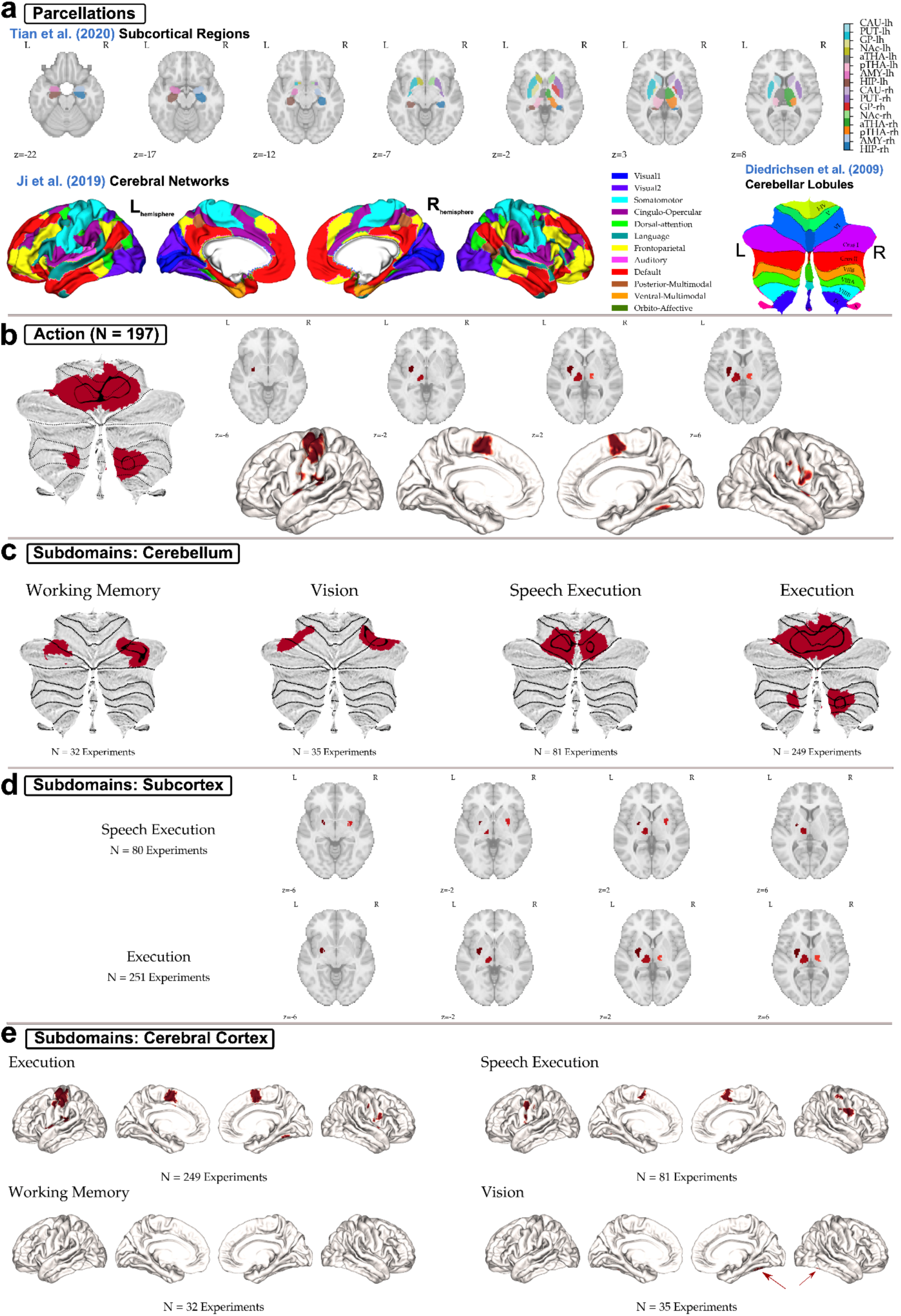
Whole-brain coactivation networks from cerebellum-specific ALE clusters. Whole-brain coactivation networks for behavioral domains (BDs) and subdomains. Specifically, the maps show results from meta-analytic connectivity modeling (MACM) using the whole-brain probability distribution for construction of the null models (see **Figure 1**, **Supplementary Figure 4**). Clusters of activity convergence in cerebellum-specific ALE (C-SALE) (**Figures 2, 3**) were used to restrict experiments for each MACM analysis (MACM seeds, **Supplementary Figure 20**). Across the figure, black outlines are overlaid on the cerebellar flatmaps to illustrate seed regions. (**a**) Shows subcortical, cerebral, and cerebellar parcellations used to contextualize MACM maps. These are a subcortical regional parcellation (Tian et al., 2020), cerebral cortical network parcellation (Ji et al., 2019), and the lobular cerebellar atlas (Diedrichsen et al., 2009). (**b**) Shows full MACM maps across cerebellum, subcortex, and cerebrum for ‘Action’. (**c-e**) Show MACM maps for subdomains, for cerebellum (**c**), subcortex (**d**), and cerebral cortex (**e**), respectively. In (**d**) only ‘Execution’ and ‘Speech Execution’ are shown, as they were the only subdomains to reach subcortical convergence. In (**e**), ‘Working Memory’ did not reach cerebral convergence. Convergence for ‘Vision’ is marked by arrows to aid visibility. Abbreviations (**a**): lh = left hemisphere; rh = right hemisphere; HIP = hippocampus; AMY = amygdala; pTHA = posterior thalamus; aTHA = anterior thalamus; NAc = nucleus accumbens; GP = globus pallidus; PUT = putamen; CAU = caudate nucleus.

Cerebellar MACM maps illustrated increased regions of convergence relative to their seeds (black outlines; **Figure 6b, c)**. Although such patterns are expected, it is noteworthy that this revealed additional and symmetrical coactivating cerebellar regions across ‘Action and ‘Execution’ (both in left VIIb-VIIIb), ‘Working Memory’ (left VI-Crus I), and ‘Vision’ (left V-Crus I). Only ‘Action’, ‘Execution’, and ‘Speech Execution’ MACMs converged in the subcortex (**Figure 6b, d**). All overlapped the left caudate, nucleus accumbens and anterior thalamus. Whereas ‘Action’ and ‘Execution’ MACMs convergence additionally partly overlapped the right nucleus accumbens and anterior thalamus, ‘Speech Execution’ instead overlapped the right caudate and, partly, the bilateral putamen (**Supplementary Figure 21**). The seeds of ‘Action’, ‘Execution’, ‘Speech Execution’ and ‘Vision’ showed significant MACM convergence in the cerebral cortex (**Figure 6b, e**). Across ‘Action’, ‘Execution’, and ‘Speech Execution’, MACM maps were highly similar, localized mainly to known sensorimotor areas. The MACM for the ‘Vision’ seed showed coactivation with the fusiform cortex. However, this cluster was a continuation of the cerebellar cluster to this part of the cerebral cortex. Overall, we showed that cerebellar behavioral clusters are systematically coactivated with distinct brain regions in cerebral cortex and subcortex.

#### 2.9.1 Correspondence of MACM maps to brain-wide parcellations

Finally, to interpret whole-brain coactivation networks, we assessed their spatial correspondence with subcortical regions (Tian et al., 2020), microstructural types (García-Cabezas et al., 2019), and cerebral functional networks (Ji et al., 2019). For each (sub)domain, we report the proportion of converging voxels that map to each subcortical area (**Supplementary Figure 21a**), or the proportion of significant vertices that map to each cortical type and Cole-Anticevic network (**Supplementary Figure 21b, c**). Meta-analytic connectivity modeling (MACM) maps in ‘Action’, ‘Execution’, and ‘Speech Execution’, converged in similar subcortical regions, at least partially due to the high number of overlapping experiments. All three converged in the left caudate nucleus, nucleus accumbens and anterior thalamus (**Supplementary Figure 21a**). ‘Speech Execution’ also overlapped substantially with the right caudate, whereas ‘Action’ and ‘Execution’ showed more overlap with the right nucleus accumbens and anterior thalamus. In the cerebral cortex, ‘Action’, ‘Execution’, and ‘Speech Execution’ MACM maps were primarily colocalized with Eulaminate (EU) II-III cortices. They partially mapped to the Koniocortex. ‘Vision’ mapped highly preferentially to EUI (**Supplementary Figure 21b**). ‘Vision’ mapped fully to the Secondary Visual network (**Supplementary Figure 21c**).

## 3. Discussion

The aim of the current study was to complement the behavioral topography of the cerebellum by summarizing the extensive neuroimaging literature. To do this, we used an adaptation of the ALE meta-analytic method, called C-SALE, that accounts for cerebellar reporting biases in the neuroimaging literature. Specifically, our method identified cerebellar regions where activity in behaviors converged beyond baseline activation rates. This was achieved by comparing the experimental foci for each behavior with null models derived from the biased probability distribution of domain-general foci. Using this method, we created a meta-analytic mapping of the cerebellum across thirty-seven behavioral (sub)domains in BrainMap (Laird et al., 2005b). We validated our approach for 101 behavioral terms in NeuroSynth (Yarkoni et al., 2011), showing highly consistent results. Standard ALE overreported convergence in superior cerebellar regions in both databases, likely driven by historical cerebellar neglect. Our updated meta-analytical method accounted for the bias and substantially shifted the locations of cerebellar behavioral convergence. Our meta-analytic maps complement previous cerebellar parcellations, which focused on distinguishing functional boundaries in the cerebellum, by identifying regions that are consistently activated in specific behaviors across the neuroimaging literature.

### 3.1 Methodological implications of C-SALE for cerebellar mapping

Across both databases, classic ALE was found unsuitable for cerebellar CBMA, since its null assumption (equally distributed foci) (Eickhoff et al., 2009) was violated. The unequal distribution of foci across the brain (Langner et al., 2014) was likely exacerbated by incomplete cerebellar coverage. Previous cerebellar CBMAs have often – but not always – used cerebellar coverage as an inclusion criterion. When this is not done, convergence may be overreported in superior regions (and vice versa). As illustrated in the ALE-to-C-SALE comparisons, behaviors may then be wrongly assigned to cerebellar subregions. Given the baseline probability hotspots in both BrainMap and NeuroSynth, convergence in regions V and Crus I is most at risk of being overreported. For future CBMAs, it is important to account for unequal distributions of reported effects alongside explicitly verifying full (cerebellar) coverage. Although manually curated CBMAs often do the latter, improving data depth in inferior regions, the former is important to account for inherent brain-wide heterogeneity in reporting, and potentially, activity patterns. Our method can be flexibly adapted to analyses at the whole-brain-level or any brain ROI to pave the way for accurate and unbiased meta-analyses, ultimately advancing future research on cerebellar function and its role in cognition and behavior.

Recently, a similar cerebellar meta-analytical study was published (Van Overwalle et al., 2023). Here, CBMAs across the NeuroSynth database (Yarkoni et al., 2011) were used to create a meta-analytic cerebellar parcellation. This study aimed to address reporting biases by comparing ALE-scores at the voxel-level (Van Overwalle et al., 2023). However, we highlight certain limitations of this approach that likely influences the reliability of the results. First, any parcellation assumes that every cerebellar voxel is involved in just one function. In contrast, state-of-the-art mapping studies (Boillat et al., 2020; Saadon-Grosman et al., 2024) indicate that overlap of cerebellar involvement across behaviors is common. Hence, a statistical summary of cerebellar activations in the literature should not be limited by this constraint. Secondly, we replicated the reporting biases revealed in the NeuroSynth atlas study and showed that these biases led to nonspecific and inaccurate maps that resembled the baseline to an analysis-specific extent (**Supplementary Figures 2**). ALE values are furthermore dependent on experimental sample size and experimental consistency (Eickhoff et al., 2016), as shown by our subsampling analyses (**Figure 4**). Together, comparing these biased ALE scores, even at the same spatial location (Van Overwalle et al., 2023), can thus not create an unbiased atlas.

Lastly, the atlas’ assumption that no biased and systematic relation between neglect of the cerebellum and task domains exists (Van Overwalle et al., 2023) needs further examination. Many have pointed out how the cerebellum has historically been neglected in non-motor functions (see primarily the consensus paper series (Adamaszek et al., 2017; Arleo et al., 2023; Caligiore et al., 2017; Koziol et al., 2014; Leto et al., 2015; Mariën et al., 2014; Van Overwalle et al., 2020b) and (Wang et al., 2025)). We briefly explored this notion, mapping reported effect locations versus cerebellar voxels for each BrainMap BD separately (**Supplementary Figure 22**). Although conflated by behavioral signal, differences in foci-to-voxel distributions across BDs underline the need to formally investigate biased cerebellar neglect and not *a priori* assume its absence.

### 3.2 C-SALE helps refine cerebellar behavioral topography through literature synthesis

In BrainMap, sixteen of thirty-seven behavioral (sub)domains reached significant convergence in specific cerebellar subregions. Importantly, this implied heightened regional activity relative to the cerebellum rather than to the whole brain. Briefly, our localizations support cerebellar subregional roles across a great diversity of behaviors, in line with a growing consensus about cerebellar involvement in a plethora of behaviors (Adamaszek et al., 2017; Koziol et al., 2014; Mariën et al., 2014; Van Overwalle et al., 2020b). Specifically, within BrainMap, we report highly stable ‘Action’ localizations (including ‘Execution’ and ‘Speech Execution’) consistent with an established dual motor mapping (Adrian, 1944; Buckner et al., 2011; Guell et al., 2018b). These motor representations are broken up by bilateral posterior-lateral cerebellar subregions predominantly involved in ‘Emotion’ and ‘Cognition’ (Baumann and Mattingley, 2012; Guell et al., 2018a; Manoli et al., 2025; Pierce et al., 2022; Van Overwalle et al., 2020a, 2014). These cerebellar regions, primarily Crura I-II, have received considerable attention due to their role in associative functions. These functions are evolutionary underlain by connectivity with integrative, transmodal networks across the brain (Balsters et al., 2013; Magielse et al., 2022; Ramnani, 2006; Strick et al., 2009) and characterized by extensive primate-general expansion (Magielse et al., 2023), exceeding that of the frontal cortex (Bush and Allman, 2004). This suggests these areas’ relevance for sophisticated behavior. Lastly, we report convergence within ‘Perception’. We interpret the specific locations of convergence more extensively in *section 3.3*.

It is important to note that in validating C-SALE in the NeuroSynth database, we found thirty-eight behavioral terms that converged in cerebellar subregions. Although we focused on BrainMap because its manual curation added to categorization specificity, our analyses showed that the behavioral categorization in both datasets provided similar results, illustrated by several strong correspondences between BrainMap (sub)domains and NeuroSynth terms (**Supplementary Figures 13, 14**). Furthermore, in NeuroSynth several behaviors reached convergence despite related (sub)domains not doing so in BrainMap (e.g., ‘Language’ vs. ‘Language Phonology’; ‘Rhythm’ vs. ‘Music’; and parts of the Working Memory representation now showing). In some, albeit fewer, cases BrainMap (sub)domains reached convergence when related NeuroSynth terms did not (i.e., ‘Action Observation’, ‘Sadness’). Although we forego detailed discussion of NeuroSynth maps, they generally correspond with current consensus on cerebellar functional topography (King et al., 2019; Nettekoven et al., 2024; Saadon-Grosman et al., 2024; Stoodley and Schmahmann, 2018, 2010; Xue et al., 2020). This replication suggests that – with large enough data – noise, crudeness of analysis (including only peak coordinates), and even a lack of manual article scanning can be overcome to provide generalizable cerebellar maps. The replication of BrainMap (sub)domain maps, especially ones that did not reach significant convergence, strengthens the reliability of our findings by ensuring they were not driven by a particular dataset.

### 3.3 C-SALE complements the cerebellar behavioral topography: a literature review

Here, we provide a brief interpretation of the BrainMap localizations in the form of a brief literature review. Since we cannot discuss all findings, we focus on the significant behavioral domains in BrainMap, considering NeuroSynth findings when they are relevant. Note that because our method aims to provide a summary of the mappings of individual studies, we do not compare our maps with selected literature findings. Instead, we discuss them in terms of what is generally known (or unknown) about cerebellar behavioral localizations. Hence, emphasis is placed on mapping to widely used cerebellar parcellations. We also focus on the potential (clinical) utility of these mapping insights.

#### 3.3.1 Action

The cerebellum plays canonical roles (Luciani, 1893) in motor execution, sequencing, learning, observation, and preparation (Doya, 1999; Doyon et al., 2002; Errante and Fogassi, 2020; Hull, 2020; Kohler et al., 2023; Manto et al., 2012; Shahshahani et al., 2024; Thach, 1998). In our study, ‘Action’ including subdomains ‘Execution’, ‘Speech Execution’, and ‘Observation’ converged in cerebellar subregions. Mappings corresponded with established motor representations spanning anterior regions of the paravermis bilaterally (primarily V-VI) (Adrian, 1944; Buckner et al., 2011; Guell et al., 2018a). ‘Action’ and ‘Execution’, mirrored the well-known dual representation, including a right posterior (and VIIb-VIIIb) cluster. ‘Action’, ‘Coordination’, and ‘Movement’ terms in NeuroSynth supported BrainMap localizations. Together, we did not find the posterior representation consistently, and the left posterior motor area was found in none of the motor-related analyses. This suggests that either the number of foci may have been too low to provide confident localizations, or that the posterior motor representation is activated less consistently. Likely it is a combination of both. Unthresholded maps for all motor-related maps across both databases did show high ALE-scores for the full motor representation. ‘Speech Execution’ (BrainMap) and ‘Speech Production’ (NeuroSynth) produced clusters in bilateral lobules V and VI, consistent with reports on bilateral involvement of the cerebellum in the motor aspect of language (Mariën et al., 2014).

‘Action’, ‘Execution’ and ‘Speech Execution’ were most stable in our subsampling analyses, indicating their localizations were highly consistent across the literature. Supporting, all three maps corresponded to motor-related maps of established atlases (King et al., 2019; Nettekoven et al., 2024). The whole brain-connectivity network of these areas also largely overlapped the brain-wide SMN network across both the cerebellum (Buckner et al., 2011; Ji et al., 2019) and cerebrum (Ji et al., 2019). They also notably overlapped other networks, such as the VAN and DAN in the cerebellum (Buckner et al., 2011) and CON in both cerebellum and cerebrum (Ji et al., 2019). ‘Action Observation’ instead converged in left VIIb-VIIIa. Its C-SALE map correlated significantly to MDTB Action Observation and Spatial Maps tasks (King et al., 2019), and attention-related parcels across atlases (Buckner et al., 2011; Ji et al., 2019; Nettekoven et al., 2024). Together, our results support highly consistent mapping of action and motor-related behavior in the cerebellum across the literature.

Motor deficits in cerebellar ataxias (Klockgether et al., 2019), neuropsychiatric (Gowen and Miall, 2007) and neurological (Chen et al., 2023; Gellersen et al., 2021) disorders are well-described and have even served as template for cerebellar non-motor functions (Leiner et al., 1987, 1986; Schmahmann, 1996; Thach, 1998, 1996). Lesion-mapping in cerebellar stroke patients shows the impact of the location of cerebellar damage: anterior lesions primarily cause motor deficits, whereas posterior lesions relate to the cerebellar cognitive affective syndrome (CCAS) (Schmahmann, 2004; Stoodley et al., 2016). Ultimately, our accurate localization of motor-related behaviors to specific cerebellar subregions can increase the understanding of cerebellar (dys)function in case of anatomical cerebellar disorder (Reumers et al., 2025).

#### 3.3.2 Cognition

Within ‘Cognition’, we found convergence in the subdomains ‘Language’, including ‘Language – Speech’ and ‘Language – Semantics’, ‘Working Memory’, and ‘Social Cognition’. Consensus papers and large-scale meta-analyses underline the cerebellar role within the socio-cognitive domain (Koziol et al., 2014; Van Overwalle et al., 2020b, 2020a, 2014). The ‘Cognition’ construct itself may be somewhat too unspecific to assign it well to cerebellar subregions through the meta-analytic method. This is because wide-ranging cognitive behaviors (e.g., ‘Language’, ‘Social Cognition’, and ‘Working Memory’) may activate distant cerebellar areas. Notwithstanding, we found a small area of convergence for ‘Cognition’ in right Crus I of the cerebellar posterior-lateral lobe. This implies that this region is consistently activated across the breadth of cognitive behaviors. Crura I and II have been widely regarded as the primary cognitive region of the cerebellum across evolutionary (Balsters et al., 2014, 2013; Magielse et al., 2022; Ramnani, 2006), developmental (Badura et al., 2018; Gaiser et al., 2024; Manoli et al., 2025), and clinical (Beuriat et al., 2022, 2020; Reumers et al., 2025; Stoodley et al., 2016) scales. Across cerebellar parcellations, this region is consistently associated with socio-linguistic and multi-demand functions (King et al., 2019; Nettekoven et al., 2024). These lobes likely gain their function in part due to their extensive connectivity with other areas within the DMN and FPN networks (Marek et al., 2018; Palesi et al., 2017, 2015; Xue et al., 2020).

##### 3.3.2.1 Language

Language’, like ‘Cognition’, captures diverse behaviors: from ‘Speech Execution’ to ‘Semantics’ and ‘Orthography’. In BrainMap, we found ‘Language’ convergence in left VI. This is interesting, as language is commonly believed to reside primarily in the right Crura I-II and posterior parts of lobule VI. Connectivity between these right-sided posterolateral cerebellar areas and left-dominant language areas of the cerebral cortex have been established structurally (Jobson et al., 2022; Kelly and Strick, 2003; Palesi et al., 2017, 2015) and functionally (Guell et al., 2018a; Marek et al., 2018; Xue et al., 2020). Within BrainMap the right postero-lateral cerebellum displayed high ALE scores but did not reach convergence, whereas the ‘Language’ and ‘Communication’ terms in NeuroSynth did. Moreover, ‘Language’ in NeuroSynth correlated strongly (r = .57, *p*_variogram, FDR_ < .001) with ‘Language – Phonology’ in BrainMap. These findings strengthen the notion that cerebellar language functions may consistently activate posterior-lateral regions of the right cerebellum. However, lesion evidence suggests that language in the cerebellum may indeed be bilaterally organized (Murdoch and Whelan, 2007). Cerebellar language experts could not reach consensus on this topic (Mariën et al., 2014). However, our results support the notion of at least some left-sided cerebellar involvement in language. This corresponds well with what is known from parcellations, as illustrated by several significant correlations between language C-SALE maps and language-related parcels (King et al., 2019; Nettekoven et al., 2024).

Language production and comprehension are known deficits in cerebellar degenerative disorders (Reumers et al., 2024). Bilateral cerebellar strokes are connected to deficits in semantics, syntax, word retrieval, and phonology. Cerebello-subcortical-cerebral networks likely support broad cerebellar roles across language (Satoer et al., 2024). Unfortunately, we were not able to perform MACM for ‘Language’, which represents a target for follow-up.

##### 3.3.2.2 Working Memory

‘Working Memory’ primarily occupied the right inferior VI-Crus I. This map corresponded significantly to memory aspects of parcellations including Object 2-back and Math activation maps (King et al., 2019). It also overlapped with the FPN (Buckner et al., 2011; Ji et al., 2019). Like our study, previous meta-analyses (Emch et al., 2019; Stoodley and Schmahmann, 2009) support mostly right-sided working memory representations. However, our replication in NeuroSynth ‘Working Memory’ shows a bilateral representation in VI-Crus I. Our subsampling analysis in BrainMap also shows convergence in the left cerebellum for some subsamples. The symmetry of ‘Memory’ and ‘Working memory’ unthresholded z-maps in BrainMap provide further support for the existence of a left-sided localization. This is consistent with a large-scale neuroimaging study (Guell et al., 2018a). Together, our combined BrainMap and NeuroSynth maps pertaining to memory function reach significant convergence across posterior-lateral and far-posterior parts of the cerebellum, coinciding with (Guell et al., 2018a).

Memory deficits are common in cerebellar degeneration (Reumers et al., 2024). The cerebellum has also increasingly been implicated in dementia (Gellersen et al., 2021; Li et al., 2023) and healthy aging (Arleo et al., 2023; Bernard et al., 2023; Romero et al., 2021), both characterized by decreasing working memory performance. Although ‘Explicit Memory’ did not converge, cerebellar deficits are implicated causally in development of episodic memory across aging (Almeida et al., 2023). ‘Episodic Memory’ in NeuroSynth did converge, signaling this topic warrants attention in future work.

##### 3.3.2.3 Social Cognition

We report highly symmetric convergence of ‘Social Cognition’ in bilateral Crura I-II (Guell et al., 2018a; Manoli et al., 2025), which was mirrored by NeuroSynth terms ‘Belief’, ‘Empathy’, and ‘Social Cognition’. Cerebellar involvement in socio-cognitive functions is now well established (Van Overwalle et al., 2020b). ‘Social Cognition’ appeared to take up a hub-like position within the cerebellum. It correlated strongly with many MDTB tasks (King et al., 2019), S2 and S3 of the socio-linguistic network (Nettekoven et al., 2024), mapped to the DMN (Buckner et al., 2011; Ji et al., 2019), and to anchors of the first two functional cerebellar gradients (Guell et al., 2018b). Clusters overlap established regions of Theory of Mind activations in adults (Haihambo et al., 2023; Manoli et al., 2025; Metoki et al., 2022; Pu et al., 2020; Van Overwalle et al., 2022, 2020a, 2014). Together, social cognition appears to be localized highly consistently across studies.

Both typically developing (Manoli et al., 2025) and pediatric populations (H. Olson et al., 2023) have shown selective involvement of Crura I-II in the undisrupted development of social cognition (I. Olson et al., 2023). In mice, social behavior was adversely affected after developmental damage to particularly these areas (Badura et al., 2018). Reports of social and cognitive deficits related to cerebellar damage (Badura et al., 2018; Chao et al., 2023; I. Olson et al., 2023; Reumers et al., 2024) and alterations in neurodevelopmental and psychiatric disorders (Abram et al., 2024; Bernard and Mittal, 2015; Kong et al., 2024; Pinheiro et al., 2021; Stoodley, 2014) have become commonplace. The cerebellum may cause these effects through widespread (dys)connectivity. For example, MACM places the cerebellum within an ASD alteration network alongside the amygdala and fusiform gyrus, associating it with language semantics and action observation. Both relate to socio-cognitive alterations in ASD (Goodwill et al., 2023).

##### 3.3.2.4 Conclusion for Cognition

Concluding, it appears that ‘Language’, ‘Working Memory’, and ‘Social Cognition’ all map to hub-like cerebellar areas, characterized by high cerebellar convergence of cerebral inputs^80^. This is suggestive of cerebellar information integration within these behaviors. Recent work using cerebellar stimulation assessed the propagation of modular network structures, supporting such an integrative role across cerebello-cerebral networks (Bansal et al., 2024), and pushing back against the idea that the cerebellum only mirrors functions of connected brain areas (Diedrichsen et al., 2019; Orban de Xivry and Diedrichsen, 2024; Wang et al., 2013). Moreover, meta-analyses of cerebellar stimulation studies show that manipulation of cerebellar activity can directly alter behavior (Gatti et al., 2021; Oldrati and Schutter, 2018; Pezzetta et al., 2024). This underlines the translational relevance of cerebellar functional localizations.

#### 3.3.3 Emotion and Interoception

Emotion is thought to be organized diffusely across the brain, and plays important regulatory roles across human behavior (Adamaszek et al., 2017). By focusing specifically on the cerebellum, several studies have found involvement of both the vermis and hemispheres in emotional behaviors including reward (anticipation and outcome) (Kruithof et al., 2023), (reactive) aggression and impulsivity (Wolfs et al., 2023c, 2023b, 2023a) and violent behavior (Klaus et al., 2024), among many others (e.g., happiness, sadness, anger, disgust, and fear (e.g. (Baumann and Mattingley, 2012), see also (Adamaszek et al., 2017)). Clinical trials of cerebellar stimulation within the emotional domain, such as in anger (Kruithof et al., 2024), threat (Kruithof et al., 2025), or fear (extinction) (Thieme et al., 2025) are already underway.

The diffuse z-maps across ‘Emotion’ subdomains suggest that individual functional studies of emotion have continuously implicated different vermal and paravermal regions in these behaviors. ‘Sadness’ and ‘Negative Emotion’ were the only affective subdomains to converge, in right VI-Crus I (Baumann and Mattingley, 2012) and right VI, respectively. These generalizable clusters may be good targets for neurostimulation in the future. Lastly, it has become appreciated that interoceptive-emotional behaviors may map together in the brain salience network (Seeley, 2019; Seeley et al., 2007). For example, pain (Moulton et al., 2010) and its emotional regulation (Adamaszek et al., 2017; Moulton et al., 2011) both activate the cerebellar hemispheres. In BrainMap, ‘Interoception’ and its subdomains did not reach convergence. It must be noted that ‘Emotion’ (bilateral Crura I-II), ‘Arousal’ (right VIIIA-B) and ‘Pain’ (right paravermal I-III) did converge in NeuroSynth. Unthresholded maps across both datasets revealed rather diffuse patterns across vermal, paravermal, and hemispheric regions. CBMAs may be less appropriate for mapping behaviors spread diffusely across the cerebellum (or brain).

#### 3.3.4 Perception

Finally, we considered ‘Perception’, where the cerebellum is believed to mismatch sensorimotor expectation and feedback through internal forward models (Ishikawa et al., 2016; Tanaka et al., 2020; Wolpert et al., 1998). Additionally, the cerebellum is increasingly discussed in outward senses such as audition (Petacchi et al., 2005) and vision (Baumann and Mattingley, 2012, 2010). Crura I is specifically implied in increased perceptual demand from auditory and visual motion (Baumann and Mattingley, 2010). Cerebellar structural and functional connections with cerebral perceptual cortices, cerebellar activations, and behavioral alterations in case of cerebellar damage, argue firmly for a cerebellar role in perceptual processes (Baumann et al., 2015).

Caution is advised in interpreting convergence across ‘Vision’ (including ‘Shape’). Inspection of these data suggest that these clusters may be cerebral in origin, as most of their cluster mass mapped to the occipital cortex. This makes sense, as the parts of the (anterior) cerebellum where we find the convergence directly neighbor occipital and temporal cortices. We noted the opposite side of the same issue in our MACM, where cerebellar clusters bled over into the fusiform cortex. Explicit cerebellar isolation is often used to overcome this BOLD-bleeding (e.g., (Manoli et al., 2025)). However, substantial spatial misalignment across the many studies we included, prevents clean isolation of the cerebellum. Fortunately, ‘Somesthesis’, and ‘Vision – Motion’ converged more medially, making it unlikely that cerebral signals caused these clusters. ‘Vision – Motion’ converged in right paravermal VI. Not much is known about the mapping of somesthesis within the cerebellum. Future work may detail cerebellar involvement in the outward senses, especially as they are often absent from cerebellar parcellations (e.g., (Buckner et al., 2011; King et al., 2019; Nettekoven et al., 2024)).

### 3.4 Stability of cerebellar mapping differs across behavioral domains

Next, we assessed the stability of C-SALE maps. ‘Action’ was the most stable BD across different subsampling strategies: parts of the anterior representation reached convergence across all subsamples, and the posterior representation reached convergence across most. In line with our findings, cerebellar parcellations have repeatedly placed motor representations in similar locations (Adrian, 1944; Buckner et al., 2011; Ji et al., 2019; King et al., 2019; Nettekoven et al., 2024), as have task-based localizations (Guell et al., 2018a) and precision-mapping approaches (Boillat et al., 2020; Marek et al., 2018; Xue et al., 2020). Differences in stability of meta-analytic maps could be explained by how behaviors are organized in the cerebellum and the rest of the brain. Cerebellar functional connectivity is organized along a unimodal-transmodal axis (Guell et al., 2018b; Katsumi et al., 2023; Liu et al., 2022), as are cerebral (Margulies et al., 2016; Paquola et al., 2019) and whole-brain (Katsumi et al., 2023) connectivity. Gradients of functional abstraction (D’Mello et al., 2020), transcriptomic and molecular expression (King et al., 2025; Wang et al., 2024), and granule cell physiology (Straub et al., 2020) underline gradual organizational aspects of the cerebellar cortex. Our C-SALE clusters for aspects of language and memory, that are topographically organized into adjacent areas of the right posterior-lateral cerebellar cortex, provide further support for this paradigm of gradual cerebellar organization. Within this framework, associative behaviors may elicit activity across small, distributed cerebellar areas, relative to more unimodal and localized connectivity patterns of the motor brain network (King et al., 2023; Paquola et al., 2019). Each set of afferents may be part of relatively separate reciprocal networks involved in distinct functions (see: cerebellar modules (Apps et al., 2018; Cerminara et al., 2015; Cerminara and Apps, 2011)). Even adjacent modules can be involved in different functions, making it difficult to expose their functions using CBMAs. Because CBMAs excel at localizing behavioral clusters at the mesoscale (due to their summarizing nature), complementary precision mapping approaches are needed to zoom into specific behaviors, and reveal closely juxtaposed cerebellar organization (Boillat et al., 2020; Saadon-Grosman et al., 2024).

‘Emotion’ and ‘Interoception’, with seemingly low regional preference, are good examples of domains where overcoming the issues of partial volume effects in the small, folded, often misaligned (Diedrichsen et al., 2009) cerebellum are especially important. Even if these task domains elicit consistently elevated activity across small distributed or juxtaposed cerebellar areas, the localizing nature of CBMAs, combined with these cerebellar challenges, may more often preclude finding statistical convergence. On the other hand, greater experimental consistency in e.g., ‘Action’, ‘Working Memory’, and ‘Vision’ may also lead to increased meta-analytic stability in those behaviors. Concretely, whereas some experiments (such as those examining motor functions) may more consistently elicit activations, other more sophisticated behavioral experiments (such as cognitive reasoning tasks) may be more variable in both experimental setup and participant response. Together, low stability and differences between our C-SALE mappings and previous CBMA clusters warn for caution in interpretation of small-to-intermediate-sized CBMAs. Nevertheless, it is reassuring that despite these limitations, the improved C-SALE method was able to identify generalizable cerebellar clusters in many behaviors (fifty-four of 138 analyses) across the literature.

### 3.5 Correspondence of C-SALE maps with cerebellar parcellations and whole-brain connectivity

Spatial correspondence to previous parcellations and mappings (Buckner et al., 2011; Guell et al., 2018b; Ji et al., 2019; King et al., 2019; Nettekoven et al., 2024) revealed that converging clusters occasionally mapped significantly to one or several existing parcels or maps. Locations of subthreshold convergence also sometimes colocalized significantly with (behaviorally) related parcels and maps. Specifically, for the MDTB maps (King et al., 2019), we observed two clusters globally separating association (sub)domains, such as ‘Cognition’ and ‘Emotion’, from sensorimotor (sub)domains, such as ‘Action’ and ‘Perception’. Notably, ‘Reasoning’, ‘Semantics’, and especially ‘Social Cognition’ mapped to a wide variety of association tasks in the MDTB, whereas ‘Action’ and ‘Execution’ demonstrated very high correlations with the MDTB finger sequence task. Similarly, in the mid-granularity hierarchical atlas (Nettekoven et al., 2024), ‘Social Cognition’ mapped to posterior sociolinguistic parcels, whereas ‘Action’ and ‘Execution’ were strongly associated with anterior sensorimotor parcels.

We report a high degree of correspondence between C-SALE maps and behaviorally related aspects of cerebellar mappings and atlases (Buckner et al., 2011; Ji et al., 2019; King et al., 2019; Nettekoven et al., 2024), and between MACM maps and subcortical parcellations and cerebral networks (García-Cabezas et al., 2019; Ji et al., 2019; Tian et al., 2020). However, many behavioral (sub)domains (e.g., those within ‘Emotion’ and ‘Interoception’) mapped only moderately or not at all to previous parcellations. This implies that these (sub)domains may thus far not be as well represented by current state-of-the-art parcellations as other behaviors are. Incongruencies with previous parcellations and mappings, imperfect correspondences between BrainMap and Neurosynth, and a lack of subsample stability together show that complementary perspectives are necessary to fully understand cerebellar behavioral topography. Our C-SALE maps may thus complement rather than replace other maps and parcellations, since they contain a different type of information. Whereas parcellations locate cerebellar borders for functional communities, behavioral mappings such as our meta-analytic maps identify activation clusters specific to individual behaviors. Even though the discriminatory performance of their borders makes parcellations useful for many applications, our findings suggest the need for a more comprehensive behavioral cerebellar topography. This would ideally be created directly from full task-activation maps (not just peak foci, as already achieved in parcellation and mapping studies), span many task domains (as in CBMAs), and include many – diverse (e.g., non-*White, Educated, Industrialized, Rich, Democratic or WEIRD)* (Henrich et al., 2010)) – individuals. Both the functional fusion framework (Nettekoven et al., 2024; Zhi et al., 2023) and neuroimaging mega-analyses are promising avenues to reach this goal. For summarizing methods, extending coordinate-based with image-based meta-analyses (while still accounting for biases) may be the way forward (Salimi-Khorshidi et al., 2009). To reach this goal, it is important that researchers publish their unthresholded statistical maps on open science databases such as NeuroVault (neurovault.org/) (Gorgolewski et al., 2015).

Lastly, we performed MACM to reveal coactivation networks between behavior-specific cerebellar subregions and the rest of the brain for several aspects of ‘Action’, as well as ‘Working Memory’ and ‘Vision’. These maps correspond largely with current knowledge about where these functions reside in the brain. The ‘Action’ and ‘Execution’ MACMs corresponded with the brain-wide motor network (Buckner et al., 2011; Ji et al., 2019; Xue et al., 2020). Action-related clusters in the cerebellum were coactivated with the cerebral motor cortex and subcortical basal ganglia. These areas are involved in established structural pathways with the cerebellum across the primate lineage (Bostan et al., 2013, 2010; Bostan and Strick, 2018; Caligiore et al., 2017; Magielse et al., 2022). Similarly, the small meta-analytic clusters in ‘Vision’ were associated with known visual regions in the occipital cortex (e.g., ref. (Brewer et al., 2005)). Unfortunately, we could only perform valid MACM analyses in five (sub)domains, due to insufficient sample sizes (a minimum of seventeen experiments is recommended (Eickhoff et al., 2016)). This leaves MACM analyses in other (sub)domains as a promising avenue.

### 3.6 Limitations and future directions

Several limitations of this study are important to consider. Typically, CBMAs start with a literature search, followed by manual text scanning to homogenize experiments (Laird et al., 2009; Müller et al., 2018; Tahmasian et al., 2019). Since this was not feasible due to the vast literature of studies reporting activations in the cerebellum, we focused on a data-driven interpretation of BrainMap data. The manual curation in BrainMap likely lowered the number of wrong categorizations of experiments. However, as suggested above, it is still possible that there is variability across experiments within BrainMap itself. For example, a language task may require different extents of motor behavior, such as tongue or hand movements, altering cerebellar activity. We could not manually monitor such experimental consistency within (sub)domains. Each analysis may thus include experiments probing subtly different behavioral aspects and extents of overlap with other behaviors. Therefore, we could not fully disentangle experimental conditions from cerebellar organization. However, the fact that most of our meta-analytic maps were consistent across both the BrainMap and the larger – and noisier – NeuroSynth database increases our confidence in the biobehavioral validity of our results. The fact that despite the crude nature of CBMA analysis (only peak coordinates are available), cerebellar clusters can be localized within abundant noise, suggests that the true biological signal in these subregions is highly generalizable.

Although we provide many meta-analytic maps, we are not able to discuss each in detail. There is still room to improve these mappings by zooming in and improving or further specifying behavioral categorizations. Consequently, every result (both in BrainMap and NeuroSynth) is an opportunity for more specific research questions aiming to map behaviors more precisely. Additionally, CBMAs may be performed at even finer granularities i.e., individual experiment types or within specific sub-populations.

For the present study, the low resolutions of older data in meta-analytic databases may ultimately limit the accuracy of localizations. Scanning at 7T can greatly improve cerebellar imaging resolutions, as can using cerebellum-optimized sequences (Priovoulos et al., 2023; Priovoulos and Bazin, 2023) and dielectric pads (Vaidya et al., 2018). Although we may be close to limits of CBMAs in the cerebellum with current databases, increased resolutions will further improve localizations. High-resolution cerebellar data in many individuals can also help improve cerebellar alignment (Diedrichsen et al., 2009) across studies, which is essential given the summarizing premise of CBMAs.

Overall, our findings contribute to an ever-expanding framework for understanding cerebellar involvement across various cognitive domains. Our specific contribution is the largest synthesis of cerebellar literature to date, that accounts for biases inherent in previous mapping efforts. As a result, our findings – particularly those highlighting behavior-subregion associations consistent with previous cerebellar mapping efforts – can inform future research by identifying regions of consistent cerebellar involvement across behaviors and studies. This understanding may enhance insights into cerebellar subregional involvement in normal and abnormal development and suggest potential targets for neurostimulation. For example, previous research has linked abnormalities in the social regions of the posterior cerebellum to the onset of neurodevelopmental disorders such as autism (Riva et al., 2013). Other research lines have targeted cerebellar regions encompassing sensorimotor and associative functions, such as language and memory, for non-invasive neurostimulation (Grimaldi et al., 2014). Our findings offer a generalizable framework for creating behavioral ROIs that could aid in diagnosing disorders and alleviating symptoms within specific cognitive domains.

However, to truly be generalizable, the included data should represent a more appropriate sample of the world’s population (Henrich et al., 2010). To support important endeavors in this direction, our framework is fully adaptable to other neuroimaging databases and brain regions. This can help refine dataset– or area-specific functional topographies as more neuroimaging data becomes available worldwide. Moving forward, incorporating this CBMA framework into broader neuroscientific research could enhance our understanding of brain-behavior relationships and inform future studies on cerebellar function in health and disease.

## 4. Conclusion

In the current work, we used the BrainMap database and an adaptation of the ALE method (C-SALE) to perform large-scale CBMAs across thirty-seven behavioral (sub)domains. Our findings highlight the systematic omission of the inferior cerebellum in neuroimaging data. This suggests that caution should be exercised when interpreting previous meta-analytic localizations, which may have been inaccurate or overly generous. C-SALE corrects cerebellar reporting biases, improving CBMA accuracy, even without full cerebellar coverage. Our new method is openly available and can be adapted to any dataset and brain volumetric ROI to improve behavioral localizations in the future. Across two experiment databases, we showed that ‘Action’, ‘Cognition’, ‘Emotion’, and ‘Perception’ behaviors converge onto distinct cerebellar subregions, further strengthening and expanding on previous cerebellar mappings and parcellations efforts. Overall, we provide a comprehensive meta-analytic cerebellar topography, emphasizing the cerebellum’s broad involvement in human behavior. While a consensus about the cerebellum’s involvement in a wide range of human behaviors has long been reached within the cerebellar community, we hope that the statistical consensus presented here will convince any remaining sceptics.

## 5. Methods

### 5.1 Overview

In this study, we leveraged the BrainMap database (Laird et al., 2005b) as our main dataset, to pull the largest available sample of manually indexed fMRI and PET literature at the whole-brain-level. In BrainMap, experiments were comprehensively and manually labeled according to sample and study characteristics, including nested BD annotations and sample sizes. This curation provided us with higher confidence in behavioral categorizations, and in turn in the meaning of cerebellar localizations and the validity of the C-SALE method. Hence, most descriptions in the following deal with BrainMap. Nevertheless, we validate our method and replicate our findings in Neurosynth (Yarkoni et al., 2011) (see *section 5.12*). In this second database, experiments were automatically categorized according to the occurrence of specific terms in associated article texts, which meant potentially less specific labels. For example, it was unclear how many of the studies mapping to the ‘action’ term involved action behavior, rather than simply containing the word ‘action’ in their article texts. However, NeuroSynth contained substantially more experiments, which allowed us to validate our method and resulted in an even more comprehensive literature sample. Moreover, the NeuroSynth analysis helped to capture the influence of automated study selection on behavioral mapping accuracy. Comprehensive access to BrainMap data and metadata was authorized by a collaborative use agreement (brainmap.org/collaborations). From the database, we initially obtained data from 7,771 whole-brain experiments (2,204 studies; 32,530 participants). Of these, 2,322 experiments (1,109 studies; 16,159 participants) reported peak coordinates within an expanded cerebellar mask (see *section 5.2*). After merging experiments within each study (Müller et al., 2018), 1,109 unique experiments remained. From these, we constructed C-SALE maps across five BDs (‘Action’, ‘Cognition’, ‘Emotion’, ‘Interoception’, and ‘Perception’) and thirty-two more fine-grained behavioral subdomains. These included several aspects of language and memory, fear and reward, and most of the outward senses. To recognize cerebellar clusters in relation to the whole-brain, we subsequently performed MACM. Both findings at the cerebellar (C-SALE) and whole-brain (MACM) level were compared to existing parcellations. For C-SALE results, we also assessed stability of cerebellar localizations across behavioral (sub)domains.

### 5.2 Study Inclusion Strategy

Experiments were included in C-SALE analyses based on the following criteria: **1.** To construct a functional map of the general healthy population, only normal mapping studies that measured within-subject activation contrasts in healthy controls (no interventions; number of subjects eight to forty-four) using fMRI or PET were included **2.** To localize behaviors to cerebellar subareas, only experiments with a peak coordinate within the cerebellar ROI were included. The cerebellar ROI was isolated using the 1 mm resolution cerebellar template (Diedrichsen et al., 2009), dilated 6 mm in all directions (hereafter “dilated cerebellar mask”) to account for spatial misalignment across studies. This step also aimed to overcome cerebral cortical BOLD-bleeding, by accounting for these signals in the biased baseline distribution. **3a.** Separate datasets were created for the five BDs indexed in BrainMap. Only activation contrasts were considered, as deactivations often had insufficient numbers of experiments. Activation foci were limited to those within the mask in **2**. and those included within each BD. **3b.** Next, in the same way, datasets were created for subdomains indexed beneath the five BDs. **4.** To prevent unjustly embellishing statistical power, it is important to prevent overlaps in included experiments (Müller et al., 2018). Within the BrainMap database, single studies may contain multiple experiments with potential overlaps in experimental contrast and participants. Hence, we merged the coordinates within a study to represent a single experiment. This validated approach (Vanasse et al., 2018) minimizes within-group and within-experiment effects (Turkeltaub et al., 2012), preventing overlapping experimental contrasts excessively contributing to convergence (Müller et al., 2018). As empirical simulation indicates that seventeen or more experiments should be included for stable meta-analysis (Eickhoff et al., 2016), ultimately five eligible BD and thirty-two eligible subdomain (N_experiments_ ≥ 17) datasets were created.

### 5.3 Construction of behavioral cerebellar maps with Cerebellum-Specific ALE

After collecting datasets for each sub(domain), we performed ALE for each using an in-house modification of the Neuroimaging Meta-Analysis Research Environment (NiMARE) version 0.2.0 (Salo et al., 2023a, 2023b). NiMARE is a Python package that facilitates programmatic interaction with BrainMap activation foci and performing CBMA based on, among other algorithms, ALE (Eickhoff et al., 2012, 2009). We added an efficient implementation of ALE using graphical processing units (GPUs) (github.com/amnsbr/nimare-gpu). This facilitated running the calculations of many permutations of ALE, and the multiple experiments therein, in parallel. These permutations are necessary to generate the null samples for each meta-analysis, as described below. In the GPU implementation, calculations pertaining to individual permutations, experiments, and foci were parallelized at two levels, across GPU “blocks” (permutations and experiments) and “threads” (foci). The GPU used for the main analyses was of the Nvidia GeForce GTX 1080 Ti model. Ultimately, parallelization was highly advantageous because of the high number of performed meta-analyses across different sub(domains) and subsample configurations. The GPU implementation led to considerable speed-ups of calculations when scaling the number of permutations and experiments. For example, we observed a speed-up of 101.3x in GPU (Nvidia Tesla P100) versus central processing units (CPUs) (AMD EPYC 7601, using a single core) when running a rather typical SALE analysis for 100 experiments with 10,000 permutations (**Supplementary Figure 23**).

Essentially, ALE tests the distribution of experimental activation foci against a null distribution that assumes random spatial associations across cerebellar GM. Put simply, it finds spatial locations where activations converge significantly more than chance. However, visualizing foci, we observed strong spatial biases in reported effects (**Figure 1a, Supplementary Figure 2)**. These were very closely replicated in the NeuroSynth database (**Supplementary Figure 2a**). Together, the non-random distribution of foci rendered the standard null hypothesis of ALE (**Figure 1b)** unsuitable (Eickhoff et al., 2012, 2009). Hence, we aimed to account for biases by incorporating the distribution of foci into the null model against which datasets were tested. This approach is comparable to the SCALE approach used for MACM (Langner et al., 2014). Since we here adapt this approach to the cerebellum, we refer to it as *cerebellum-specific activation likelihood estimation* (C-SALE). Note that the method can be flexibly adapted to any (volumetric) brain region. To assess improvements of C-SALE (described later), we compared its mappings results for BDs and subdomains with classic ALE results. The methods for classic ALE are discussed here first. For a full illustration of the commonalities and differences between classic ALE and C-SALE, see **Figure 7**.

**Figure 7:**
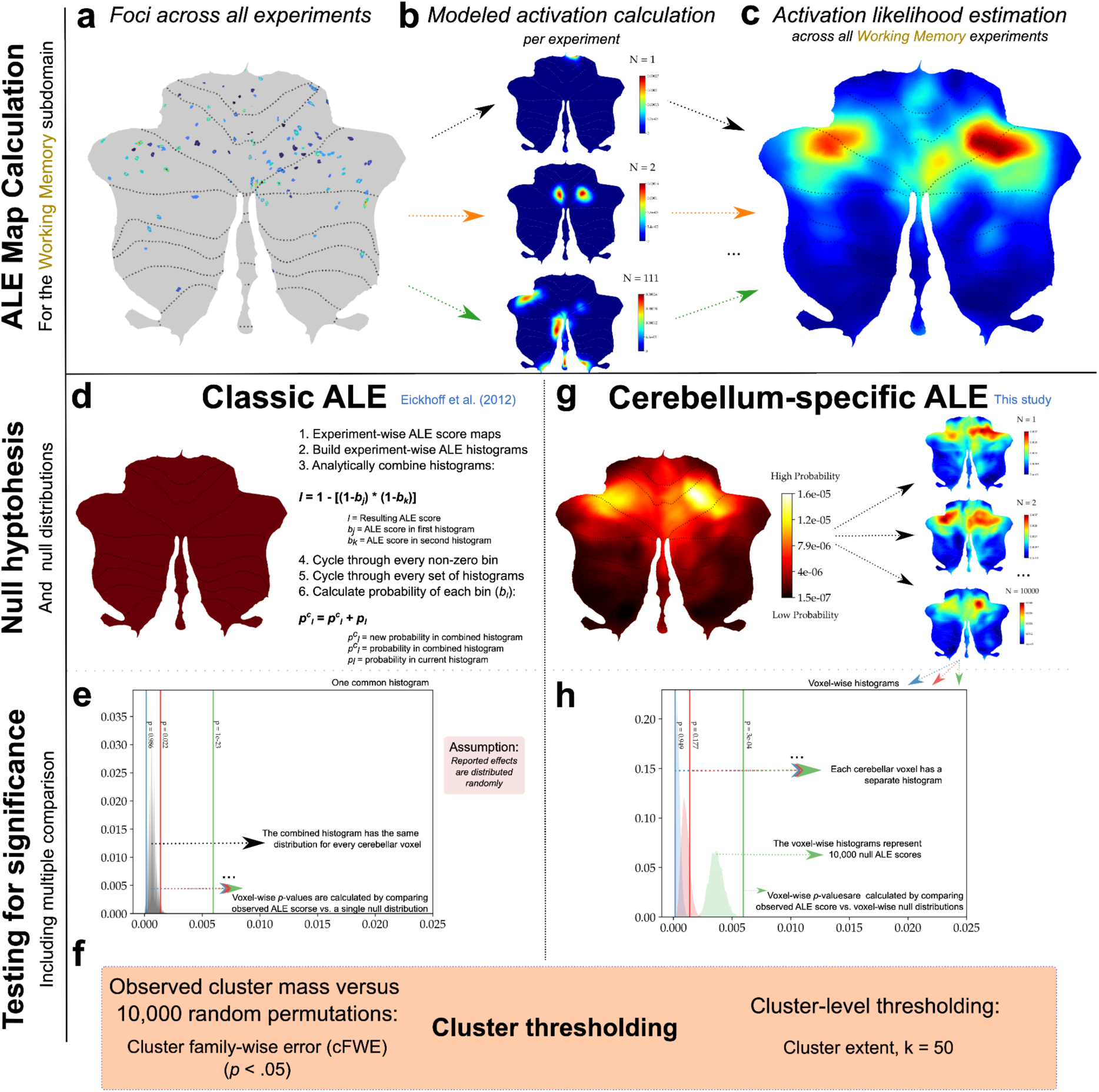
Commonalities and differences across classic ALE and cerebellum-specific ALE methods. (**a-c**) ALE score map calculation is the same across classic ALE and C-SALE. Working Memory is used here as an example. (**a**) First, peak activation coordinates (foci) across a behavioral domain or term are collected. (**b**) Then, for each experiment separately, foci are convolved with Gaussian kernels with their full width at half maximum inversely proportional to the experiment’s sample size. The convolved foci maps are combined into modeled activation (MA) maps by taking their voxel-wise union. (**c**) Last, experiment-wise MA maps (N = 111 in this example) are combined into an ALE score map, by again taking their voxel-wise union. Classic ALE and C-SALE diverge due to different null hypotheses (**d, g**). (**d)** In classic ALE (**d,e,f**), the ALE score map (**a-c**) is tested against the null hypothesis that foci are distributed randomly. The current implementation of ALE (Eickhoff et al., 2012) analytically combines experiment-wise ALE score histograms (**d**). (**e**) Ultimately, this leads to one common histogram of ALE scores, against which each voxel’s ALE score (**a-c**) is compared to obtain uncorrected *p*-values, *p_voxel_*. (**f**) Clusters are defined based on *p_voxel_ < .001*, and a permutation-based strategy (cluster family-wise error; cFWE) is used to determine how observed cluster masses compared to cluster masses in 10,000 randomly generated maps. Clusters are considered significant if they exceed *p_cFWE_* < .05. In C-SALE (**f,g,h**), instead of assuming spatial homogeneity of reported effects, unequal spatial distributions are assumed (**g**). The null hypothesis describes the base likelihood of finding foci at cerebellar voxels (**g**; left). Next, 10,000 biased MA null maps are created by randomly sampling coordinates relative to the probability distribution (**g**; right). This bias is reflected in shifts in ALE score histograms across voxels (**h**). This means that when testing the ALE score map of a behavioral domain (**a-c**), each cerebellar voxel is tested against its unique distribution of ALE values (**h**). As illustrated by the colors, voxels require different ALE scores to reach convergence (*p_voxel_* < .001), depending on their baseline probabilities. (**f**) Lastly, to threshold cluster sizes, we opted for a lower limit of k = 50 voxels to prevent small cerebellar regions reaching convergence.

### 5.4 Classic ALE analysis

The classic implementation of ALE (Eickhoff et al., 2012, 2009) assesses convergence of reported effects (activation foci) against a null hypothesis that assumes foci are distributed equally across the cerebellar GM. For each analysis, we did the following: **1.** Convolve activation foci (**Figure 7a**) by a 3D Gaussian kernel. Importantly, these kernels had a full width at half maximum (FWHM) inversely related to sample size. This means narrower distributions – and thus higher statistical certainty – were assigned to experiments with larger samples (and vice versa). **2.** The convolved foci were then combined, first at the level of individual experiments. The union of all convolved foci within an experiment was taken, creating modeled activation (MA) maps (**Figure 7b**). These maps reflect voxel-wise maxima across convolved foci. **3.** Next, experiments within a BD were combined. This was again done by taking their union, resulting in an ALE score map for each BD (**Figure 7c**). **4.** These ALE score maps were then tested against the null assumption of random spatial associations (**Figure 7d**), resulting in voxel-wise *p*-values (**Figure 7e**). **5.** These *p*-values were thresholded at *p_voxel_* < .001. Using a permutation method, 10,000 random sets of experimental foci were created under the null hypothesis. For each of these coordinate sets, steps **1-5** were repeated. **6**. Lastly, cluster-level family-wise error (cFWE) was used to test statistical significance of resulting clusters from step **5** while accounting for multiple comparisons. Here, cluster mass of the actual experimental set within each sub(domain) was compared against the null distribution of cluster masses in the 10,000 random ALE score maps. Cluster masses were thresholded at *p_cFWE_* < .05 (**Figure 7f**).

### 5.5 Accounting for the spatial bias of reported effects in probabilistic C-SALE

However, as discussed above, there were substantial spatial biases, with many more foci in superior regions. This rendered the standard null hypothesis unsuitable (**Figures 1, 7**). In C-SALE, we thus tested where reported effects converged in cerebellar subregions beyond the baseline, spatially biased, probability distribution (**Figures 1b, 7g**). Here, we outline our method for calculating voxel-wise baseline (null) probabilities in reported effects. Next, we describe how we tested each (sub)domain’s ALE score map against this new null hypothesis.

#### 5.5.1 Creating a bias-accounting null model

To calculate voxel-wise baseline probabilities within the cerebellum, we first created a whole-brain probability map (**Supplementary Figure 4**) by taking all activation and deactivation foci in the BrainMap database (its February 2024 release) for contrasts in healthy subjects (8,408 unique experiments; 2,214 studies; 32,660 participants; 69,703 coordinates). Then, each coordinate was convolved by a 3D Gaussian kernel with its FWHM inversely proportional to the experimental sample size of the coordinate. Next, convolved foci were summed voxel-wise to obtain a whole-brain probability map of reported effects. The resulting map was subsequently masked to the dilated cerebellar mask and normalized to a sum of one across the voxels. This resulted in a baseline probability map of finding foci at any cerebellar voxel, regardless of task domain. By testing behavioral activations against this baseline, C-SALE accounted for literature reporting biases. In this way, all behavioral data could be included, regardless of cerebellar coverage, to produce accurate maps. Constructing a probabilistic, rather than deterministic (as in (Langner et al., 2014)), null model had some advantages. By convolving foci by a FWHM that widens or narrows based on experimental sample size (included in BrainMap (Laird et al., 2005b)), the null model accounts for foci uncertainty. Connectedly, even regions without foci can be sampled into the null model at low probabilities, creating a continuous sampling space. By using the dilated cerebellar mask, the probabilistic null model may also partly account for influences of BOLD-signals from adjacent visual and temporal cerebral cortices. Unequal distributions of reported effects are ubiquitous across the brain (**Supplementary Figure 4**). Hence, by calculating the sum of convolved foci at the brain-level and then restricting this probability distribution to any ROI, the null model can be flexibly adapted to any volumetric brain region for future CBMAs.

### 5.6 Creating meta-analytic maps with C-SALE

Following this calculation of the baseline (null) probability map, we performed C-SALE for each (sub)domain as follows: **1.** The ALE score maps were calculated in the same way as for the classic ALE approach (**Figure 7a-c**). **2.** Next, in contrast to classic ALE, we constructed voxel-wise null distributions for each (sub)domain separately using Monte Carlo permutations (n = 10,000). Specifically, in each permutation, random foci were sampled from the dilated cerebellar mask, weighted by the baseline probability map (**Figure 7g**). Next, a null ALE score map was calculated. **3.** Each voxel’s observed ALE score was tested against its specific null distribution to calculate voxel-wise *p*-values (**Figure 7h**). **4.** The resulting *p*-values were thresholded at *p_voxel_* < .001. Cluster extent was thresholded at k > 50 (**Figure 7f**). Note that we used k = 50 as cluster size threshold, because adapting the preferred method for cluster-extent thresholding, cFWE, is not trivial as it will require an additional layer of permutations on top of permutations used to calculate voxel-wise *p-*values, which was not feasible computationally. Future work may improve cluster thresholding for the C-SALE method.

### 5.7 Comparing performance of C-SALE to classic ALE

We assessed if C-SALE improved the specificity of cerebellar clusters. The main aim of our study was to identify cerebellar subregions with above-chance convergence across behaviors, relative to the cerebellum. Given the premise of this within-cerebellum analysis and the assumption that every cerebellar area has a function, it should be expected that behavioral clusters are spread relatively evenly throughout the cerebellar GM. Thus, first, we compared z-maps resulting from both methods and inspected cerebellar coverage across domains (**Figure 1c, d**). Secondly, we quantified specificity by calculating spatial correlations across (sub)domains. Whereas some spatial overlap is expected, extensive correlations imply the mapping method was not able to differentiate (sub)domains (low specificity), placing them in overly similar cerebellar locations. For every pair of (sub)domains, we calculated spatial correlation (Pearson’s R) of unthresholded z-maps and assessed the correlation coefficients across domains (**Supplementary Figures 6, 7**). To test if the specificity improvement was significant, we performed a paired-sample t-test between the ALE and C-SALE correlation tables (**Figure 1e**). Lastly, to reveal the extent to which both methods were influenced by the biased baseline, we correlated the unthresholded ALE and C-SALE maps with this baseline (**Figure 1f**). It is important to note that baseline reporting rates are calculated across domain-general task findings. In this context, it would be extremely unlikely that all functions across domains are localized to a single area within the cerebellum. Instead, in within-cerebellar CBMA, clusters should be spread evenly. A more likely explanation of excessive spatial correlations with the baseline distribution, would thus be that analyses fail to report for these baseline activation rates.

### 5.8 Assessing stability of cerebellar C-SALE maps

After establishing that C-SALE provided substantial improvements over classic ALE, we investigated C-SALE maps through a repeated subsampling strategy. (Sub)domains contained a range of different numbers of experiments. Since convergence is more likely in larger analyses (Eickhoff et al., 2016), we aimed to assess the stability of different (sub)domain’s C-SALE maps (see **Figures 1-3)** in relation to sample size differences.

Thus, to give complementary perspectives on the (sub)domain maps, we used two subsampling strategies: 1) with subsample sizes fixed to an absolute number of experiments; and 2) with subsample sizes proportional to the (sub)domain. To understand how stability of unthresholded z-maps developed depending on (absolute and proportional) sample sizes, we examined the effects of a range of parameters on BD C-SALE maps. Here, we created subsamples at different absolute (n_subsample_ = 25, 50, 75, 100) and proportional (n_subsample_ = .2 or .4 or .6 or .8 * n_BD_) sample sizes. For subdomains, to minimize the many rather costly computations, we calculated C-SALE maps for subsamples with the arbitrarily selected parameters of n_subsample_ = 50 and (n_subsample_ = .2 * n_subdomain_). One advantage of these parameters was that they facilitated analysis of many (valid) subdomain subsamples. First, this subsampling strategy examined the effect of using the same absolute number of experiments. However, when using an absolute number, the proportion of experiments (from the total sample) is vastly different when examining rather intermediate (e.g., Interoception [n = 68] to large e.g., Cognition [n = 741] samples). Therefore, we also constructed maps at fixed proportions, providing additional perspective on stability that facilitates better comparison across (sub)domains. Subsample analyses were run for all five BDs and the seventeen subdomains with n_experiments_ ≥ 60. For each (sub)domain and subsampling parameter, we created fifty random sets of experiments from the total set of experiments, rerunning C-SALE in each. We then calculated the distribution of spatial correlation coefficients (Pearson’s) across pairs of unthresholded subsampled z-maps within each (sub)domain. To visualize stability of converging (significant) results specifically, we mapped to a common flatmap, per (sub)domain, the proportions of C-SALE subsamples that reached the threshold for significance (*p_voxel_* < .001 and k = 50) at each voxel. This was done for each configuration described above.

### 5.9 Correspondence with previous functional parcellations and mappings

To relate findings to established functional cerebellar subdivisions, we calculated spatial correspondence of our C-SALE maps with several published cerebellar maps. Comparisons were made either between unthresholded z-maps and continuous parcellations and mappings, or between unthresholded z-maps and discrete parcellations. For both types of comparison, heatmaps were hierarchically clustered. To establish significant spatial correspondence beyond autocorrelation, we accounted for SA and multiple comparisons (*p_variogram, FDR_* < .05) for every comparison.

For continuous maps, spatial correlations were calculated between each *C-SALE map* and *target map* combination. BD and subdomain z-maps were compared to the MDTB and probabilistic cerebellar lobular segmentation. Specifically, for MDTB, we calculated spatial correlation between unthresholded z-maps and the continuous group-level task activation maps reported in the paper of (King et al., 2019). For the probabilistic lobular segmentation (Diedrichsen et al., 2009), C-SALE maps were compared against every lobule separately. In both sets of comparisons, to account for SA, we first calculated the actual spatial correlation between unthresholded C-SALE z-maps and each target map parcel. We then compared this correlation against the null distribution obtained from *target map* correlations with 10,000 SA-preserving surrogate maps, created from C-SALE maps using BrainSmash (Burt et al., 2020; Viladomat et al., 2014). SA is inherent in neuroimaging data, as close regions are generally more likely to have correlated activity (Alexander-Bloch et al., 2018; Burt et al., 2020; Viladomat et al., 2014). It is important to account for these patterns when testing for significant spatial overlap, which may otherwise largely reflect proximity.

For parcellations, because they are discrete mappings, heatmaps report average z-values for each C-SALE map within each (target) parcel, instead of correlations. We assessed correspondence with several parcellations: **1.** The Nettekoven hierarchical cerebellar atlas (mid-granularity symmetrical atlas with sixteen parcels after merging left and right) (Nettekoven et al., 2024); **2.** the Buckner resting-state seven-network cerebellar atlas (Buckner et al., 2011); **3.** the Cole-Anticevic cortical-subcortical atlas (ten cerebellar parcels) (Ji et al., 2019); and lastly **4.** the deterministic cerebellar lobular segmentation (Diedrichsen et al., 2009). Specifically, for each parcellation, we calculated the average z-values of each (sub)domains unthresholded z-maps within each of the parcels. To assess statistical significance, we used a non-parametric test in which observed mean z-values within each parcel of the original C-SALE map were compared against a null distribution of average z-values in SA-preserving pseudorandom surrogate maps created using BrainSmash (Burt et al., 2020). We reused the set of SA-preserving C-SALE surrogate maps used in the continuous map comparisons.

Lastly, we calculated loadings of each voxel within the thresholded C-SALE mask onto the primary (G1) and secondary (G2) functional cerebellar gradients (Guell et al., 2018b) using the LittleBrain Toolbox (Guell et al., 2019) (see *Data availability*). These gradients use orthogonal axes to explain maximal variation in cerebellar resting-state fMRI patterns. The first two axes, those explaining most variance, separate motor from DMN regions (low-to-high loading on G1) and task-unfocused from task-focused regions (low-to-high on G2) (Guell et al., 2018b). Every voxel is colored by the seven-parcel Buckner resting-state network (Buckner et al., 2011) it colocalized with. Note that the sign and unit of gradients are arbitrary.

### 5.10 Meta-analytic connectivity modelling

The functions of cerebellar subareas are tightly connected to those of connected brain regions (King et al., 2019; Magielse et al., 2022; Ramnani, 2006; Wang et al., 2013). Hence, we wanted to examine meta-analytic connectivity profiles across (sub)domains. We adapted the SCALE method, a version of MACM that accounts for baseline activations by sampling null coordinates from the dataset in question (Langner et al., 2014).

Our method tested activations per (sub)domain not against a deterministic null model (as is done in SCALE), but against a whole-brain probabilistic null model (see *section 5.5*). Specifically, for each (sub)domain, we used the thresholded (*p_voxel_* < .001 and k = 50) C-SALE clusters as seed. We first restricted the set of experiments to those that reported at least one peak coordinate within each 3D (cerebellar) seed mask. The set of coordinates included in these experiments were then used as input to whole-brain probabilistic SCALE analyses. These analyses aimed to reveal where in the brain coactivation with the seed regions occurred more than chance, given the biased baseline probability distribution across the brain. Importantly, the set and number of experiments was different from the C-SALE analysis: ultimately, we were able to run analyses for the ‘Action’ BD and four subdomains (N_experiments in seed_ ≥ 17) (Eickhoff et al., 2016).

For our probabilistic implementation of SCALE, the method mirrored that described for C-SALE. The only differences were the set of input experiments (limited to the C-SALE result masks) and the ROI (the whole brain GM mask; >10 % GM probability at 2 mm resolution (Grey10)). Accordingly, for probabilistic SCALE, the null probability map was constructed by normalizing the sum of MA maps of all BrainMap experiments to one within the whole-brain GM mask (Grey10). Again, for each analysis, we created 10,000 null ALE maps by randomly sampling foci from the GM mask, weighted by the baseline reporting probability (**Supplementary Figure 4**). Subsequently, in each analysis, the observed ALE score was compared against the permutation-based null distribution to calculate voxel-wise *p*-values. *P*-values were then thresholded at *p_voxel_* < .001 and k = 50. This revealed where brain-wide coactivations with each (sub)domain’s C-SALE cluster occurred more than chance given baseline activations across the brain.

### 5.11 Correspondence of meta-analytic connectivity maps to brain-wide parcellations

For every MACM result, we calculated spatial overlap with subcortical and cerebral cortical parcellations. This was done for subcortical region (Tian et al., 2020), cerebral cortical-subcortical networks (Ji et al., 2019), and, to interpret findings in terms of brain cytoarchitecture, microstructural cortical types (García-Cabezas et al., 2019; Saberi et al., 2023). For all parcellations separately, we calculated proportions of thresholded MACM maps that colocalized with each parcel in Montreal Neurological Institute (MNI) (2mm) space (Fonov et al., 2009). These proportions are reported as hierarchically clustered heatmaps.

### 5.12 NeuroSynth replication analysis

To validate our method, we contrasted our main findings in BrainMap to results obtained from Neurosynth (v.7), which is an alternative database of the neuroimaging literature (Yarkoni et al., 2011). This database is curated automatically, and the experiments are categorized according to the occurrence of specific terms in their articles texts. We selected 101 Neurosynth terms that were defined in the Cognitive Atlas (cognitiveatlas.org/concepts/i/) and were associated with behavioral and cognitive functions. Out of the 14,371 experiments included in Neurosynth across the 101 selected terms, 5,898 experiments had at least one activation in the dilated cerebellar mask. These experiments were included in (C-S)ALE analysis. See **Supplementary Figure 1b** for a bar plot of the number of experiments included for all 101 terms. The ALE and C-SALE meta-analyses were performed separately for each term associated with seventeen or more experiments (Eickhoff et al., 2016). Of note, since Neurosynth is curated automatically, the sample sizes of included experiments were not available. We thus assumed a fixed sample size of twenty subjects across these experiments (used for convolving foci). By comparing resultant ALE and C-SALE maps, we validated the methodological improvements of C-SALE in NeuroSynth (as described in *section 5.7*) (**Supplementary Figure 2**). The baseline distribution in both datasets was highly similar (r = .96). We last compared spatial distributions of the unthresholded C-SALE z-maps for Neurosynth terms with those for BrainMap behavioral (sub)domains using variogram tests (accounting for SA), followed by FDR-correction (**Supplementary Figures 13, 14**).

## Glossary

*Activation likelihood estimation*

A popular method for performing *coordinate-based meta-analyses.* The method tests where in the brain peak coordinates across a set of imaging experiments converge. To reveal signal amidst (random) noise, the method compares convergence maps against a permuted null model that assumes random spatial distributions of reported activations (or ‘effects’).

*Behavioral topography*

A mapping of behavior(s) (-al domains) to distinct subregions of the brain. In the current study, we use an in-house adaptation of *activation likelihood estimation* to find convergence across a list of behaviors within the cerebellum. The overall mapping of these behaviors to cerebellar subregions constitutes the cerebellar behavioral topography.

*BrainMap database*

One of the two most used neuroimaging databases (along with *NeuroSynth*). In mentioning this database, we refer specifically to its functional (fMRI and PET) data. The BrainMap team has manually curated individual imaging contrasts. These are categorized into distinct, nested task domains and provide information on sample sizes, allowing *activation likelihood estimation* to assign differential confidence to individual activations.

*Cerebellar flatmap*

An unfolded, two-dimensional, representation of the cerebellar gray matter. Since the cerebellum is highly folded, it is normally difficult to visualize its complete outer surface. The flatmap allows a complete overview (in a single image) and has hence become widely adapted. Note that the cerebellar flatmap is typically associated with its own imaging space, the Spatially Unbiased InfraTentorial (SUIT) space.

*Cerebellum-specific activation likelihood estimation*

Our new implementation of *activation likelihood estimation*, that was specifically designed to facilitate unbiased and accurate behavioral mapping in the cerebellum. The method accounts for an unequal distribution of activations (*reported effects or foci*), that is especially pronounced in the cerebellum. In contrast to its name, the method can be flexibly adapted to improve accuracy and specificity of *coordinate-based meta-analyses* in any brain region.

*Coordinate-based meta-analyses*

A collection of methods that summarizes the locations of brain activations across studies. They aim to find regions in the brain that have activity that is significantly elevated above chance relative to other regions. In contrast to more common, effect-size meta-analyses, *coordinate-based meta-analyses* statistically summarize effects within a three-dimensional brain space.

*Meta-analytic coactivation modeling*

An adaptation of *activation likelihood estimation* that aims to reveal coactivation networks of a region-of-interest (or seed). A brain-wide *activation likelihood estimation* analysis is performed, but input experiments are restricted to those mapping to the seed. This reveals brain regions that are coactivated with the seed region significantly above chance. In this study, we account for the unequal baseline of activations (*reported effects or foci*) in the brain.

*NeuroSynth database*

The largest neuroimaging meta-analysis database, that is widely used across the neuroimaging literature. It is often used for *coordinate-based meta-analysis*, and to contextualize neuroimaging findings in terms of involvement in behaviors. *NeuroSynth* used an algorithm to abstract peak coordinates from papers’ main texts.

*Reported effects* or *foci*

Peak coordinates of experimental contrasts, in this case those reported in the *BrainMap* and *NeuroSynth* functional database. These coordinates are often reported in the MNI imaging space, and for *BrainMap* are associated with sample sizes, and belong to one or several task domains.

*Spatial autocorrelation*

*Spatial autocorrelation* refers to the concept that brain areas that are close in space are expected to have more similar characteristics, such as activity. Hence, you might expect to find excessive correlations across proximal structures. Hence, (e.g.,) spin-tests aim to compare correlations of interest to a null-model that quantifies this proximity effect.

*Winner-takes-all parcellations*

A strategy for assigning every cerebellar voxel to a unique (exhaustive and non-overlapping) community, called a parcel. This strategy assigns every voxel fully to the community that it is most similar with.

## Data availability

The BrainMap database is publicly accessible online, via the Sleuth filtered-search application (brainmap.org/software), via the BrainMap Community Portal (portal.brainmap.org), or by comprehensive download when authorized by a collaborative use agreement (CUA) (brainmap.org/collaborations). For this study, we obtained the February 2024 data release through CUA. The NeuroSynth database is publicly accessible online neurosynth.org<u>/</u>, and full data was obtained from github.com/neurosynth/neurosynth-data. Full intermediate data resulting from this publication, including figure notebooks, are uploaded to (github.com/CNG-LAB/cerebellum_specific_ALE). Final figures were put together in Inkscape and can be obtained for reuse upon request.

## Code availability

All programmatic code used to obtain the results in this article are made available on GitHub: (github.com/CNG-LAB/cerebellum_specific_ALE). All code necessary to perform bias-accounting coordinated-based meta-analyses, for the whole brain or any volumetric brain region-of-interest, are made available. Also made available is our graphical processing unit implementation of NiMARE which helps speed up modeled activation map calculation (github.com/CNG-LAB/nimare-gpu), and works for both classic activation likelihood estimation, and deterministic and probabilistic versions of cerebellum-specific activation likelihood estimation (as in the current study). The LittleBrain toolbox, used to relate our maps to cerebellar gradients, has been openly shared by Guell et al. at github.com/xaviergp/littlebrain (Guell et al., 2019).

## Materials and Correspondence

Correspondence can be addressed to Neville Magielse and Sofie L. Valk. Permanent address: Max Planck Institute for Human Cognitive and Brain Sciences, Dr. Sofie L. Valk, Stephanstraße 1A, 04103 Leipzig, Saxony, Germany. Telephone: <u>+49 341 9940-2658</u> | Fax: <u>+49 341 9940-104</u>.

## Ethics statement

This study uses data from the publicly available BrainMap meta-analytic database. Data consists of aggregated fMRI and PET data. Specifically, only peak coordinate locations and sample size were used (and available), and hence no individual participants could be identified. No contact was made (nor possible) with any participants, nor was any individual’s data handled at any point.

## Author contributions

**NM.** Conceptualization, Methodology, Software, Validation, Formal Analysis, Investigation, Data Curation, Writing – Original Draft, Writing – Review & Editing, Visualization, Project Administration.

**AM.** Conceptualization, Methodology, Validation, Writing – Review & Editing.

**SBE.** Conceptualization; Methodology, Software; Resources, Supervision, Funding Acquisition.

**PTF.** Methodology, Resources.

**AS*.** Conceptualization, Methodology, Software, Validation, Formal analysis, Resources, Writing – Review & Editing, Visualization, Project Administration.

**SLV*.** Conceptualization, Validation, Investigation, Writing – Review & Editing, Supervision, Funding Acquisition.

***^#^*** *Authors **AS** and **SLV** contributed to this manuscript equally and hence share last authorship*.

## Acknowledgements

We would like to thank Lennart Frahm for insightful discussions on the activation likelihood estimation method and statistics.

