## Supplementary Information for "A Bias-Accounting Meta-Analytic Approach Refines and Expands the Cerebellar Behavioral Topography"

Magielse et al.

### Supplementary Figures

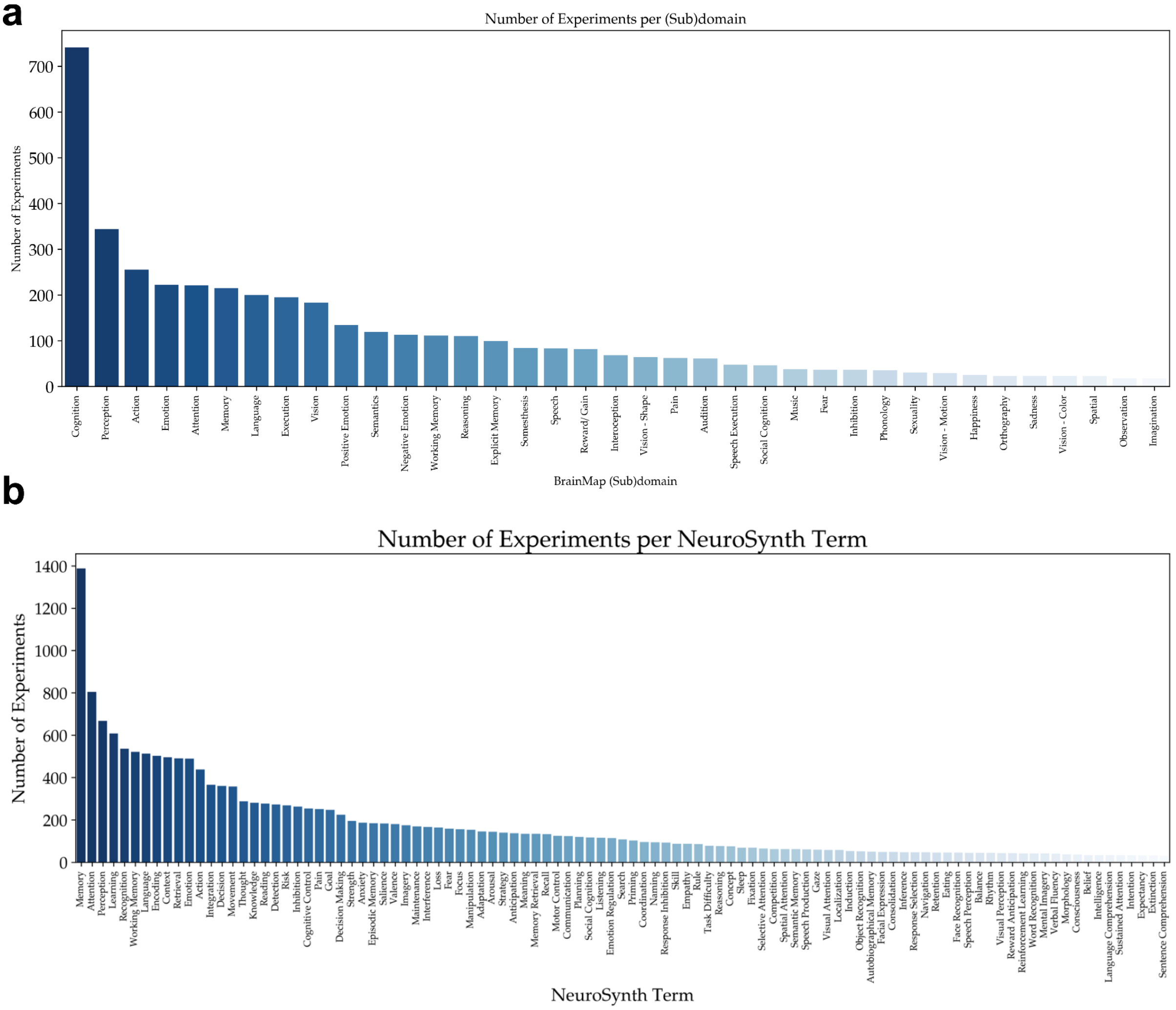


**Supplementary Figure 1: Number of experiments included in each analysis.** (**a-b**) The number of experiments that were included for each C-SALE (and thus ALE) analysis for the main (**a**) and replication (**b**) sample. These experiments thus included at least one cerebellar peak coordinate within the dilated cerebellar mask. In both subfigures, datasets are ordered from large to small. Full experimental statistics can be found on GitHub.

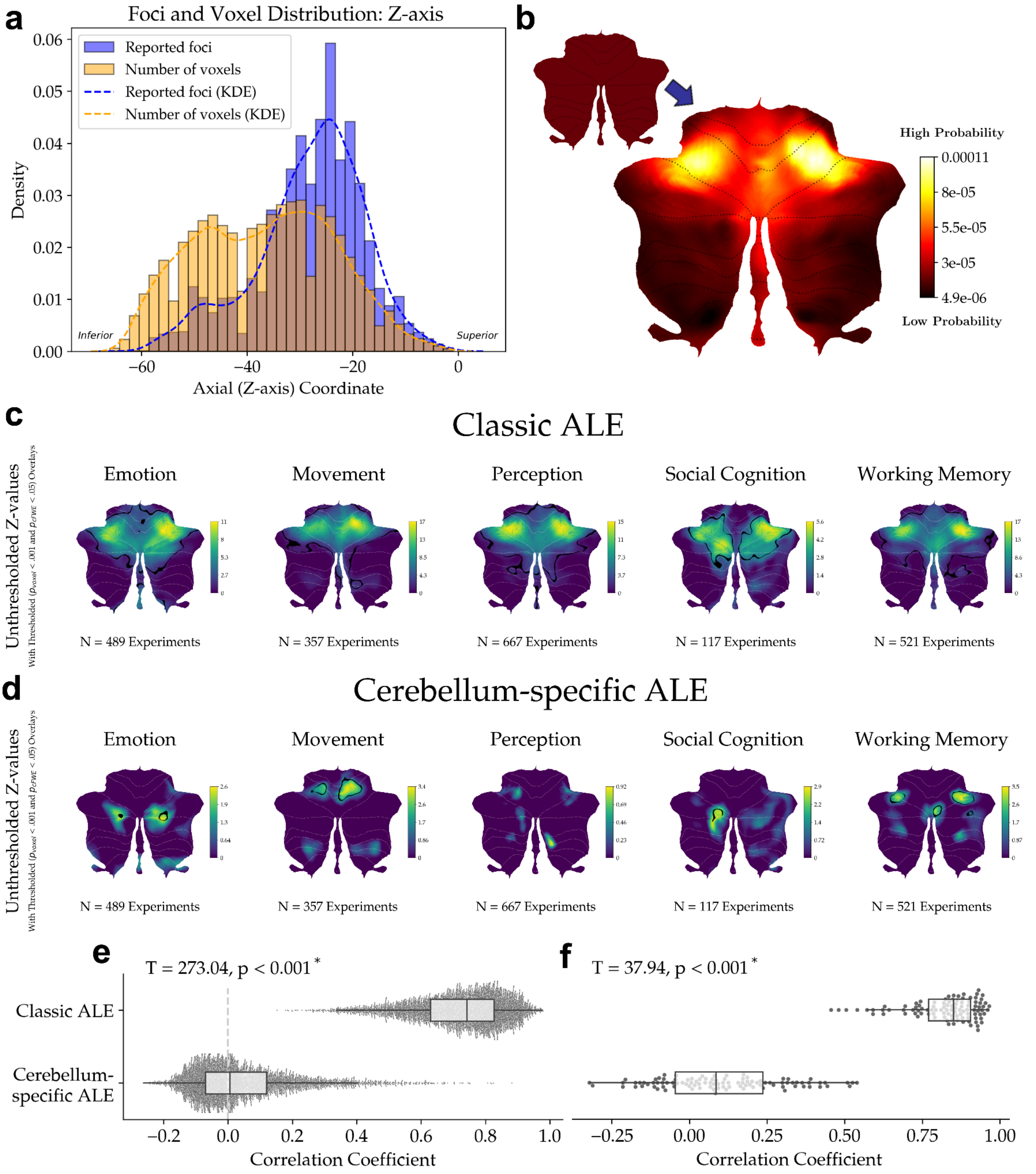


**Supplementary Figure 2: Replication of improvements of the cerebellum-specific ALE method over classic ALE for cerebellar meta-analysis.** Methodological replication of cerebellum-specific activation-likelihood estimation (C-SALE) improvements in the NeuroSynth database. (**a**) Maps the distribution of reported effects versus the number of cerebellar voxels across the Z-axis. (**b**) Illustrates the null hypotheses of classic ALE (smaller flatmap) and C-SALE (larger flatmap). (**c, d**) Show differences between classic ALE (**c**) and C-SALE (**d**) maps for five terms. Unthresholded z-maps are shown with an outline of clusters that reached convergence ($p_{voxel} < .001$ and k = 50 for C-SALE and $p_{\mathrm{cFWE.}} < .05$ using a height threshold of $p_{voxel} < .001$ for classic ALE). Terms are ordered alphabetically. (**e**) Shows distributions of correlation coefficients between each pair of unthresholded z-maps. Correlations were significantly lower in C-SALE (T = 273.04; $p < .001$*). The vertical dotted line illustrates the null assumption that within-cerebellum spatial correlation centers on 0.0 in accurate within-cerebellar CBMA. (**f**) Shows correlations of each z-map with the biased baselines distribution (pictured in **b**). C-SALE resembled the baseline significantly less (T = 37.94; $p < .001$*). KDE = kernel density estimation.

#### Supplementary Information

Magielse et al.

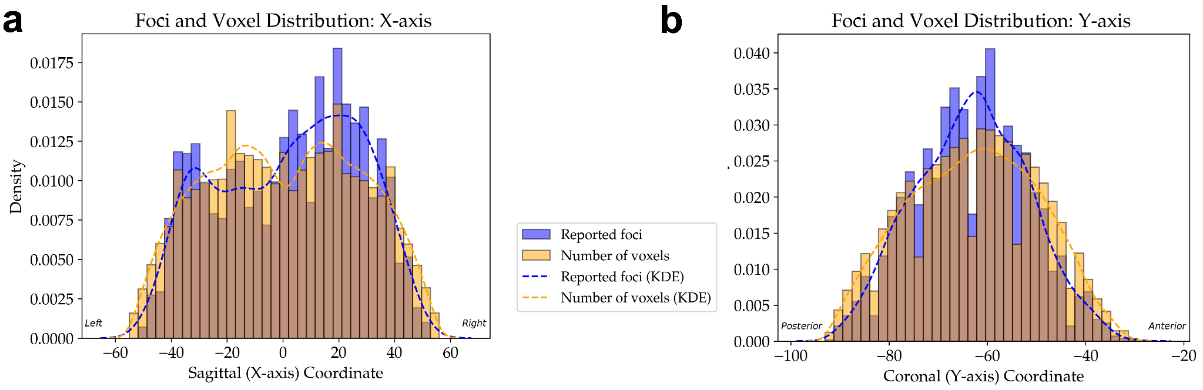


**Supplementary Figure 3: Voxel and foci distributions across other brain axes. (a-b**) Distributions of foci (orange bars) relative to voxel (blue bars) density across the X- (sagittal) and Y-axes (coronal) are shown. Whereas a substantial shift in foci distribution relative to that of voxels is noted across the Z-axis (**Figure 1**), such shifts are more subtle for the X- and Y-axes (**a, b**). Across both axes, foci tend to follow the voxel distribution. Nonetheless, foci map sparsely to the extremities of both axes: there are relatively fewer far-left, -right, -posterior, and -anterior foci (**a, b**). In (**a**) specifically, foci appear to be somewhat overrepresented in the right relative to the left cerebellum.  Clear discrepancies are not evident in (**b**), where foci follow the distribution of voxels, albeit with a narrower distribution.

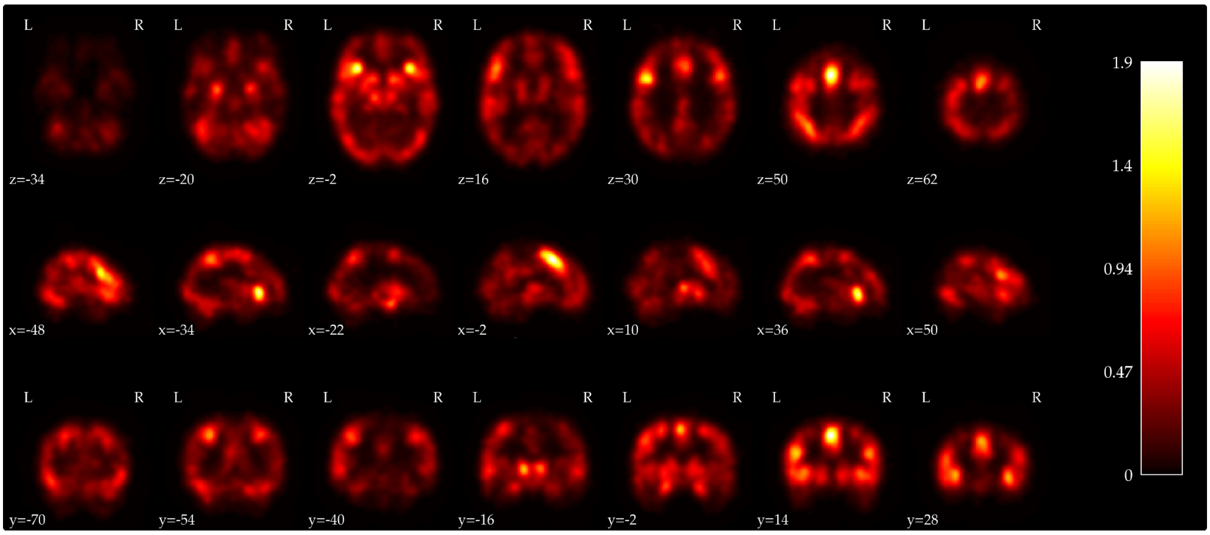


**Supplementary Figure 4: Reported effects are unequally distributed across the brain.** To construct cerebellar null models that account for the unequal distribution of reported effects, we first created a whole-brain probability distribution from the BrainMap data. This allowed us to 1. account for spatial uncertainties in cerebellar foci; and 2. flexibly adapt our method to any brain region-of-interest. Specifically, all normal mapping (healthy participants) experiments across domains were combined by first convolving each experimental focus with a kernel with full width at half maximum (FWHM) inversely proportional to sample size of the respective experiment. The smoothed map for each experiment was then summed, creating the whole brain probability distribution shown here. This then allowed us to isolate the cerebellum and normalize the sum of probabilities to one, to represent a null space of finding a focus at any cerebellar voxel. Note the several activation hotspots within the whole-brain probability distribution. The cerebellum is generally underreported and shows a sharp superior-inferior boundary. More generally, the probability of finding a focus differs greatly across the brain, with connected implications for the standard null hypothesis, and hence the findings of previous meta-analyses. The colorbar represents summed probabilities across all experiments and can thus exceed one.

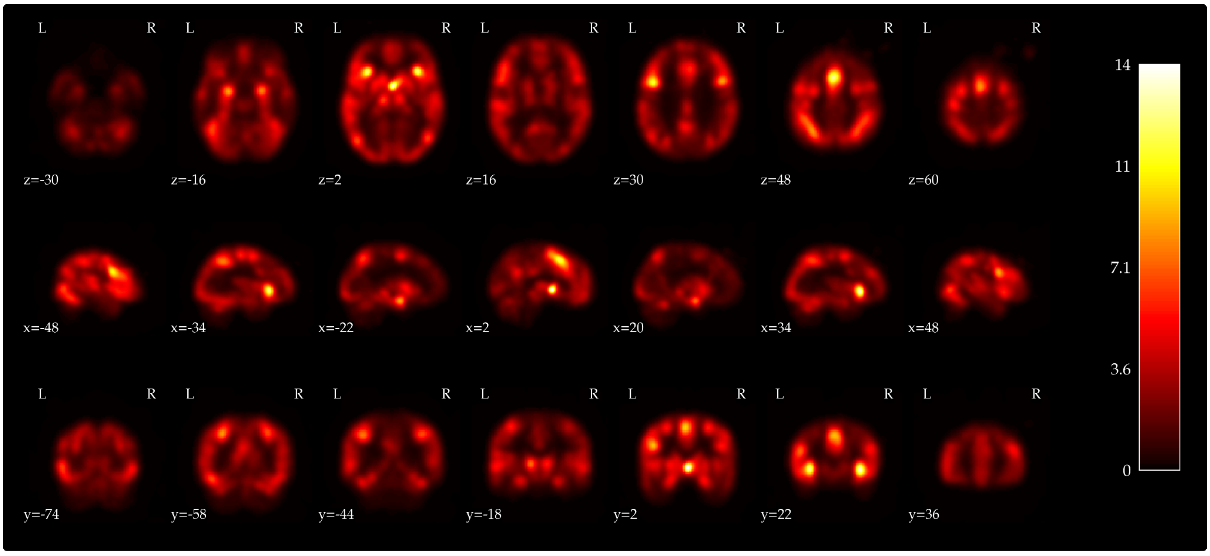


**Supplementary Figure 5: Replication of unequal distributions of reported effects across the brain.** We replicated the unequal whole-brain foci probability distribution in NeuroSynth. Again, all normal mapping experiments across the 101 terms were combined by first convolving each experimental focus with a kernel with full width at half maximum (FWHM) proportional to an assumed sample size of twenty participants per experiment. We assumed an average number of experiments because NeuroSynth does not provide experimental sample size information. The smoothed map for each experiment was then summed, creating the whole brain probability distribution shown here. This then allowed us to isolate the cerebellum and normalize the sum of probabilities to one, representing a null space of finding a focus at any cerebellar voxel. This map was then used to sample the null model for NeuroSynth C-SALE analyses. Note that the NeuroSynth whole-brain probability distribution closely matches that created from BrainMap data. Activation hotspots can be found in similar locations. The cerebellum is again underreported and shows a sharp superior-inferior boundary. Also in NeuroSynth, the probability of finding a focus differs greatly across the brain, with connected implications for the standard null hypothesis, and hence the findings of previous meta-analyses. The colorbar represents summed probabilities across all experiments and can thus exceed one.

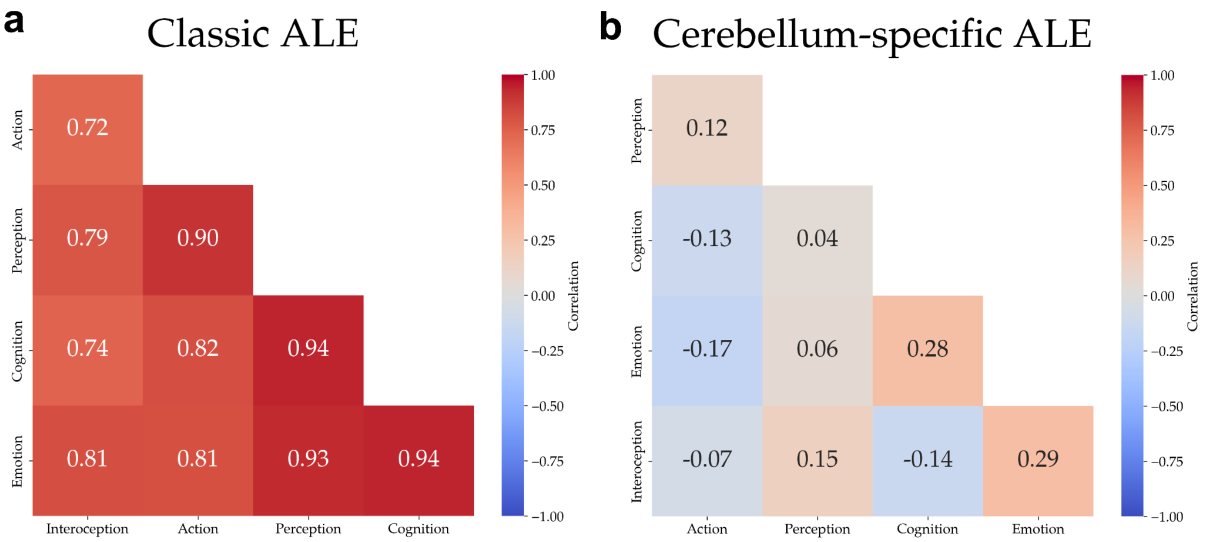


**Supplementary Figure 6: Comparison of behavioral domain spatial correlations between classic ALE and cerebellum-specific ALE (C-SALE).** To assess specificity improvements of C-SALE over classic ALE, we assessed spatial correlations across behavioral domains (BDs) (**a,b**). Given that we aimed to map behaviors to cerebellar subregions, low spatial correlation between (sub)domains can be assumed to signal specificity. We report hierarchically clustered spatial correlations unthresholded z-maps for each *BD x BD* combination. This shows that while regular ALE reports universally strong positive correlations (**a**), C-SALE reports moderate positive and negative correlations (**b**). This indicates that C-SALE, in contrast to regular ALE, can differentiate cerebellar subregions converging in different BDs.

**a**

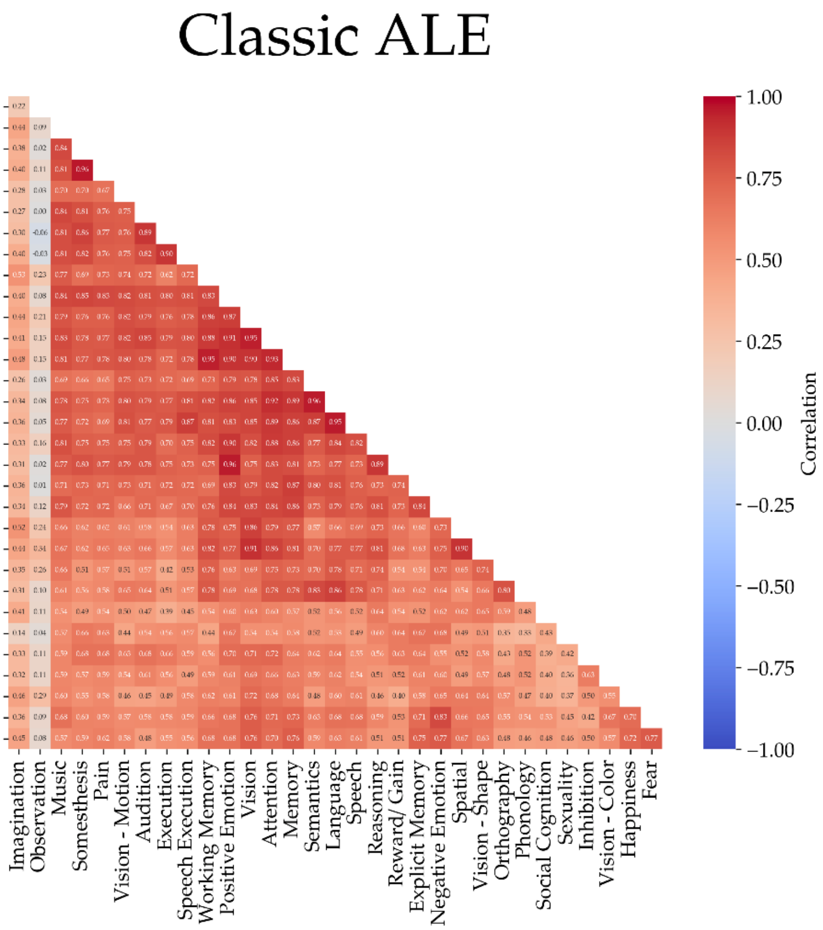
Classic ALE
Observation
Music
Somesthesis
Pain
Vision - Motion
Audition
Execution
Speech Execution
Working Memory
Positive Emotion
Vision
Attention
Memory
Semantics
Language
Speech
Reasoning
Reward/ Gain
Explicit Memory
Negative Emotion
Spatial
Vision - Shape
Orthography
Phonology
Social Cognition
Sexuality
Inhibition
Vision - Color
Happiness
Fear
Sadness
Imagination
Observation
Music
Somesthesis
Pain
Vision - Motion
Audition
Execution
Speech Execution
Working Memory
Positive Emotion
Vision
Attention
Memory
Semantics
Language
Speech
Reasoning
Reward/ Gain
Explicit Memory
Negative Emotion
Spatial
Vision - Shape
Orthography
Phonology
Social Cognition
Sexuality
Inhibition
Vision - Color
Happiness
Fear
1.00
0.75
0.50
0.25
0.00
−0.25
−0.50
−0.75
−1.00
Correlation

**b**

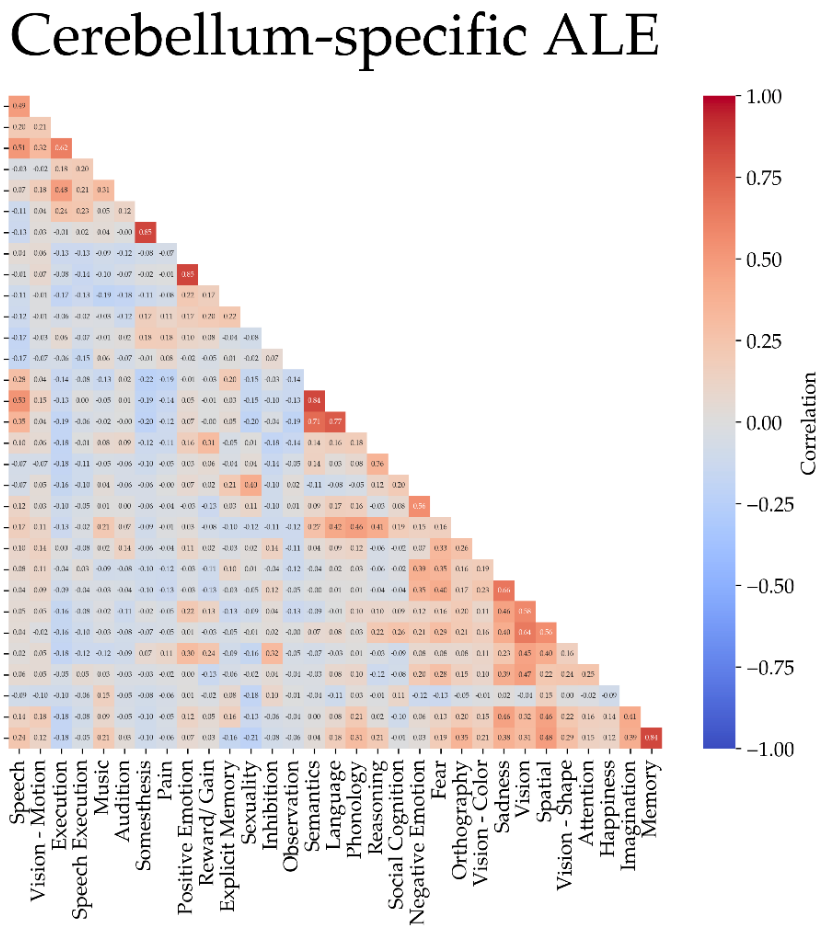
Cerebellum-specific ALE
Vision - Motion
Execution
Speech Execution
Music
Audition
Somesthesis
Pain
Positive Emotion
Reward/ Gain
Explicit Memory
Sexuality
Inhibition
Observation
Semantics
Language
Phonology
Reasoning
Social Cognition
Negative Emotion
Fear
Orthography
Vision - Color
Sadness
Vision
Spatial
Vision - Shape
Attention
Happiness
Imagination
Memory
Working Memory
Speech
Vision - Motion
Execution
Speech Execution
Music
Audition
Somesthesis
Pain
Positive Emotion
Reward/ Gain
Explicit Memory
Sexuality
Inhibition
Observation
Semantics
Language
Phonology
Reasoning
Social Cognition
Negative Emotion
Fear
Orthography
Vision - Color
Sadness
Vision
Spatial
Vision - Shape
Attention
Happiness
Imagination
Memory
1.00
0.75
0.50
0.25
0.00
−0.25
−0.50
−0.75
−1.00
Correlation

**Supplementary Figure 7: Comparison of behavioral subdomain spatial correlations between classic ALE and cerebellum-specific ALE.** We also assessed spatial correlations for each *subdomain x subdomain* combination across both methods. Again, low spatial correlations are taken to indicate that the method can disentangle cerebellar locations of convergence. We report hierarchically clustered spatial correlations of unthresholded z-maps for each *subdomain x subdomain* combination. This shows that while ALE reports only strong positive correlations (**a**) (except for Action Observation), cerebellum-specific ALE (C-SALE) generally reports moderate positive and negative correlations (**b**). This again supports the notion that C-SALE assigns behavioral subdomains to specific cerebellar subregions. Several high correlations stand out. These correspond primarily to closely related behavioral constructs that likely have substantial overlap in their set of experiments.

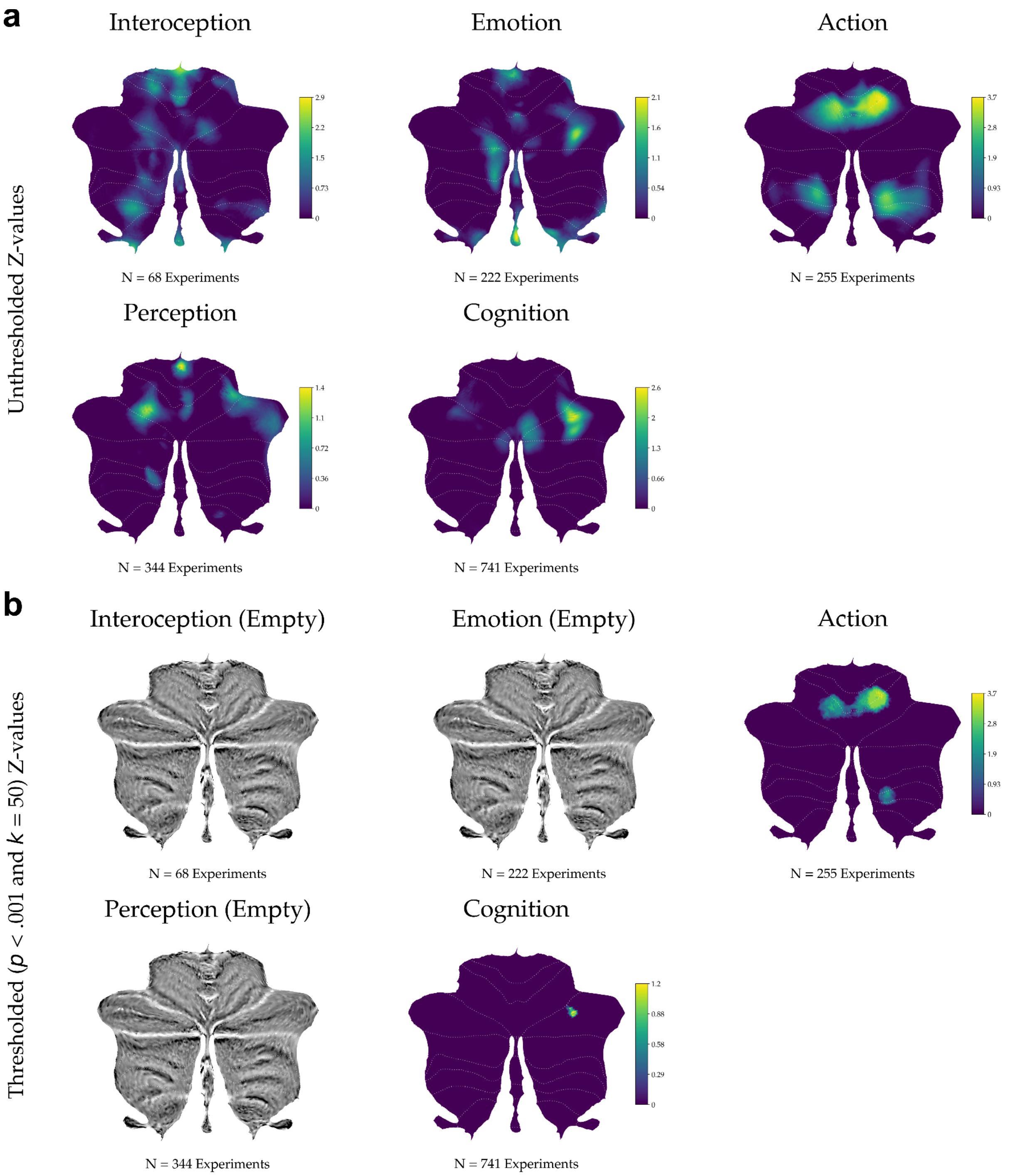


**Supplementary Figure 8. Unthresholded and thresholded behavioral domain cerebellum-specific ALE maps.** Full z-maps are shown for unthresholded (**a**) and thresholded ($p$ < .001 and $k$ = 50) (**c**) cerebellum-specific ALE results. In (**b**), behavioral domains (BDs) that did not reach convergence are represented by empty cerebellar flatmaps. All maps are ordered by BD sample size.

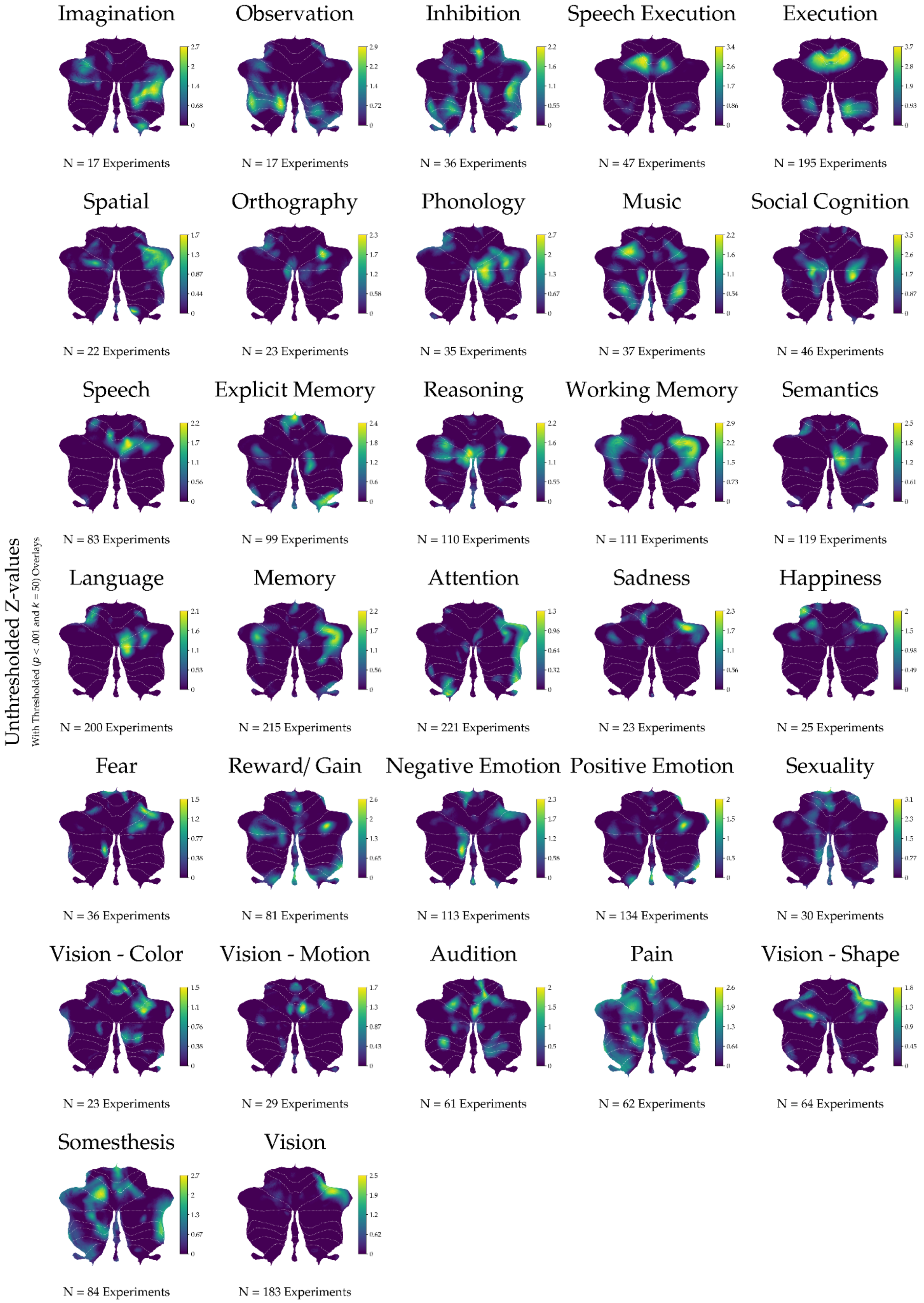


**Supplementary Figure 9: Untresholded subdomain cerebellum-specific ALE z-maps.** All subdomain unthresholded cerebellum-specific ALE z-maps are shown. Subdomains are first ordered alphabetically by behavioral domain (BD), then by sample size from small to large.

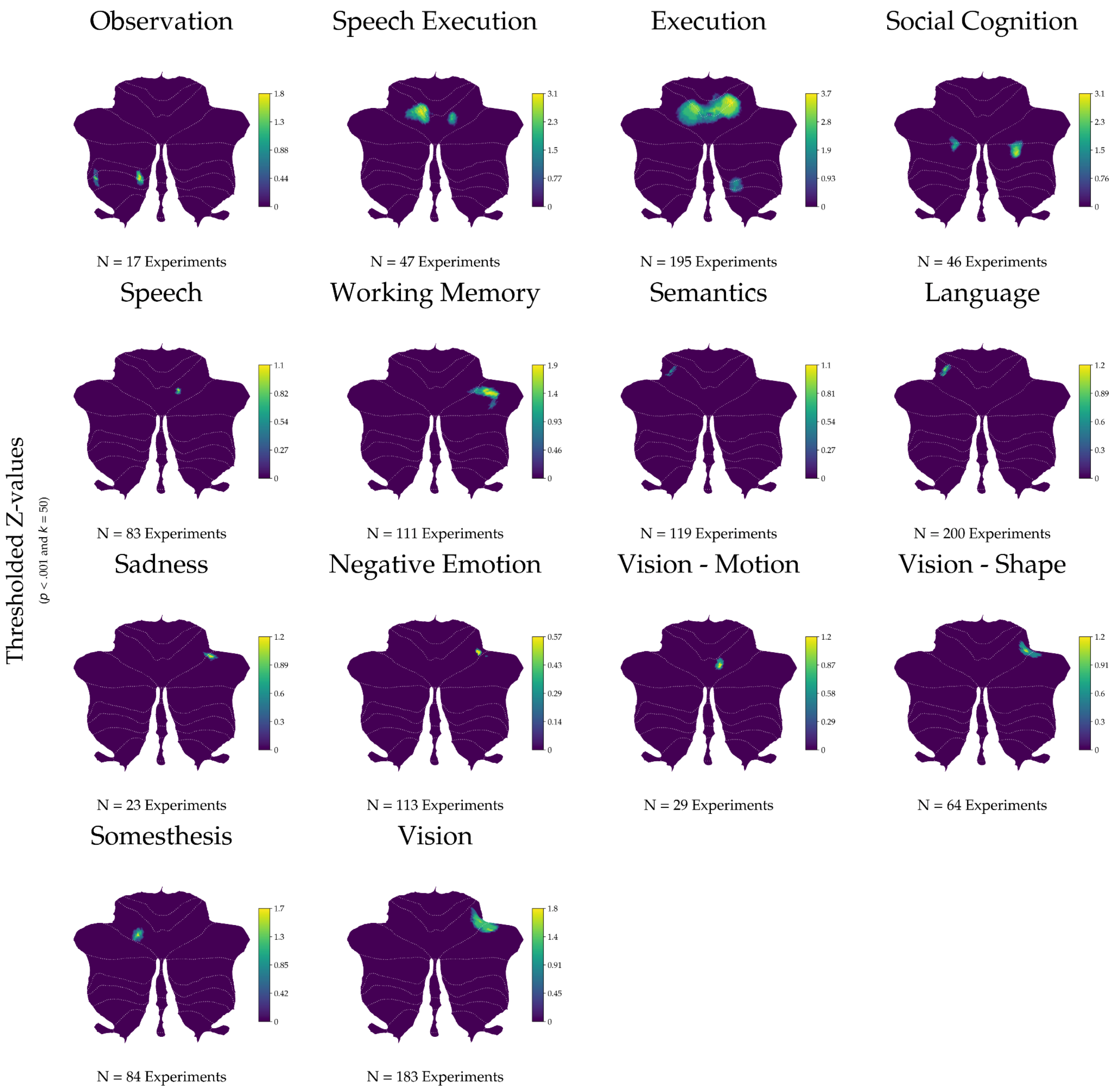


**Supplementary Figure 10: Thresholded subdomain cerebellum specific ALE z-maps.** All thresholded ($p$ < .001 and $k$ = 50) cerebellum-specific ALE z-maps are shown. Subdomains are first ordered by behavioral domain (BD), then by sample size from small to large.

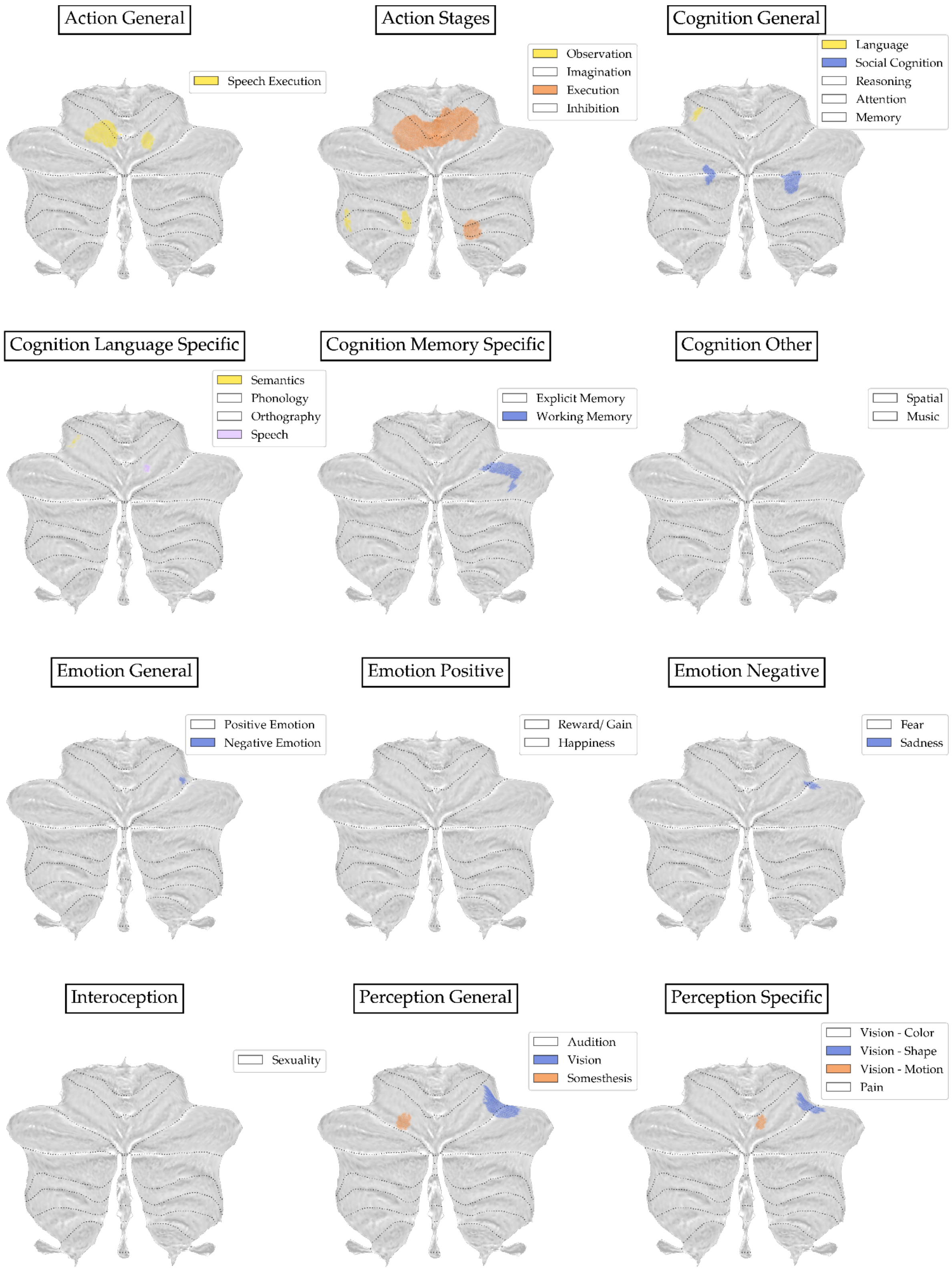


**Supplementary Figure 11: Binary localisations for thresholded cerebellum-specific ALE maps across behavioral subdomains.** Binary locations of converging regions across subdomains ($p$ < .001 and $k$ = 50) are plotted onto cerebellar flatmaps. Each of the BrainMap subdomains is ordered into an author-imposed semantically meaningful category. Colors are repeated to maximize (color blind) visibility and have no further meaning. White labels indicate convergence did not reach the threshold in the respective subdomain. Clusters in each flatmap are ordered from large (top label) to small (bottom label) behavioral subdomain sample size.

#### Supplementary Information

Magielse et al.

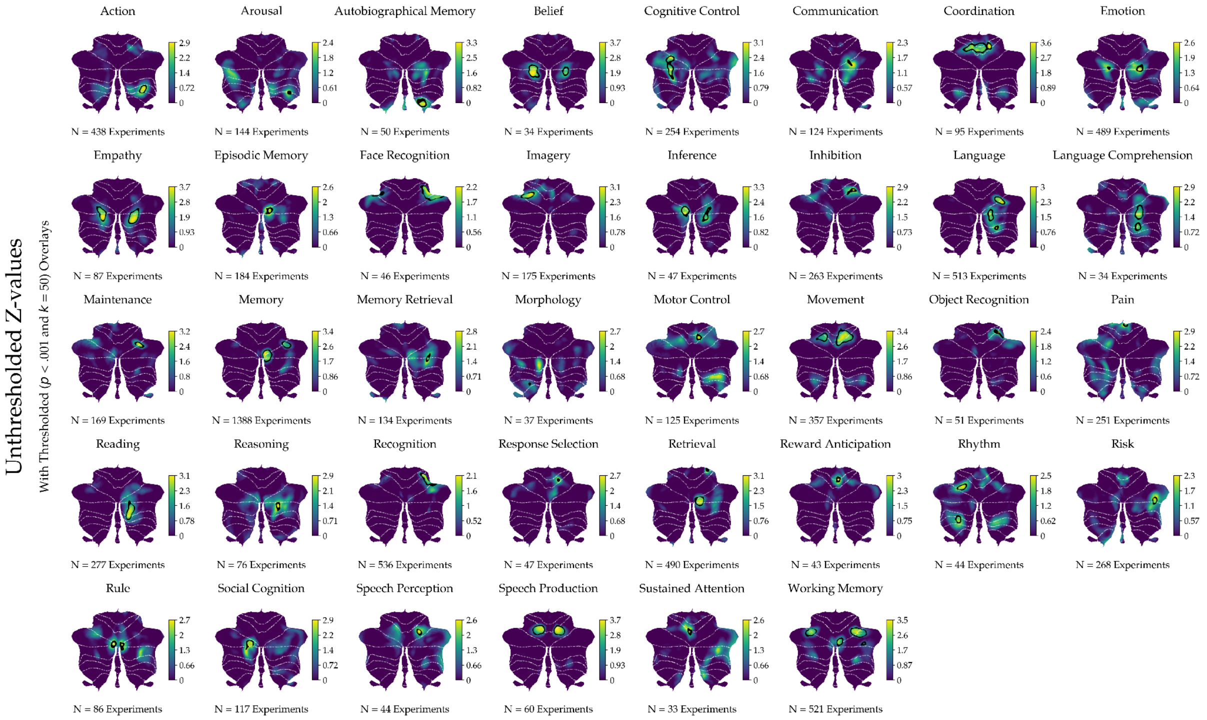


**Supplementary Figure 12: Cerebellum-specific ALE maps for behavioral terms in NeuroSynth.** C-SALE results in the NeuroSynth replication sample, as in **Figure 3.** Thirty-eight of 101 behavioral terms reached convergence. Unthresholded z-maps are shown with an outline of the locations that reached convergence ($p_{voxel}$ < .001 and k = 50). Terms are ordered alphabetically.

#### Supplementary Information

Magielse et al.

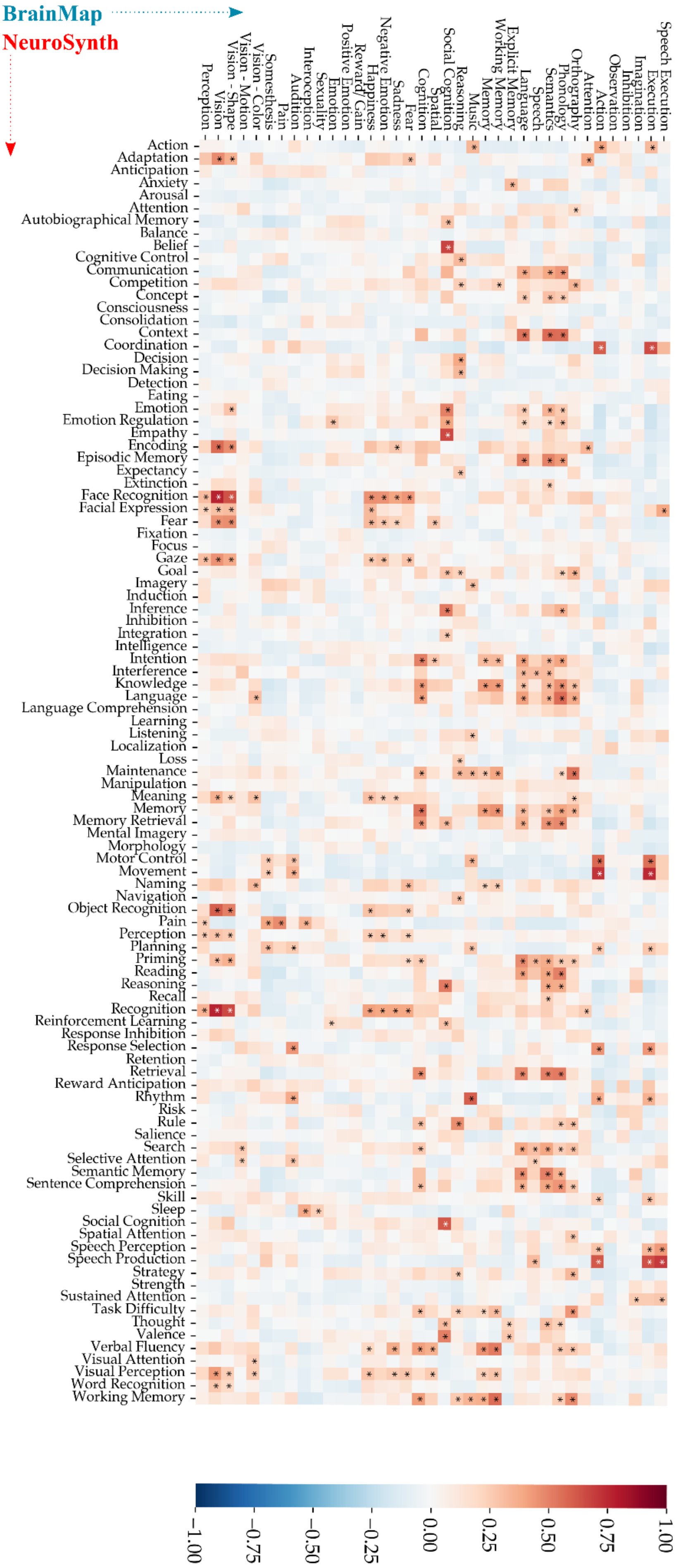

**Supplementary Figure 13: Spatial correlations between BrainMap and NeuroSynth C-SALE maps.** To assess correspondence of C-SALE maps between the main (BrainMap) and replication (NeuroSynth) samples, we ran spatial correlations between each *(sub)domain x term* pair. Variograms were constructed for BrainMap C-SALE maps to account for spatial autocorrelation (SA). Asterisks indicate combinations that remained significant after accounting for SA and multiple comparisons ($p_{variogram,\ FDR} < .05$). Asterisk color (white/ black) was set to aid visibility and is not relevant to the interpretation of results.

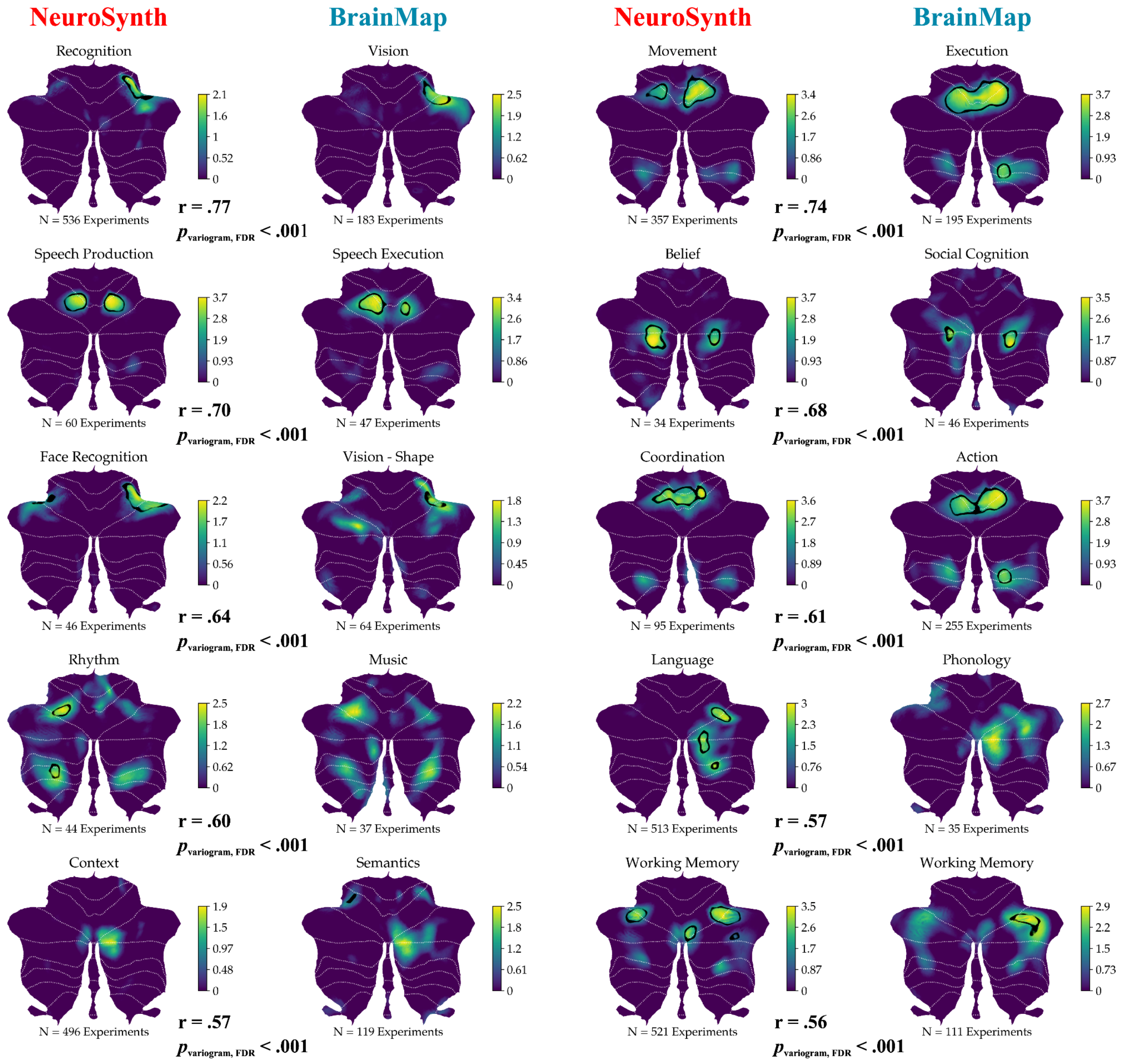


**Supplementary Figure 14: Top spatial correlations between BrainMap and NeuroSynth C-SALE maps.** To highlight correspondence between BrainMap behavioral (sub)domains and NeuroSynth behavioral terms, we visualized ten example pairs that were strongly spatially correlated. These ten pairs were selected by finding pairs with the highest overall R-values, with the restriction that each map could only appear once. r Denotes spatial correlation between shown pairs, whereas $p_{variogram, FDR}$ denotes FDR-corrected variogram-based p-values.

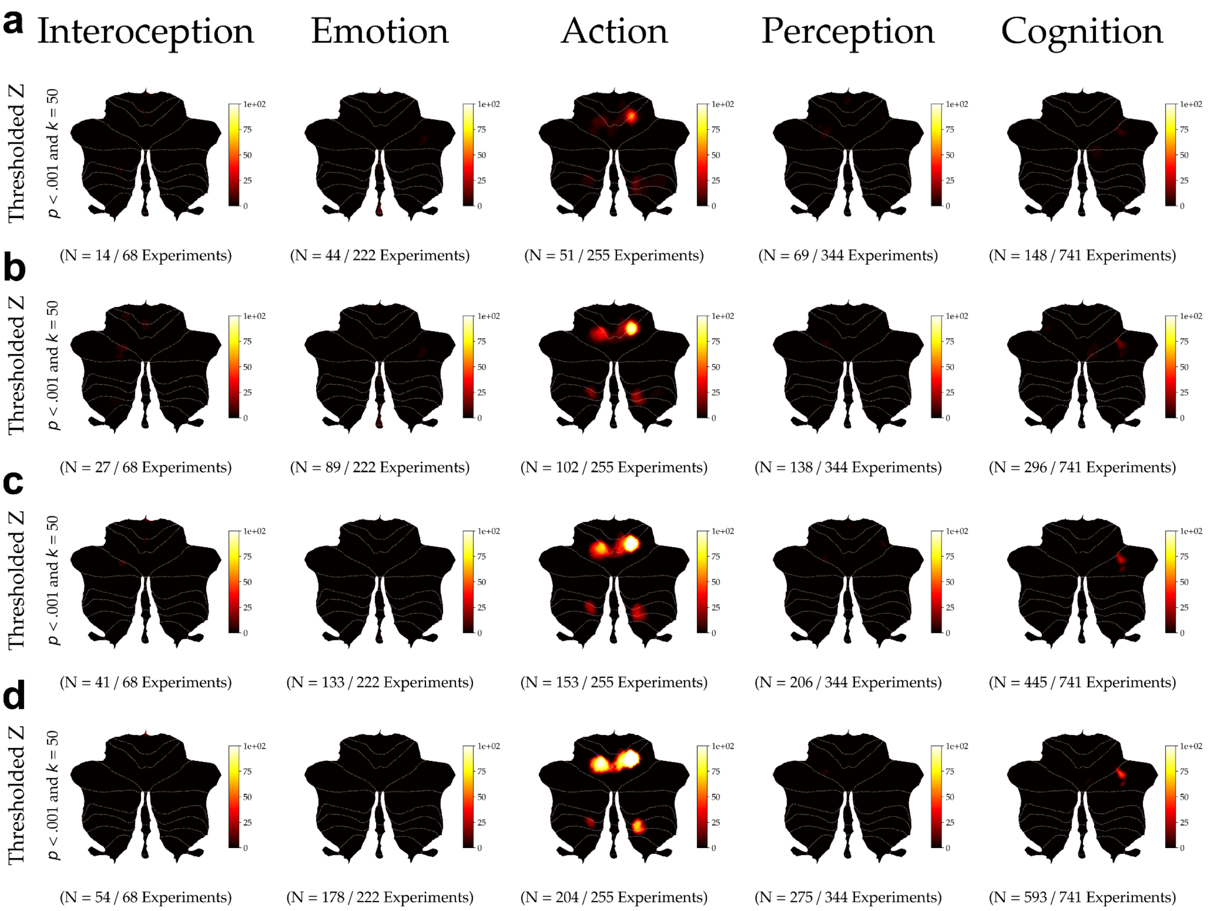


**Supplementary Figure 15: Significance heatmaps for behavioral domains across sliding subsampling proportions.** To assess how consistently convergence passed the threshold ($p$ < .001 and k = 50) across behavioral domains (BDs), we mapped the percentage of 50 subsamples that reached convergence at each voxel. We did this for subsamples at 0.2 (**a**), 0.4 (**b**), 0,6 (**c**), and 0.8 (**d**) proportions of the total BD sample size. In each subfigure, BDs are ordered by sample size from small to large. All color bars map from 0 to 100 percent.

#### Supplementary Information

Magielse et al.

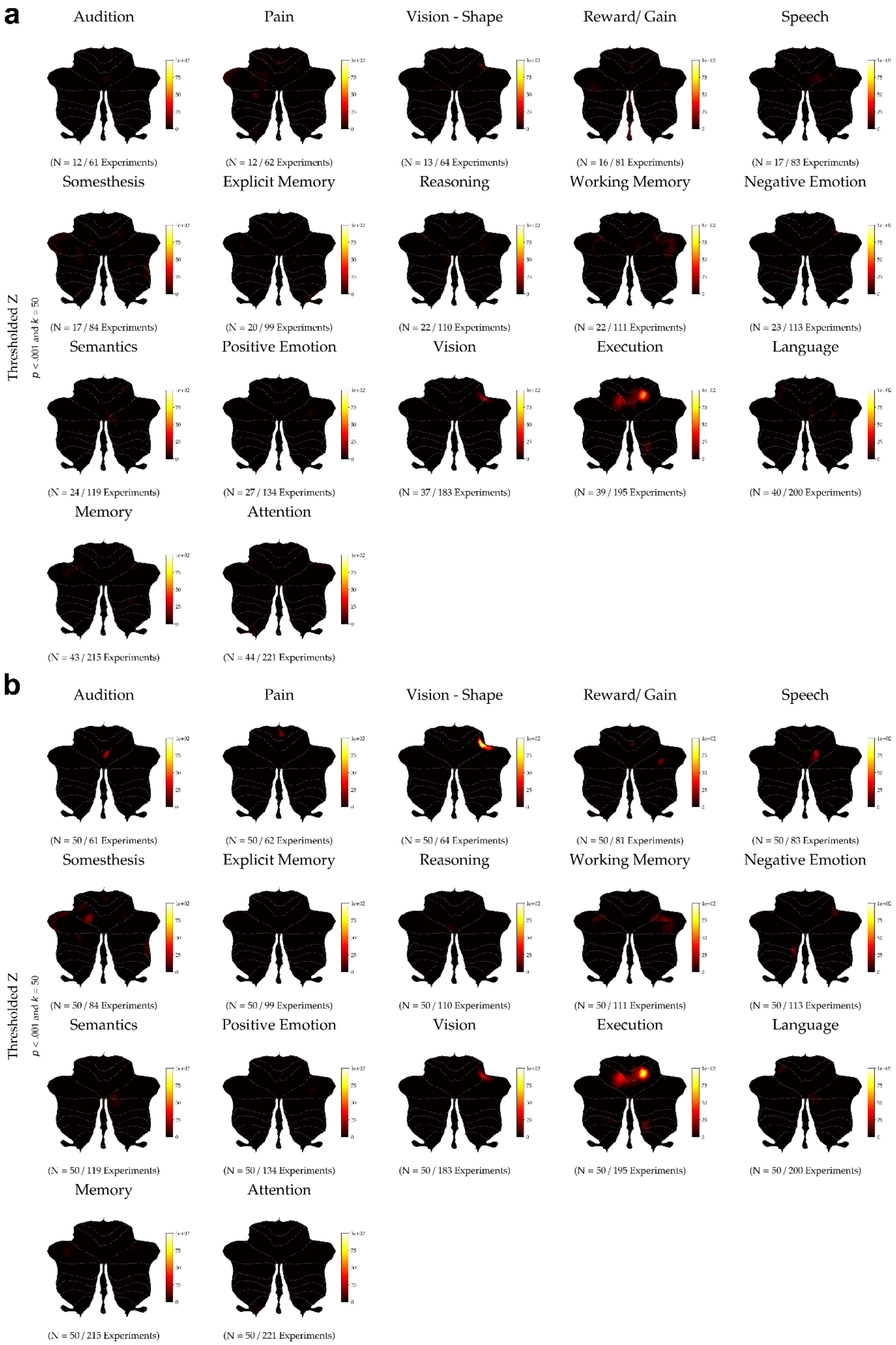

**Supplementary Figure 16: Significance heatmaps for behavioral subdomains for fixed sample sizes and sampling proportions.** To assess how consistently convergence passed the threshold ($p$ < .001 and k = 50) across subdomains we mapped the percentage of 50 subsamples that reached convergence at each voxel. We did this for random subsamples of N = 50 experiments (**a**) and for a 0.2-proportion of the total subdomain sample size (**b**). In each subfigure, subdomains are ordered by sample size from small to large. All color bars map from 0 to 100 percent.

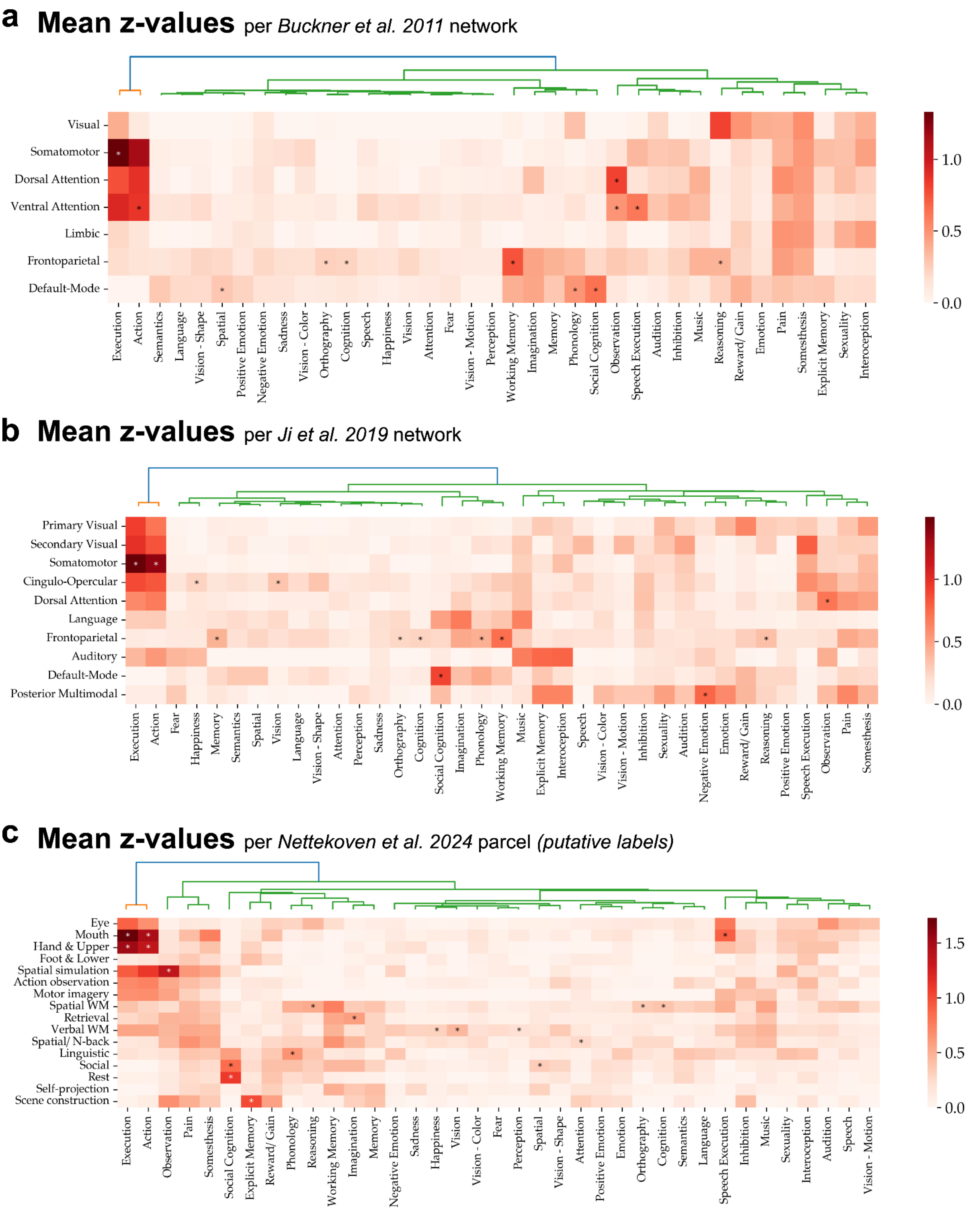


**Supplementary Figure 17: Correspondence between cerebellum-specific ALE maps and previous cerebellar parcellations.** We assessed spatial correspondence of our (sub)domain cerebellum-specific ALE (C-SALE) maps with existing cerebellar functional subdivisions. Specifically, we calculated the mean z-value for each C-SALE map within each of the resting-state networks of (Buckner et al., 2011) (**a**), the cerebellar cortical-subcortical networks of (Ji et al., 2019) (**b**), and each parcel of the mid-level symmetric hierarchical atlas of (Nettekoven et al., 2024) (**c**). Note that (**c**) shows the same heatmap as **Figure 5d**, but with putative behavioral labels to provide a notion of behavioral correspondence between both sets of maps. Importantly, these labels do not represent a definitive mapping of functions to these parcels and may change as we learn more about what cerebellar functions they separate. All heatmaps were hierarchically clustered. Variograms were constructed for

#### Supplementary Information

Magielse et al.

C-SALE maps to account for spatial autocorrelation (SA). Asterisks indicate combinations that remained significant after accounting for SA and multiple comparisons ($p_{variogram, FDR} < .05$). Asterisk color (white/ black) was set to aid visibility and is not relevant to the interpretation of results.

#### Supplementary Information

Magielse et al.

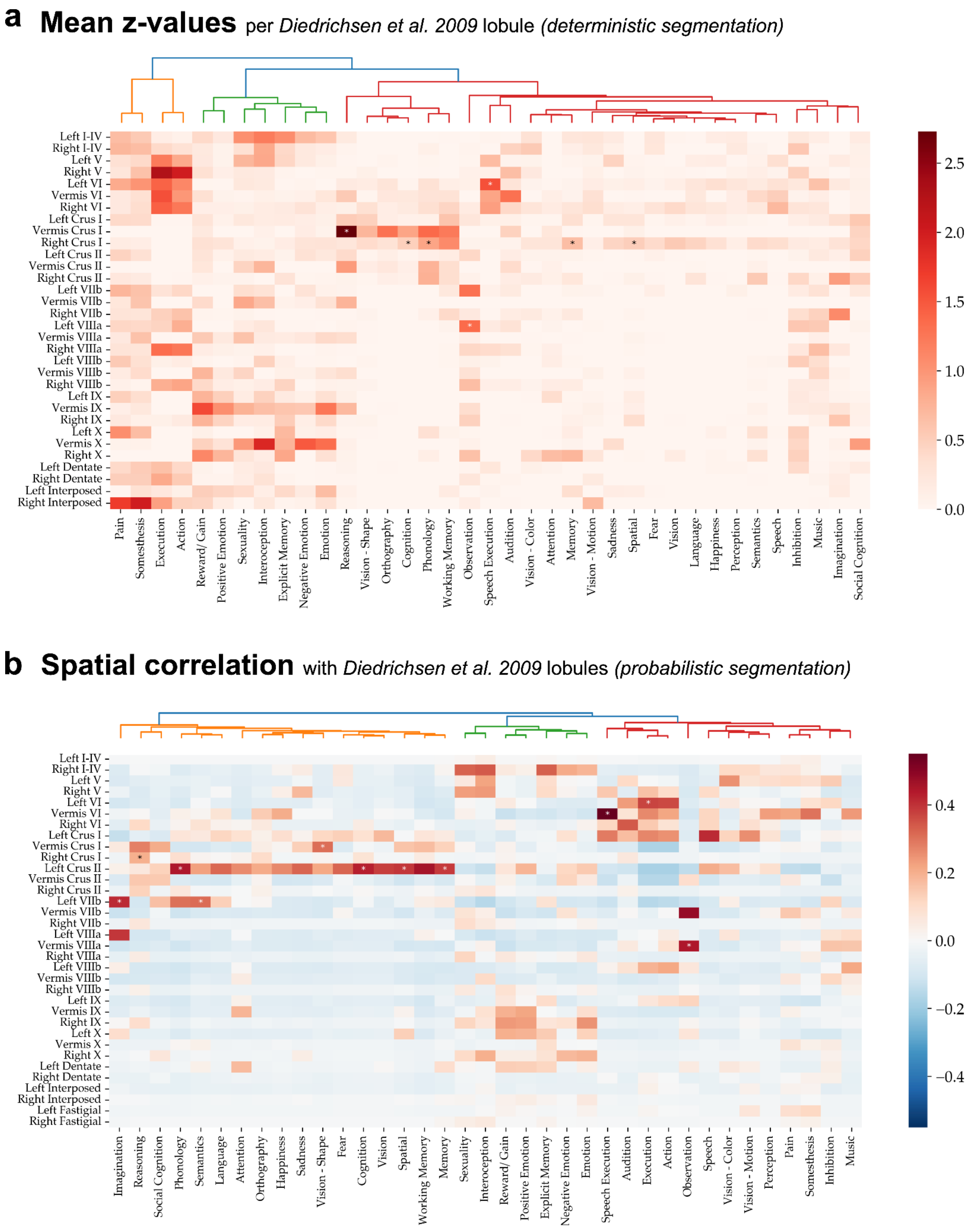


**Supplementary Figure 18: Correspondence between cerebellum-specific ALE maps and cerebellar lobules.** To assess how well lobular boundaries correspond to locations of behavioral convergence, we assessed spatial correspondence with lobular segmentation. (**a**) Shows mean z-values for each cerebellum-specific ALE (C-SALE) map within each cerebellar lobule (Diedrichsen et al. 2009; deterministic segmentation (Diedrichsen et al., 2009)). (**b**) Shows spatial correlations between each unthresholded C-SALE z-map and the probabilistic lobular segmentation of (Diedrichsen et al., 2009). In both, left and right lobules were not merged, to highlight potential asymmetry in C-SALE mappings. All heatmaps were hierarchically clustered. Variograms were constructed for C-SALE maps to account for spatial autocorrelation (SA). Asterisks indicate combinations that remained significant after accounting for SA and multiple comparisons ($p_{variogram,\ FDR} < .05$). Asterisk color (white/ black) was set to aid visibility and is not relevant to interpretation of results.

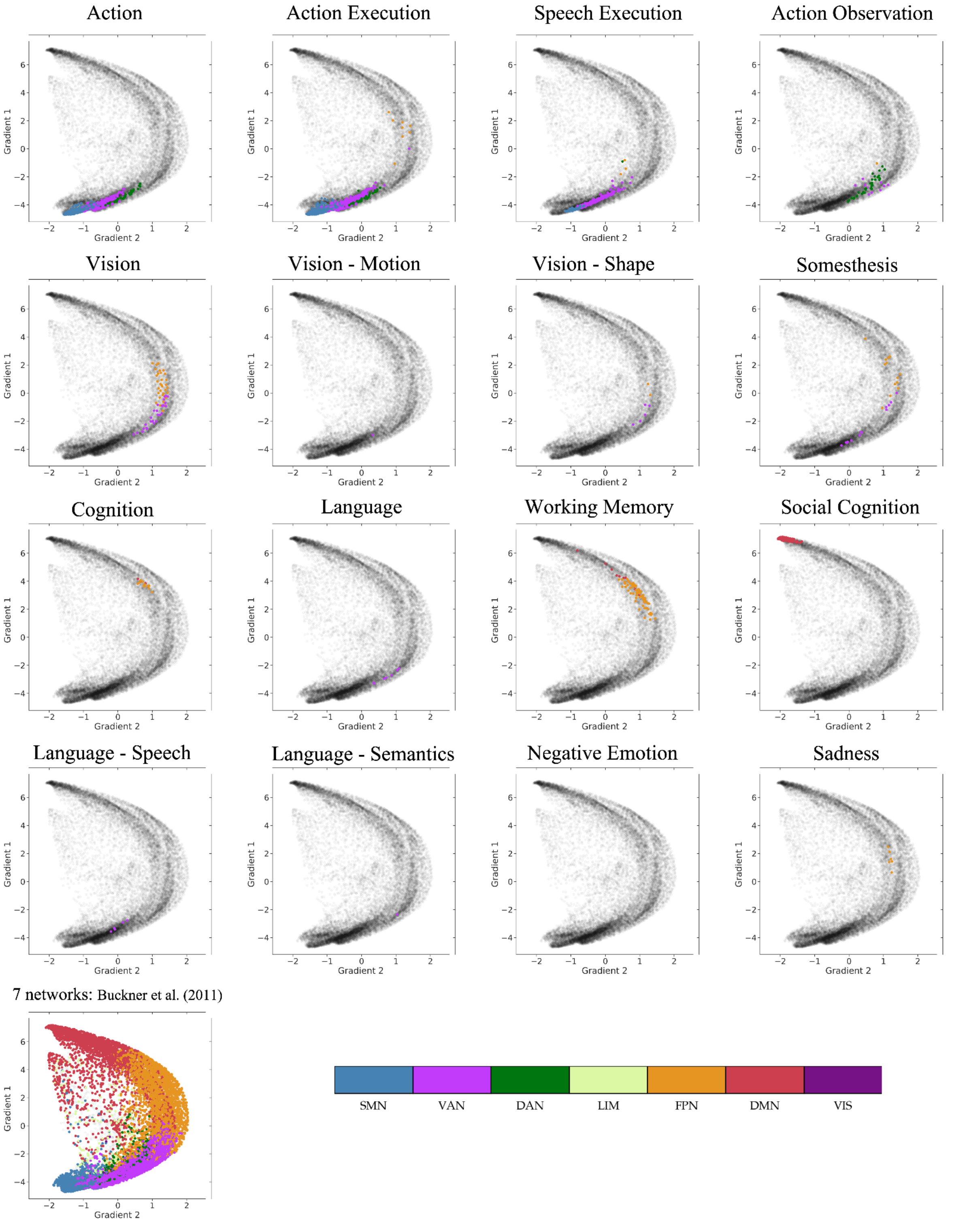


**Supplementary Figure 19: Cerebellum-specific ALE map loadings onto primary and secondary cerebellar resting-state gradients.** We assessed how thresholded cerebellum-specific ALE (C-SALE) maps loaded onto the primary and secondary cerebellar resting-state gradients (Guell et al., 2018). Dots in each subfigure represent loadings of every cerebellar voxel on the principal and secondary cerebellar gradients. Colored dots represent those voxels that reached the threshold ($p < .001$ and $k = 50$) in the respective (sub)domain. Voxels are colored by the Buckner et al. 2011 cerebellar resting-state network (Buckner et al., 2011) they map to. The bottom left panel illustrates how the 7-network parcellation maps to the primary and secondary gradients. The bottom right legend illustrates what network each color represents. Note that no thresholded C-SALE voxels mapped to the limbic (LIM) network, and the Buckner cerebellar parcellation inherently does not have any representation of the visual (VIS) network(Buckner et al., 2011). SMN = Somatomotor Network; VAN = Ventral Attention Network; DAN = Dorsal Attention Network; LIM = Limbic Network; FPN = Frontoparietal Network; DMN = Default-Mode Network; and VIS = Visual Network. All subfigures were created using the LittleBrain toolbox (Guell et al., 2019).

#### Supplementary Information

Magielse et al.

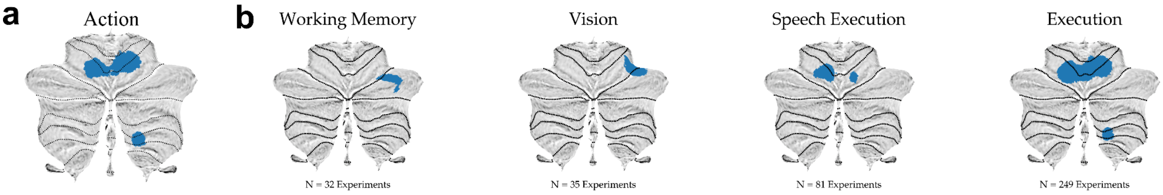


**Supplementary Figure 20: Cerebellar seeds for behavioral (sub)domains used for meta-analytic connectivity-modeling.** Meta-analytic connectivity-modeling (MACM) uses a seed (region-of-interest) to restrict experiments as input to a whole-brain-level meta-analysis. A binarized mask of converging ($p < .001$ and K = 50) cerebellum-specific ALE (C-SALE) results was used as seed region (shown in blue). For each MACM-analysis, experiments with at least one coordinate within the subdomain C-SALE mask were used as input to a whole-brain ALE analysis. Like C-SALE analyses, MACM null models were sampled from the biased probability distribution (**Supplementary Figure 2**). Subdomains are ordered by number of experiments mapping to the seed and thus used in MACM.

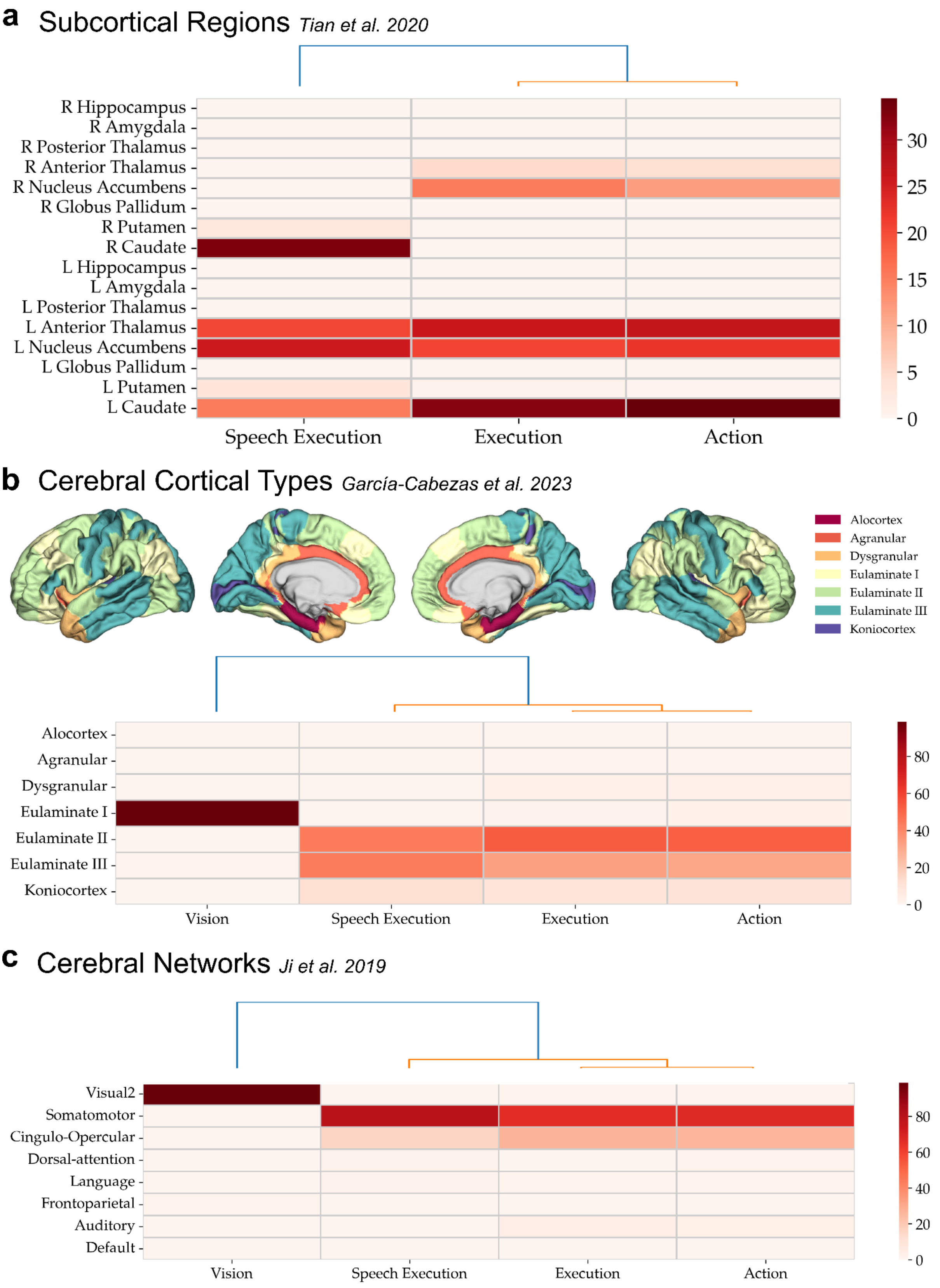


**Supplementary Figure 21: Correspondence of meta-analytic connectivity modeling maps with subcortical regions, cerebral networks and cortical types.** To contextualize brain-wide coactivation patterns in meta-analytic connectivity-modeling (MACM), we compared spatial correspondence with Tian's subcortical regions (Tian et al., 2020) (**a**), García-Cabezas' cerebral cortical types (García-Cabezas et al., 2019; Saberi et al., 2023) (**b**), and lastly the 12 Cole-Anticevic networks (Ji et al., 2019) (**c**). Specifically, we calculated the number of voxels (**a**) and vertices (**b, c**) that reached convergence for each (sub)domain's MACM map. We then calculated the proportion of voxels or vertices that fell within each parcel. Heatmaps are hierarchically clustered. Across all figures, only MACM analyses that reach convergence are reported. r = right, l = left.

### Supplementary Information

Magielse et al.

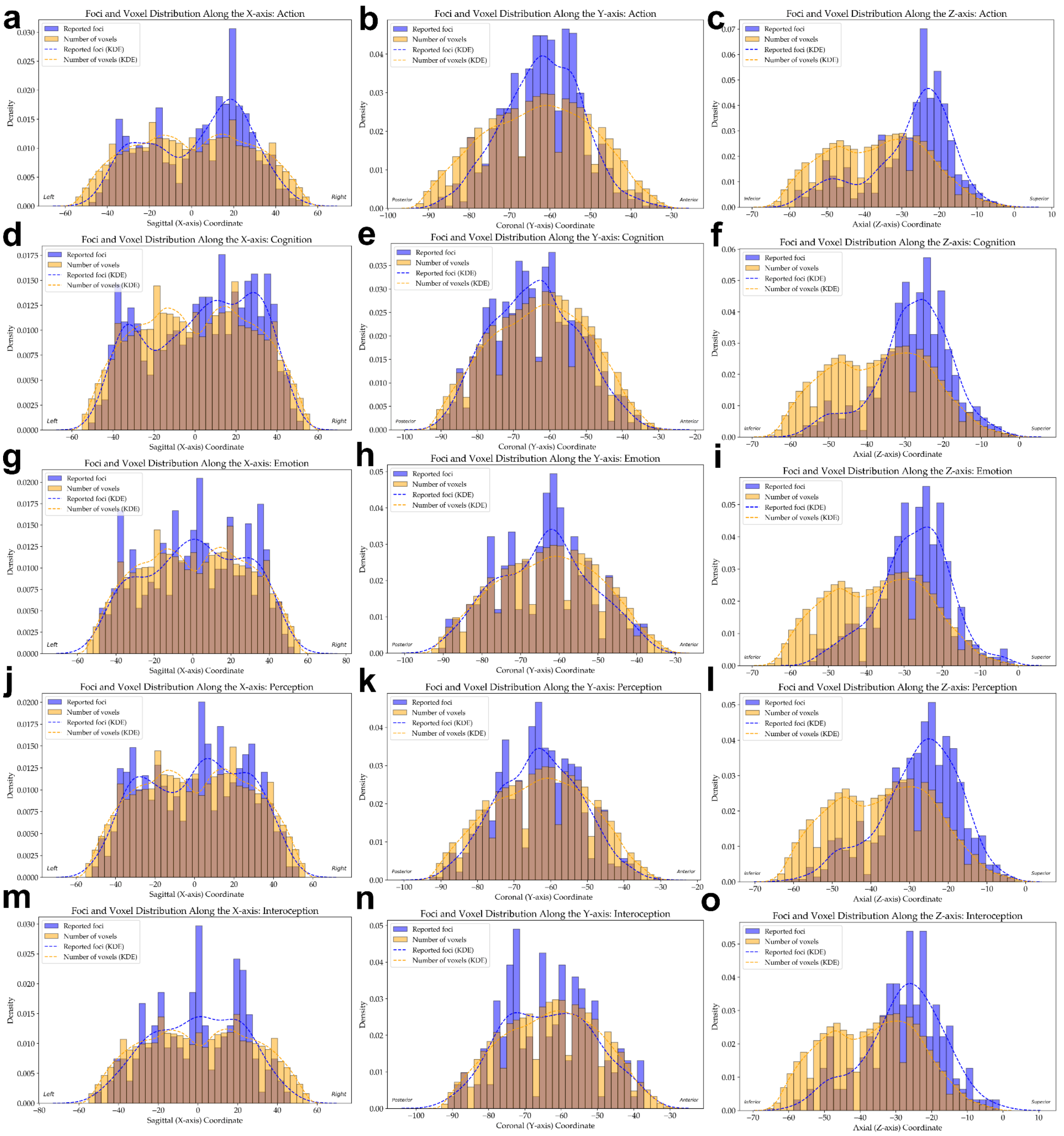


**Supplementary Figure 22: Foci distribution across brain axes per behavioral domain.** To explore the idea that a systematic and biased association may exist between neglect of the cerebellum and a behavioral domain (BD), we plotted foci distributions for each BD and brain axis separately. In each subplot, the distribution of foci is plotted relative to the distribution of cerebellar voxels, for the X- (**a, d, g, j, m**) Y- (**b, e, h, k, n**), and Z-axes (**c, f, i, l, o**) separately. Each row represents a separate BD: Action (**a, b, c**), Cognition (**d, e, f**), Emotion (**g, h, i**), Interoception (**j, k, l**), and Perception (**m, n, o**).

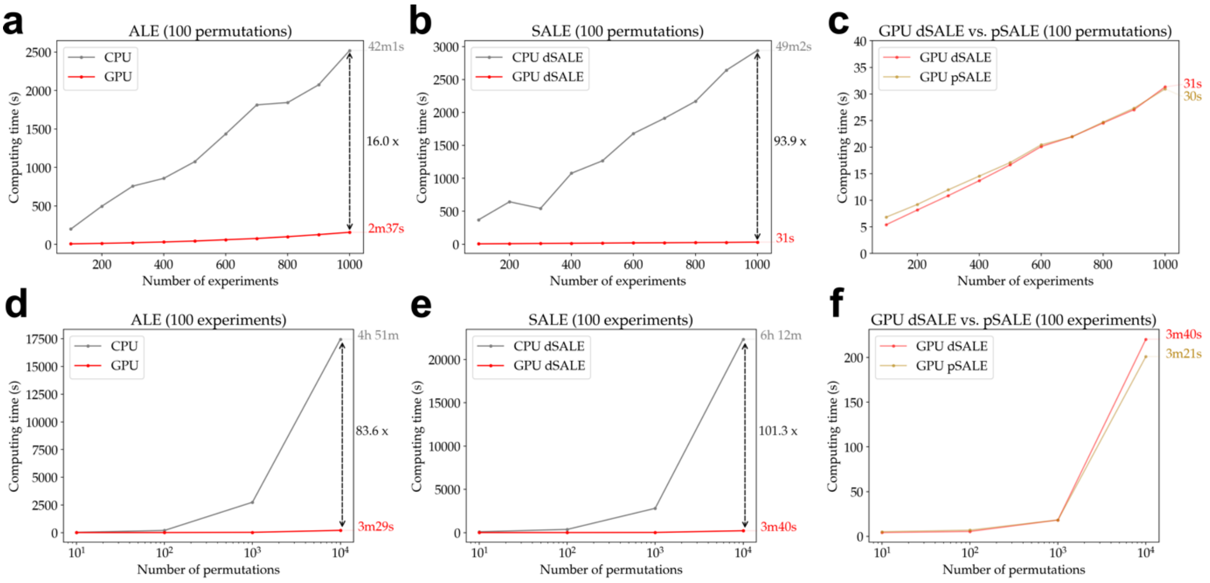


**Supplementary Figure 23: Graphical processing unit speed-up of activation likelihood activation calculations.** We show speed-ups when using graphical processing units (GPUs) relative to central procession units (CPUs), for different implementations of activation likelihood estimation (ALE): classic ALE and both deterministic (dSALE) and probabilistic versions (pSALE) of specific ALE (SALE, as in C-SALE). In essence, our GPU implementation of ALE speeds up modeled activation (MA) map calculation, the most time intensive step of ALE analysis. (**a-c**) Show performances for a fixed number of a hundred permutations, varying the number of experiments. (**d-f**) Show performances for a fixed number of a hundred experiments, varying the number of permutations. Although our implementation is not optimized for classic ALE calculations, (**a**) illustrates a sixteen-time speed-up at a thousand experiments, facilitating large-scale ALE analysis within minutes. (**b**) Shows that for dSALE, GPU was 93.9 times faster, reducing the calculation time to half a minute for a thousand experiments. Note that we compared CPU and GPU performance for dSALE, as NiMARE does not have a (CPU) pSALE implementation. As we used a cerebellum-specific version of pSALE (C-SALE), not dSALE, we show that pSALE and dSALE computing times are similar in (**c**), with pSALE being slightly faster. In (**d**), we show that GPU computing times remain low, even with many permutations. At 10,000 permutations, common for ALE, the GPU implementation is 83.6 times faster than CPU, despite not being optimized for this analysis. (**e**) Shows that GPU dSALE is 101.3 times faster at 10,000 permutations. Again, (**f**) shows that dSALE and pSALE need comparable computing times across different $N_{permutations}$, with pSALE being slightly faster at 10,000 permutations. For both ALE and SALE, GPU implementations reduced computing times of typical analyses ($N_{experiments}$ = 100; $N_{permutations}$ = 10,000) from hours to minutes. This speed-up should increase further as $N_{experiments}$ (and/ or $N_{permutations}$) increase(s). The specific machines used were an AMD EPYC 7601 (using a single core) (CPU) and a Nvidia Tesla P100 (GPU).

#### Supplementary Information

Magielse et al.
